# *Trans-*acting mutations reveal non-nuclear modulators of both intrinsic and extrinsic gene expression noise in a eukaryote

**DOI:** 10.1101/2025.11.22.689898

**Authors:** Nena Martin, Julie Kleine-Schultjann, Elena Paris, Quentin Boussau, Agnès Dumont, Hélène Duplus-Bottin, Margaux Prieux, Adelme Bazin, Laurent Modolo, Gaël Yvert, Patricia J. Wittkopp, Fabien Duveau

## Abstract

Gene expression regulation is a stochastic process that can be modified by mutations not only in a deterministic manner but also in a probabilistic way, for example by changing the extent of cell-to-cell variability in gene expression (also called “gene expression noise”). Although systematic studies have successfully analyzed the properties of *cis*-acting mutations that modulate expression noise at their own locus, less is known about the type of genetic changes that alter expression noise in *trans* (at loci distant from the mutation). Here, we applied genetic mapping on yeast strains (*Saccharomyces cerevisiae*) generated by random mutagenesis and identified three mutations (in *chs1*, *yme2* and *msh1* genes) that changed the expression noise of a reporter gene in the nuclear genome regulated by the yeast *TDH3* promoter. These mutations affected either extrinsic noise (variability due to cell-specific factors), intrinsic noise (inherent variability within each cell) or both, and their effects were found to not be specific to the *TDH3* promoter. Surprisingly, all three mutations targeted proteins located outside of the nucleus: Yme2 and Msh1 being involved in the maintenance of mitochondrial genome integrity, and Chs1 being necessary for the repair of cell-wall defects in freshly-born daughter cells. Our results reveal that mitochondrial state can modulate the extent of intrinsic expression noise of eukaryotic nuclear genes.

## INTRODUCTION

Gene expression levels vary among cells even in isogenic populations originating from a common cell type and maintained in a constant environment. This variability - commonly named *gene expression noise -* results from random fluctuations that inevitably occur along molecular and cellular processes impacting gene expression (reviewed in Pal & Dhar, 2024; Sanchez & Golding, 2013; Verhagen & Hansen, 2025). Expression noise can be modified by genetic changes, leading to differences in cell-to-cell variability among genotypes that can be either detrimental, neutral or beneficial depending on the gene and on the environmental condition; expression noise is therefore a substrate for evolution (M. Richard and Yvert, 2014). Previous studies showed that genetic variation of expression noise could impact population fitness (Duveau et al., 2018; Schmiedel et al., 2019) and therefore be targeted by selection (Metzger et al., 2015). Importantly, a global relationship is known to exist between mean expression and noise, whether among different genes (Newman et al., 2006; Taniguchi et al., 2010), among different genetic variants of a promoter (Carey et al., 2013; Hornung et al., 2012) or among environmental conditions (Bar-Even et al., 2006). Although the confounding effect of mean expression has complicated the study of the genetic sources and functional consequences of variation in expression noise, statistical and experimental approaches have been specifically designed to uncouple variation in these two metrics.

Seminal studies revealed that total noise in gene expression had two components, *extrinsic* and *intrinsic*, each attributable to distinct sources (Elowitz et al., 2002; Raser & O’Shea, 2004). Technically, extrinsic and intrinsic noise can be measured in cells where transcriptional activities of two copies of the same promoter are distinguishable (for instance using two fluorescent reporter genes of different colors; but see Stewart-Ornstein et al. (2012) for an alternative approach). In this setup, extrinsic noise represents expression variation among isogenic cells (or over time in a cell) that is correlated between the two copies of the promoter, while intrinsic noise represents independent variation of expression between the two copies. Biologically, e*xtrinsic noise* results from sources generating heterogeneity in global state among cells, with consequences on how individual cells express the gene. In contrast, *intrinsic noise* corresponds to fluctuations that remain in cells after accounting for their state due to the inherent stochasticity of regulatory processes. Experimental and theoretical work have previously studied the properties of these two components and how specific steps of gene regulation contribute to them. The predominant source of intrinsic noise was shown to be transcription, which occurs stochastically in bursts (Blake et al., 2006; Harper et al., 2011; Pedraza & Paulsson, 2008; Quarton et al., 2020; Suter et al., 2011). The extent to which transcription contributes to intrinsic noise depends on chromatin remodeling (Raser & O’Shea, 2004), on nucleosome occupancy (Tirosh & Barkai, 2008), on the presence of TATA boxes in regulatory sequences (Blake et al., 2006; Choi & Kim, 2009; Hornung et al., 2012; Sun & Zhang, 2020) and on the number and type of transcription factors binding to the promoter (Parab et al., 2022; Sharon et al., 2014). Extrinsic noise is known to result from various cell-to-cell differences such as their pool of RNA polymerase (Yang et al., 2014), micro-environment (Snijder et al., 2009), age (P. Liu et al., 2017; Martinez-Jimenez et al., 2017), cell cycle stage (Sukys & Grima, 2025; Zopf et al., 2013), global transcription rate (Sherman et al., 2015), mitochondrial content (Guantes et al., 2015) and proliferation rate (Keren et al., 2015). Extrinsic noise was also shown to be reduced for genes regulated by miRNAs (Siciliano et al., 2013).

All factors modulating gene expression noise are possible targets for mutations and therefore provide a potential substrate for the natural evolution of cell-to-cell variability. As for any trait, two driving forces may influence the evolution of gene expression noise: the effects of random mutations on noise and the effects of selection on the fate of these mutations in natural populations. Regarding the action of selection on expression noise, several important observations have been reported. First, gene expression noise can be evolved experimentally through artificial selection (Wolf et al., 2015; You et al., 2019). Second, several studies provided evidence of negative selection acting against elevated expression noise in natural populations, suggesting that the detrimental effect of elevated noise could be counter-selected separately from any effect of a genetic variant on mean expression (Barroso et al., 2018; Metzger et al., 2015; Puzović et al., 2023; Schor et al., 2017; Vlková & Silander, 2022) . Reciprocally, other studies showed that gene expression noise could be positively selected in certain cases, particularly in fluctuating environments (Acar et al., 2008; Eldar & Elowitz, 2010; Rocabert et al., 2020; Schmutzer & Wagner, 2020). When cells are maintained in a steady environment, increasing expression noise of a gene can be either beneficial or detrimental depending on the mean expression level and on the shape of the relationship that exists between the expression level of the gene and the rate of cell proliferation (Duveau et al., 2018; Schmiedel et al., 2019). In yeast, transcriptome-wide analyses revealed a particularly high level of expression noise for plasma-membrane transporters, which play a critical role in cellular responses to environmental changes (Zhang et al., 2009). Experimental studies in bacteria (Wolf et al., 2015) and yeast (Bódi et al., 2017) showed that elevated expression noise could facilitate the evolution of gene regulation. By generating fitness variation among individuals, expression noise can also potentially reduce the efficacy of selection and therefore affect phenotypic evolution (Mineta et al., 2015; Wang & Zhang, 2011). In multicellular organisms, expression noise was associated with cell fate decision (Balázsi et al., 2011; Coomer et al., 2022; Ohnishi et al., 2014; Pal & Dhar, 2024; A. Richard et al., 2016; Wernet et al., 2006) and in the resistance of cancer cells to chemotherapy (Shaffer et al., 2017). Finding what genetic variants and molecular mechanisms can possibly modulate expression noise is therefore essential for understanding many fundamental biological processes.

In this regard, important discoveries were made about the type of genetic changes that can tune gene expression noise and their mechanisms of action. First, genetic variants segregating in natural populations were found to generate different levels of gene expression noise among individuals. This was first shown in yeast (Ansel et al., 2008; Fehrmann et al., 2013; J. Liu et al., 2015; Metzger et al., 2015) and later in bacteria (Vlková & Silander, 2022), flies (Huang et al., 2015) and human cells (Lu et al., 2016; Morgan et al., 2020; Triqueneaux et al., 2020). Second, as previously demonstrated for changes in mean expression (Wittkopp et al., 2004), a mutation may modify the expression noise of a focal gene in two different ways (Raser & O’Shea, 2004): the mutation may act in *cis* (if the mutation is at the locus of the regulated gene) or in *trans* (if the mutation is at a locus physically distant from the regulated gene). Importantly, a single mutation may act in *cis* on the expression of a nearby gene and in *trans* on the expression of another more distant gene, for example if the first gene is a regulator of the second gene. *Cis* effects of mutations are better accessible for investigations than *trans* effects because of the close proximity in the genome between *cis*-acting mutations and the gene with altered expression. Several studies were therefore conducted on specific genes to understand how *cis*-located mutations modify the expression noise of these genes. A seminal study focusing on the *GAL1* promoter in yeast showed how mutations in the TATA box could modulate noise by altering the dynamics of transcriptional bursts (Blake et al., 2006). In a later study, a more comprehensive random mutagenesis of 22 yeast promoters revealed a mathematical relationship between mean expression and expression noise, showing that mutations primarily altered the frequency of transcriptional bursts while burst size differed among promoters, producing promoter-specific relationships between mean and noise (Hornung et al., 2012). Using an ingenious high-throughput approach to quantify mean expression and expression noise from thousands of designed synthetic promoters, another yeast study showed that both the number of functional transcription-factor binding sites and the frequency of poly(dT/dA) motifs - which favor nucleosome positioning – were key determinants of expression noise (Sharon et al., 2014). A fourth study compared the effects of 236 artificial mutations and 45 natural polymorphisms in the promoter of the yeast gene *TDH3*, providing evidence of negative selection against *cis*-acting mutations that increased noise of *TDH3* promoter activity (Metzger et al., 2015).

Although there are only few known examples of mutations affecting gene expression noise in *trans*, their effects on noise may differ from those of *cis*-acting mutations. In particular, the relative contribution of *cis-* and *trans*-acting mutations on intrinsic and extrinsic noise may not be the same. Indeed, theoretical models predict that *cis*-acting mutations in a regulatory sequence may modulate intrinsic noise by altering the frequency and magnitude of transcriptional bursts (Pedraza & Paulsson, 2008; Zenklusen et al., 2008). It is more difficult to imagine a scenario where a *cis*-acting mutation would modulate extrinsic noise. Mutations affecting expression noise in *trans*, however, could modulate either intrinsic or extrinsic noise. For instance, while mutations in a transcription factor may affect intrinsic noise of its target genes, mutations in a cell cycle regulator may impact extrinsic noise of other genes by generating heterogeneity of states among cells. Given these differences, it is necessary to further characterize what type of mutations can modulate gene expression noise in *trans* and what their mechanisms of action are to understand how expression noise is controlled genetically and evolves. One way to do so is by introducing target mutations in molecular complexes participating to gene expression and determine how they modify cell-to-cell variability in protein levels, as was done for mutations targeting chromatin remodeling (Raser & O’Shea, 2004) or transcription elongation (Ansel et al., 2008; Trinh et al., 2023). Such targeted approach, however, is driven by existing knowledge on gene regulation and does not interrogate the possible roles of unexpected players. An alternative approach is to identify *trans*-regulators segregating in natural populations using adapted GWAS or QTL mapping methods (Chuffart et al., 2016; Fehrmann et al., 2013; Lu et al., 2016; Morgan et al., 2020; Sarkar et al., 2019; Yvert, 2014). Although this approach is well-suited to characterize genetic variants that have been filtered by selection, it does not capture all possible genetic sources of variation in expression noise (Hill et al., 2021).

Here we used a random mutagenesis screen followed by genetic mapping and functional assays to identify mutations that changed in *trans* the expression noise of a focal gene in yeast. We took advantage of a collection of 1241 mutant strains obtained by chemical mutagenesis of a parental strain expressing a fluorescent reporter gene controlled by the *TDH3* promoter (*P_TDH3_-YFP*). In its native context, the *TDH3* promoter drives strong constitutive expression of a glyceraldehyde-3-phosphate dehydrogenase involved in glycolysis and gluconeogenesis (Partow et al., 2010). Using flow cytometry, we identified five mutants from the collection that showed altered *P_TDH3_-YFP* expression noise relative to the parental strain without significant change in mean expression. Via genetic mapping, we identified six candidate causal mutations in these mutants and we validated the effects of three of them. These mutations were found to increase expression noise driven not only by the *TDH3* promoter, but also by two other yeast promoters. We then characterized the contribution of these mutations to extrinsic and intrinsic noise using two-color reporters. The three mutations altered the coding sequences of *YME2*, *MSH1* and *CHS1* genes, none of which were previously associated with the control of noise or mean gene expression. In particular, our results suggest a potentially central and unexpected role of mitochondrial genome integrity in the modulation of intrinsic expression noise of nuclear genes.

## RESULTS

### Mapping mutations with *trans*-acting effects on expression noise

We previously identified mutations that affected the median expression level of a focal gene (*P_TDH3_-YFP*, where expression of a *YFP* reporter gene is controlled by the *TDH3* promoter) from a collection of haploid yeast strains obtained by ethyl methanesulfonate (EMS) mutagenesis of a reference strain (Duveau et al., 2021; Metzger et al., 2016). Here, we took advantage of this collection of 1241 strains to identify and characterize mutations that altered expression noise of the focal gene in *trans* (*i.e.* mutations outside of the *TDH3* promoter and causing increased or decreased cell-to-cell variability of fluorescence level). By measuring the fluorescence level of thousands of cells from each strain using flow cytometry in two consecutive screens (**Figure 1a**; **Figure 1 – figure supplement 1**), we identified three mutants with increased *P_TDH3_-YFP* expression noise and two mutants with decreased *P_TDH3_-YFP* expression noise relative to the reference strain (**Figure 1a**; **Table 1**). These five mutants showed reproducible change in expression noise, but no significant change in median expression level. The procedure we followed to choose the five strains is further described in **Supplementary File 5**.

**Figure 1.**
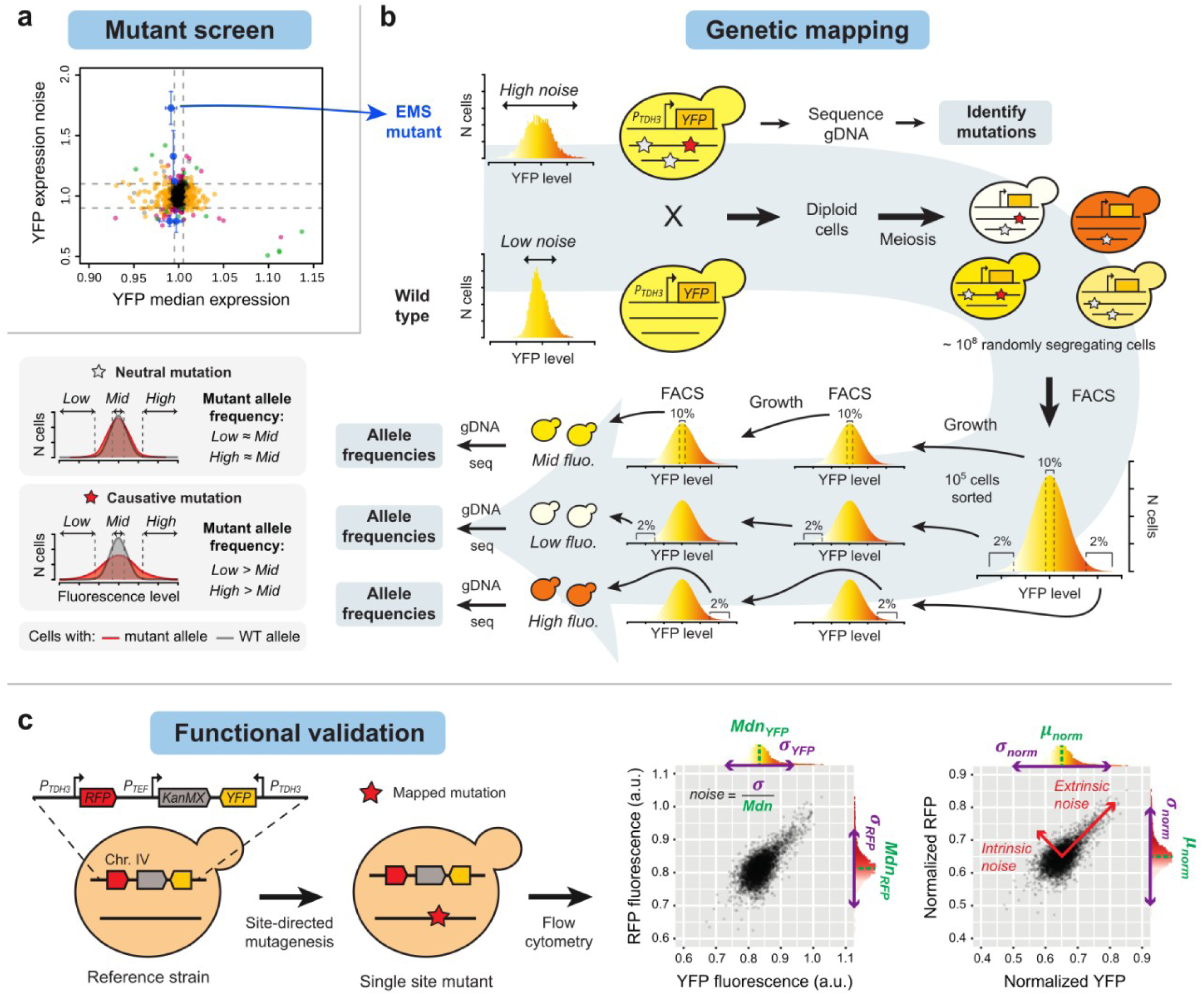
Identification of mutations affecting cell-to-cell variability of expression of a target gene. **(a)** First mutagenesis screen performed to identify 5 strains (blue dots) with altered expression noise and unchanged median expression of the *P_TDH3_-YFP* reporter. Each dot represents the mean value of median expression or expression noise among four replicate samples relative to the reference strain. Expression noise was computed as the standard deviation divided by the median normalized fluorescence among cells, as described in Methods. Error bars show 95% confidence intervals. Green and blue dots: strains with significant change of expression noise both in screen 1 and screen 2. Magenta dots: strains with significant change of expression noise only in screen 1. Orange dots: other strains included in screen 2 (with altered median expression in screen 1). Gray dots: strains with non-significant change of median expression or expression noise in screen 1 (not included in screen 2). See Methods for details on how these categories were determined. **(b)** Scheme of the bulked segregant analysis and whole-genome sequencing mapping strategy (BSA-seq) used to determine which mutations were statistically associated with expression noise. **(c)** Strategy used to confirm the effects of mapped mutations on expression noise. Mapped mutation were individually introduced in the genome of a strain expressing two different reporter genes (*RFP* and *YFP*), each under control of a copy of the *TDH3* promoter. These single-site mutants allowed us to quantify the effects of each mutation not only on expression noise of the two fluorescent reporters, but also on intrinsic and extrinsic noise (from the covariation of fluorescence of the two reporters, see Methods).

**Figure 1 – figure supplement 1.**
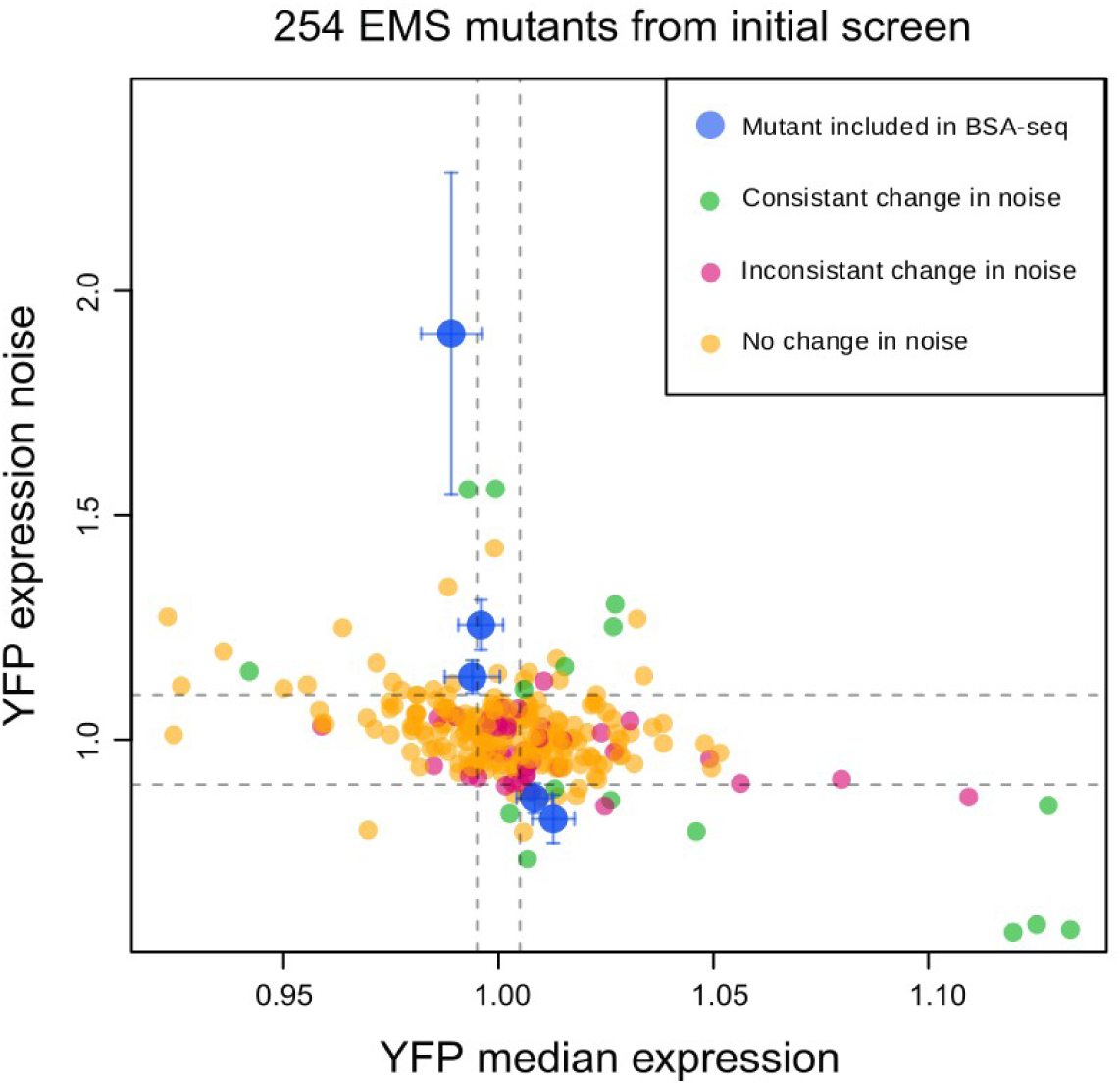
Second fluorescence screen used to select mutant strains with reproducible changes of expression noise for genetic mapping. Median expression and expression noise of YFP were quantified in 254 mutant strains that showed significant changes of YFP expression (median and/or noise) relative to the reference strain in the initial screen ( Figure 1a). Each dot represents the mean value of median expression or expression noise among four replicate samples relative to the reference strain. Expression noise was computed as the standard deviation divided by the median normalized fluorescence among cells, as described in Methods. Error bars show 95% confidence intervals. The five strains selected for genetic mapping are shown in blue. Other colors indicate whether a strain showed significant change of expression noise in both screen 1 and screen 2 or not (as indicated in top right inset).

**Figure 1 – figure supplement 2.**
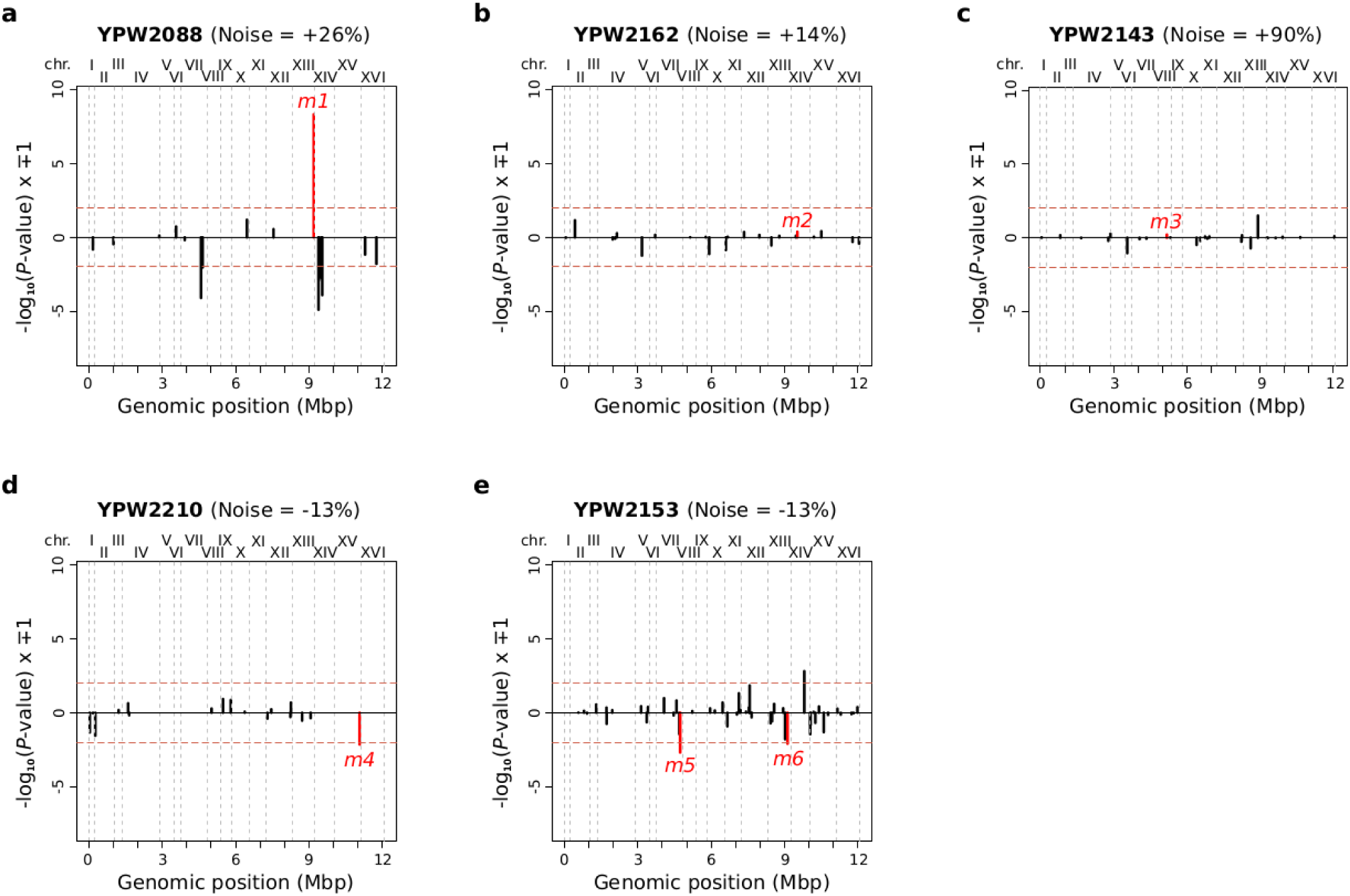
Statistical comparisons of allele frequencies between different fluorescent bulks at each mutation site in the genome of segregants from five crosses. For each mutation site (each vertical bar) in **(a)**, **(b)**, **(d)**, and **(e)**, the *P*-value was computed using a *G*- test (as described in Methods) to compare read counts with mutant and wild-type alleles between i) low and high fluorescence bulks combined and ii) mid fluorescence bulk. For the cross involving mutant YPW2143 **(c)**, we only compared high and mid fluorescence bulks because a single haplotype was nearly fixed in the low fluorescence bulk. *P*-values were adjusted using the Benjamini-Hochberg procedure to control the false-discovery rate. To visualize directions of effects, we represented −log_10_ ( *P*) values with a positive sign if a mutation was found at higher frequency in the low and high bulks than in the medium bulk (*i.e.* mutation associated with increased expression noise) and a negative sign if a mutation was found at lower frequency in the low and high bulks than in the medium bulk (*i.e.* mutation associated with decreased expression noise). Vertical dotted lines show boundaries between chromosomes. Horizontal red dotted lines show *P*-value thresholds of 0.01 for mutations with positive and negative effects. The six mutations for which we attempted to validate the effect on expression noise are represented in red.

**Figure 1 – figure supplement 3.**
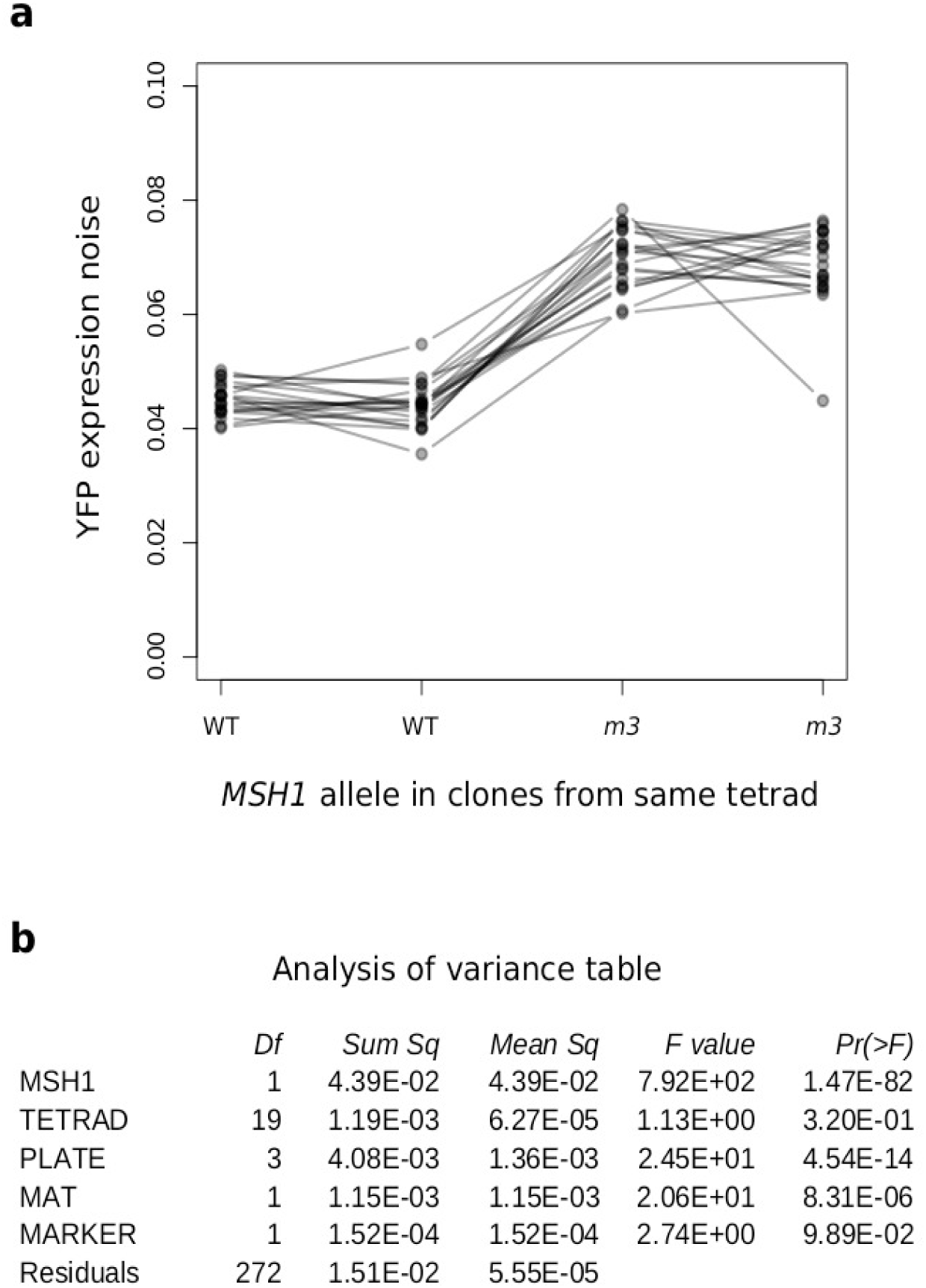
Linkage analysis showing a strong association between expression noise and segregation of *m3* mutation (*msh1(G1262a)*) in F2 hybrids. *P_TDH3_-YFP* expression noise was quantified by flow cytometry and *msh1* genotype was determined by Sanger sequencing in 80 F2 clones isolated from 20 tetrads. F1 diploid cells were obtained by crossing mutant YPW2143 with the mapping strain YPW1240. **(a)** Each dot shows the mean expression noise measured from 6094 to 8799 cells in 3 or 4 replicate populations for each F2 clone. Lines connect the four clones issued from the same tetrad. **(b)** Results of an ANOVA used to determine the statistical significance of the additive effects of the following explanatory variables on expression noise: i) *MSH1* genotype, ii) the tetrad of origin of each clone, iii) the culture plate (each replicate was grown in a different plate), iv) the mating type and v) the drug resistance marker segregating in F2 clones. The adjusted *R*-squared of the linear model used in the ANOVA is 0.7489 and the associated *F*-statistic is 36.43 (*P*-value = 3.30 x 10^-72^).

**Table 1.** Properties of the reference strain and of the 5 mutants with altered *P_TDH3_-YFP* expression noise included in the genetic mapping experiments.

| Strain name | Type | Number of mutations | YFP median expr. (relative) Assay 1 | YFP noise (relative) Assay 1 | YFP noise (absolute) Assay 1 | YFP median expr. (relative) Assay 2 | YFP noise (relative) Assay 2 | YFP noise (absolute) Assay 2 |
| --- | --- | --- | --- | --- | --- | --- | --- | --- |
| YPW1139 | Reference | 0 | 1 | 1 | 0.0152 | 1 | 1 | 0.0165 |
| YPW2088 | EMS mutant | 16 | 0.994 | 1.329 | 0.0203 | 0.996 | 1.255 | 0.0208 |
| YPW2162 | EMS mutant | 22 | 0.995 | 1.123 | 0.0172 | 0.994 | 1.140 | 0.0188 |
| YPW2143 | EMS mutant | 26 | 0.991 | 1.728 | 0.0261 | 0.989 | 1.904 | 0.0309 |
| YPW2210 | EMS mutant | 17 | 0.990 | 0.792 | 0.0121 | 1.008 | 0.870 | 0.0146 |
| YPW2153 | EMS mutant | 47 | 0.997 | 0.791 | 0.0120 | 1.013 | 0.824 | 0.0134 |

Using whole-genome sequencing with an average coverage of 100x, we found between 16 and 47 mutations in the genomes of these five mutants (**Table 1**), which was consistent with the range (14 to 41) of mutations previously identified in other mutants from the same collection (Duveau et al., 2021). All mutations were fixed and we did not observe any evidence of genetic heterogeneity or previously unknown structural variant in each mutant strain (**Supplementary File 4**). Given that the vast majority of mutants in the collection showed no significant change in expression noise, we expected that only one or a few mutations were responsible for the altered expression noise in each of the five selected mutants. We therefore applied a genetic mapping strategy specifically designed to identify these presumably-rare mutations (**Figure 1b**). We used a bulk segregant analysis approach where each haploid mutant was crossed to a wild-type strain also expressing *P_TDH3_-YFP* and random segregants obtained from the meiosis of resulting diploid cells were sorted by FACS based on their fluorescence levels (see Methods). Among all segregants, the subset of cells that inherited a mutation responsible for a change in expression noise of the fluorescent reporter (a causative mutation) should show either a broader or a narrower distribution of fluorescence levels than the subset of cells inheriting the wild-type allele. For this reason, we sorted random segregants in three bulks that captured distinct ranges of fluorescence levels: one bulk corresponding to the 2% of cells with lowest fluorescence (low bulk), one bulk corresponding to the 2% of cells with highest fluorescence (high bulk) and one bulk corresponding to the 10% of cells with fluorescence closest to the median fluorescence of all segregants (mid bulk). We reasoned that a causative mutation should be found at different frequencies in the mid bulk than in the low and high bulks. To further increase this difference, we repeated the fluorescence selection twice for each bulk with a few generations of growth between each selection step (**Figure 1b**). For each cross, frequencies of all segregating mutations were then quantified by deep sequencing of genomic DNA extracted from each bulk. Likelihood-ratio tests (*G*-tests) were used to determine if the frequency of each mutation was significantly different between the mid bulk and the combined low and high bulks (**Figure 1 – figure supplement 2**), in which case the mutation was considered causative (or genetically linked to a causative mutation). Using this approach, we found one candidate causative mutations in two out of five mutants analyzed and two candidate mutations in another mutant (**Table 2**). For a fourth mutant (YPW2162), although no mutation was significantly associated with expression noise, a single mutation was found at very low frequency in all fluorescence bulks (*m2* in **Table 2**). This indicated that the mutation was strongly counter-selected during the procedure used to establish fluorescent bulks, which greatly reduced the statistical power to detect potential association with expression noise. Therefore, we could not exclude that mutation *m2* was causative and we decided to further investigate its impact on expression noise in the functional assays described below. For the last mutant (YPW2143), we noticed that the frequency of all 26 mutations was either near zero or near one in the low bulk of segregants. This unexpected observation suggested that a single haplotype was nearly fixed in the low bulk, either due to a strong selective advantage of this haplotype or more likely due to a strong bottleneck in population size when this bulk was established. For this reason, we only compared allele frequencies between mid and high bulks, but no mutation was found at a significantly different frequency between these two bulks (**Figure 1 – figure supplement 2c**). Therefore, we used another strategy based on linkage analysis and tetrad dissection to determine which mutation in mutant YPW2143 was associated with variation of expression noise. The candidate mutation we tested first was mutation *m3* (**Table 2**) because it was found at a slightly higher frequency in the high bulk and this mutation appeared to be counter-selected in both bulks, which reduced the statistical power to detect different frequencies of the mutation between bulks. After meiosis of diploid cells from the cross between YPW2143 and the mapping wild-type strain, we measured *P_TDH3_-YFP* expression noise in 80 segregant clones issued from 20 dissected tetrads. We observed a very strong association between expression noise and the presence or absence of mutation *m3* in each clone (**Figure 1 – figure supplement 3**; ANOVA, *F* value = 792.31, *P* = 1.47 x 10^-82^), suggesting this mutation was causative or tightly linked to a causative mutation.

**Table 2.**
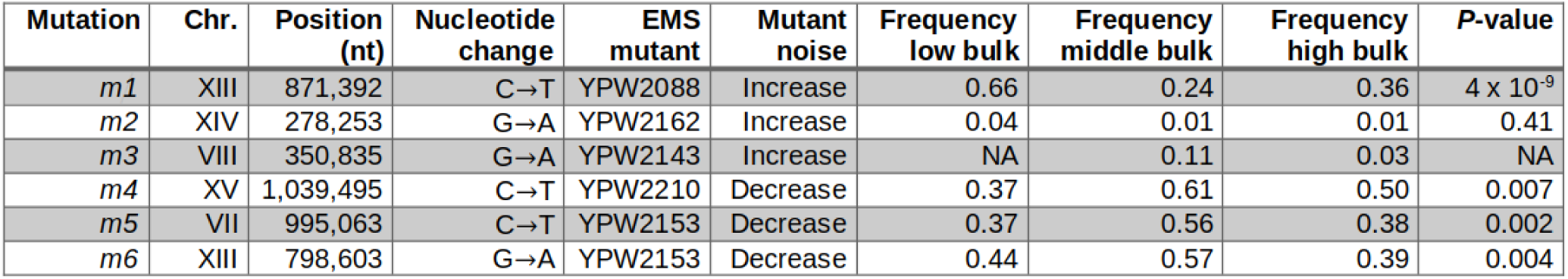
Candidate mutations showing statistical association with variation of *P_TDH3_-YFP* expression noise or low frequency in all fluorescence bulks.

| Mutation | Chr. | Position (nt) | Nucleotide change | EMS mutant | Mutant noise | Frequency low bulk | Frequency middle bulk | Frequency high bulk | P-value |
| --- | --- | --- | --- | --- | --- | --- | --- | --- | --- |
| *m1* | XIII | 871,392 | C→T | YPW2088 | Increase | 0.66 | 0.24 | 0.36 | $4 \times 10^{-9}$ |
| *m2* | XIV | 278,253 | G→A | YPW2162 | Increase | 0.04 | 0.01 | 0.01 | 0.41 |
| *m3* | VIII | 350,835 | G→A | YPW2143 | Increase | NA | 0.11 | 0.03 | NA |
| *m4* | XV | 1,039,495 | C→T | YPW2210 | Decrease | 0.37 | 0.61 | 0.50 | 0.007 |
| *m5* | VII | 995,063 | C→T | YPW2153 | Decrease | 0.37 | 0.56 | 0.38 | 0.002 |
| *m6* | XIII | 798,603 | G→A | YPW2153 | Decrease | 0.44 | 0.57 | 0.39 | 0.004 |

### Functional validation of effects of candidate mutations on expression noise

Statistical associations described above may be explained by a causal effect of candidate mutations on *P_TDH3_-YFP* expression noise, but they could also result from genetic linkage with a causative mutation or from unsuspected selection for another trait. To demonstrate the causal impact of each candidate mutation (six mutations in **Table 2**) on expression noise, we used site-directed mutagenesis to introduce each mutation in a common genetic background (reference strain) that does not harbor any other mutations found in EMS mutants and that contains two distinct reporter genes inserted in opposite orientations at the *ho* locus (**Figure 1c**). As further described in the next sections, the use of two reporter genes (*YFP* and *RFP* coding sequences transcribed under control of distinct copies of the *TDH3* promoter) allowed us to distinguish two sources of expression noise. We then used flow cytometry to quantify median expression levels and expression noise of all site-directed mutants, the reference strain and EMS mutants from which candidate mutations were identified (note that EMS mutants do not express the *RFP* reporter gene). In most experiments reported below, we included two independent clones for each genotype (*i.e.* two strains established from two colonies from the same transformation) to detect cases where a secondary mutation not shared by all transformed cells would contribute to phenotypic variation (*i.e.* cases where the two independent clones would show statistically different phenotypes).

For three candidate mutations (*m1*, *m2* and *m3*), we observed a significant difference of expression noise between site-directed mutants and the reference strain (*t*-test, adjusted *P* < 0.01), both for YFP (**Figure 2a-c**) and for RFP (**Figure 2 – Figure supplement 2g-i**). In all three cases, the mutation caused an increase in expression noise relative to the allele of the reference strain. In addition, no significant difference of YFP expression noise was observed between each site-directed mutant and the EMS mutant carrying the same mutation (**Figure 2a-c**), indicating that the three candidate mutations were responsible for most, if not all, of the variation of expression noise observed in EMS mutants from which they were identified. We also showed that the effect of the three mutations did not change when using two different metrics of expression noise (**Figure 2 – figure supplement 4**). As expected, we observed no significant difference of median expression between the reference strain and site-directed or EMS mutants carrying mutations *m1* and *m2* (*t*-test, adjusted *P* ⩾ 0.01), both for YFP (**Figure 2 – figure supplement 1a-b**) and for RFP (**Figure 2 – figure supplement 2a-b**). However, all strains carrying mutation *m3* showed a significant increase of YFP (**Figure 2 – figure supplement 1c**) and RFP (**Figure 2 – figure supplement 2c**) median expression levels relative to the reference strain (*t*-test, adjusted *P* < 0.01), which was unexpected based on the initial screen (**Figure 1a**). We further investigated the cause of this apparent inconsistency below (**Figure 6 – figure supplement 2**).

**Figure 2.**
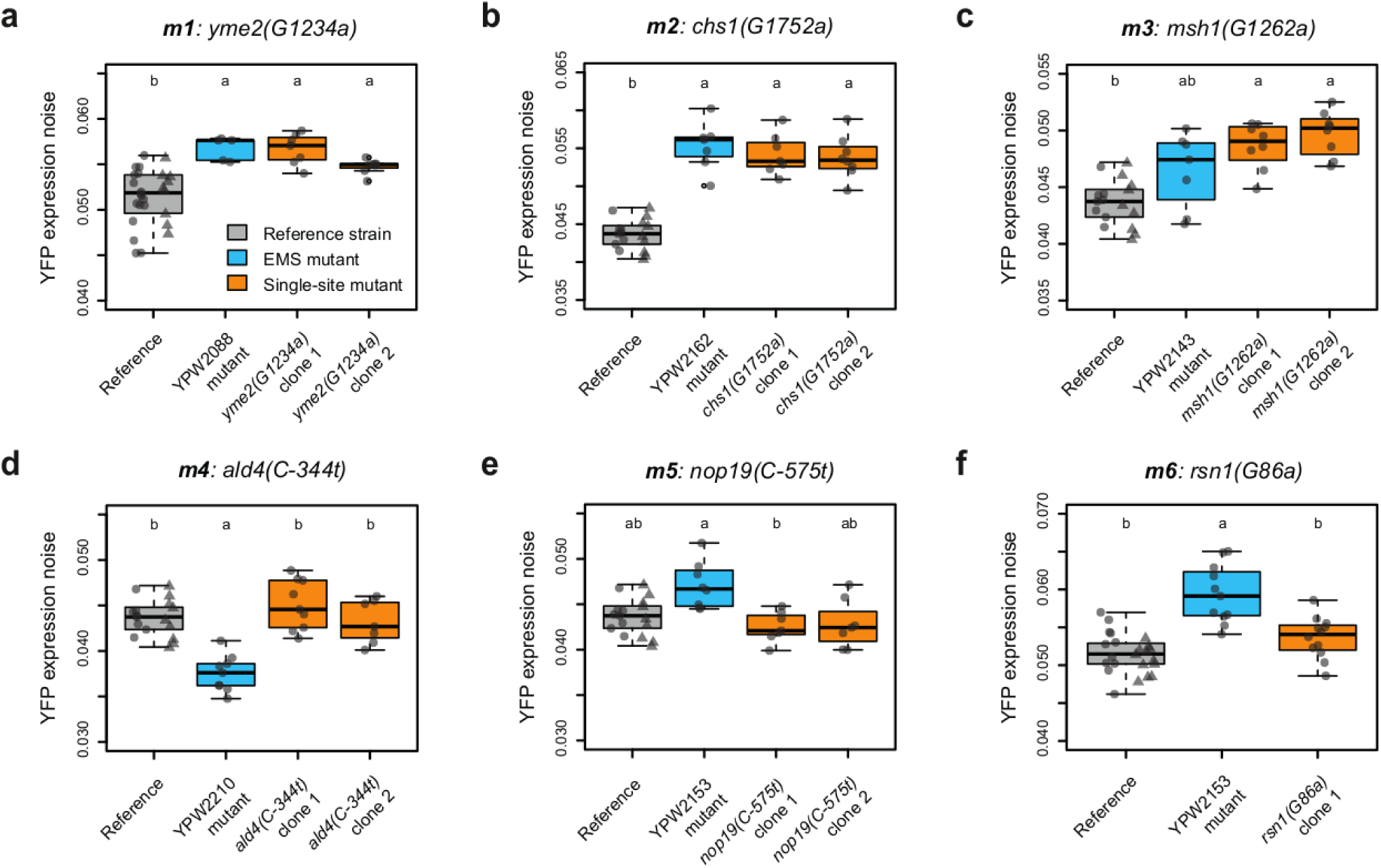
Testing the effects of six mapped mutations on YFP expression noise using single-site mutants. Thick lines, boxes and whiskers represent the median, interquartile range (IQR) and 1.5 IQR of expression noise measured among 7 to 26 samples, with colors corresponding to different genotypes as indicated in the legend of panel **(a)**. Dots represent the expression noise of individual samples measured from 2102 to 4707 cells. Expression noise was computed as the standard deviation divided by the median normalized fluorescence among cells, as described in Methods. For the reference genotype, circles and triangles represent data collected for two independent clones obtained from the same transformation experiment. For genotypes carrying single mutations, data collected for independent clones from the same transformation are shown in two orange boxes. Pairwise *t*-tests were performed to compare the mean expression noise among replicates between pairs of genotypes, with *P*-values adjusted using Benjamini-Hochberg correction. For each panel, the mean expression noise is statistically different between two genotypes (adjusted *P*-value < 0.05) if no letter is shared above the boxes of the two genotypes. Above each plot, positions of mutations are indicated relative to the start codon of the gene with positive and negative numbers if mutations are respectively downstream ( *i.e.* in coding sequence) and upstream of the start codon. Capital and lowercase letters indicate respectively reference and alternate nucleotides at each mutation site.

**Figure 2 – figure supplement 1.**
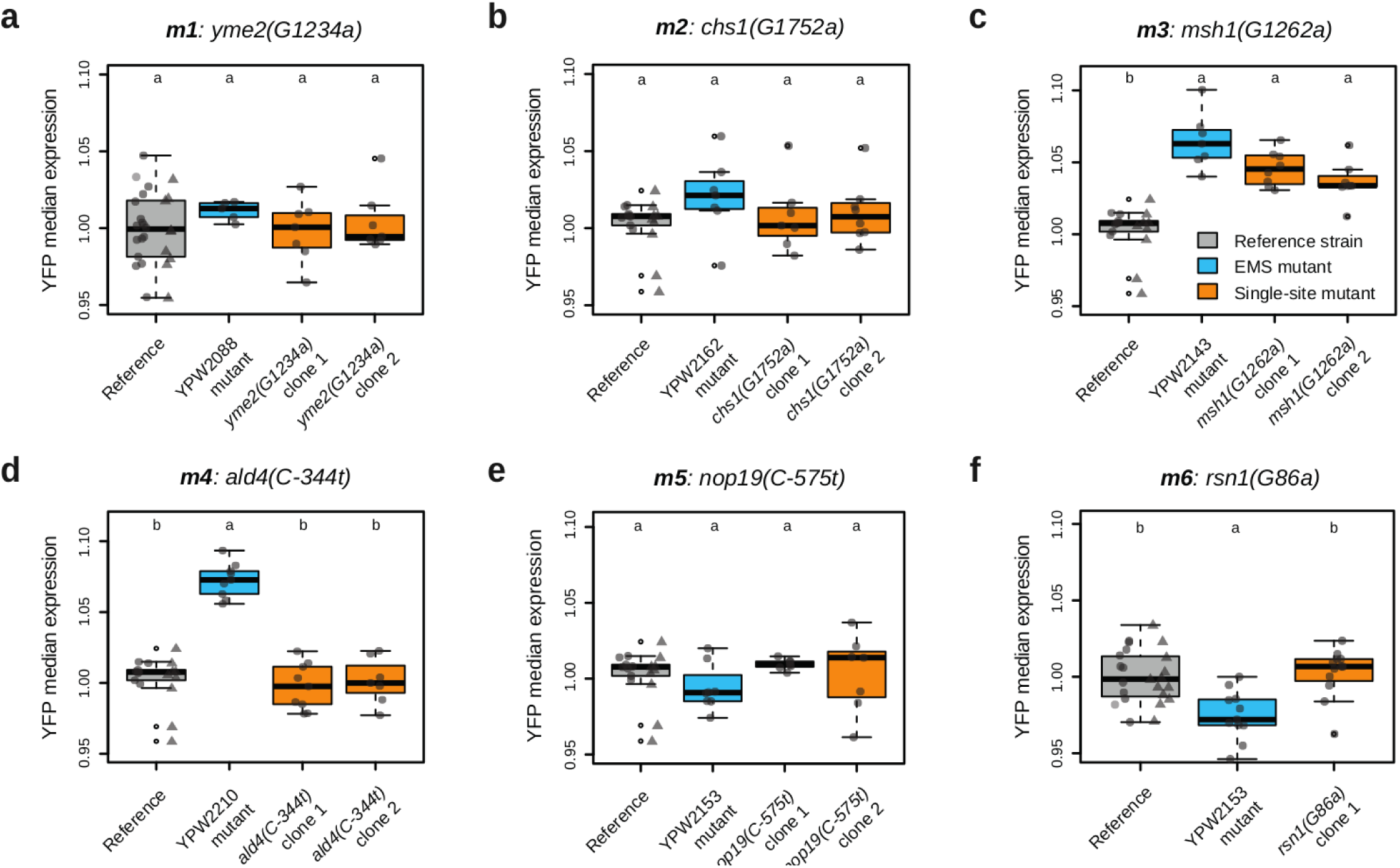
Testing the effects of six mapped mutations on YFP median expression using single-site mutants. Thick lines, boxes and whiskers represent the median, interquartile range (IQR) and 1.5 IQR of median expression measured among 7 to 26 samples, with colors corresponding to different genotypes as indicated in the legend of panel **(c)**. Dots represent the median expression of individual samples measured from 2102 to 4707 cells. For the reference genotype, circles and triangles represent data collected for two independent clones obtained from the same transformation experiment. For genotypes carrying single mutations, data collected for independent clones from the same transformation are shown in two orange boxes. Pairwise *t*-tests were performed to compare the mean of median expression levels among replicates between pairs of genotypes, with *P*-values adjusted using Benjamini-Hochberg correction. For each panel, the median expression is statistically different between two genotypes (adjusted *P*-value < 0.05) if no letter is shared above the boxes of the two genotypes. Above each plot, positions of mutations are indicated relative to the start codon of the gene with positive and negative numbers if mutations are respectively downstream (*i.e.* in coding sequence) and upstream of the start codon. Capital and lowercase letters indicate respectively reference and alternate nucleotides at each mutation site.

**Figure 2 – figure supplement 2.**
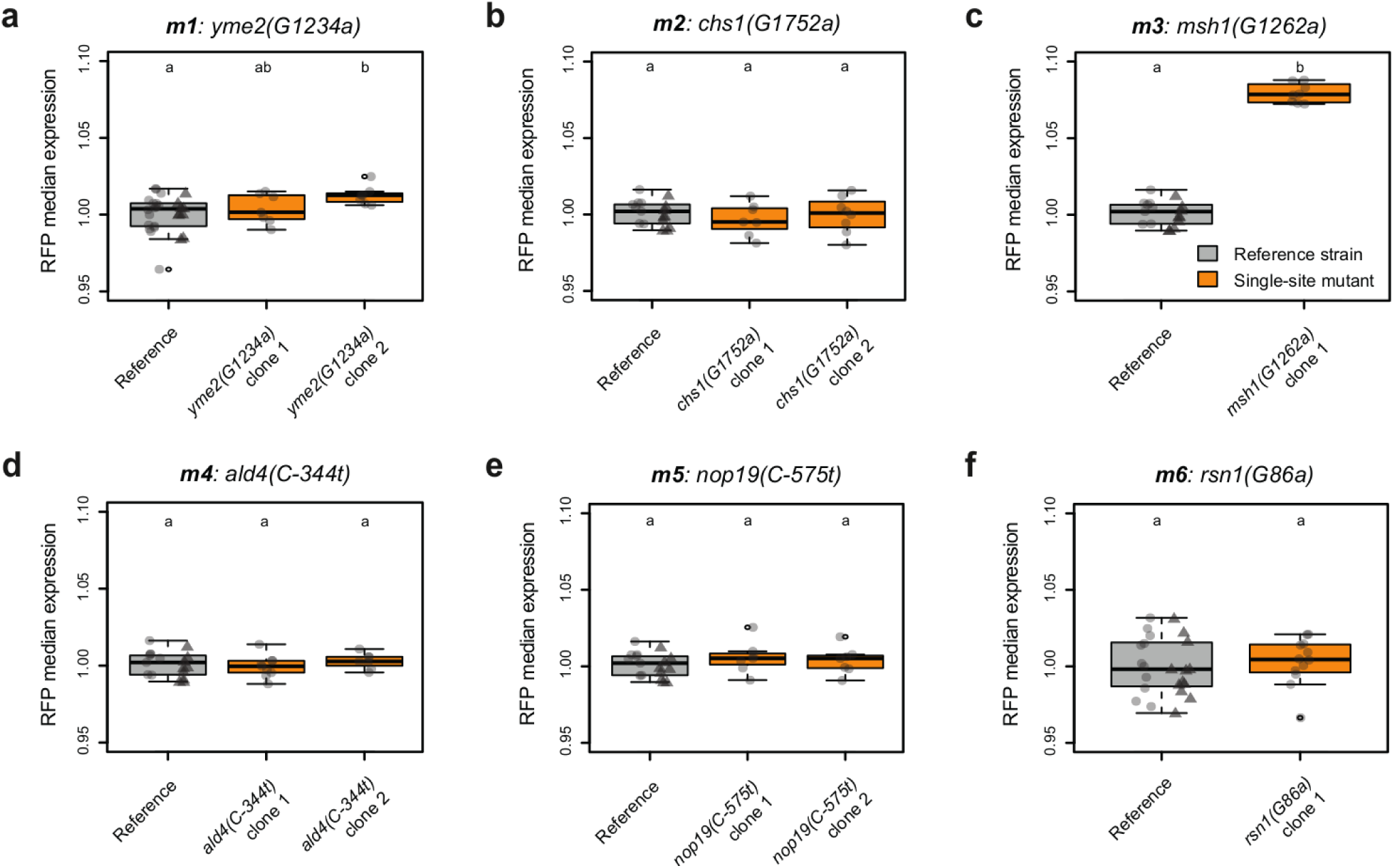
Testing the effects of six mapped mutations on RFP median expression and expression noise using single-site mutants. Thick lines, boxes and whiskers represent the median, interquartile range (IQR) and 1.5 IQR of median expression **(a-f)** or expression noise **(g-l)** measured among 7 to 26 samples, with colors corresponding to different genotypes as indicated in the legend of panel **(c)**. Dots represent the median expression or expression noise of individual samples measured from 2102 to 4707 cells. For the reference genotype, circles and triangles represent data collected for two independent clones obtained from the same transformation experiment. For genotypes carrying single mutations, data collected for independent clones from the same transformation are shown in two orange boxes. Pairwise *t*-tests were performed to compare the mean of median expression levels (or expression noise) among replicates between pairs of genotypes, with *P*-values adjusted using Benjamini-Hochberg correction. For each panel, the median expression (or expression noise) is statistically different between two genotypes (adjusted *P*-value < 0.01) if no letter is shared above the boxes of the two genotypes. Above each plot, positions of mutations are indicated relative to the start codon of the gene with positive and negative numbers if mutations are respectively downstream (*i.e.* in coding sequence) and upstream of the start codon. Capital and lowercase letters indicate respectively reference and alternate nucleotides at each mutation site.

**Figure 2 – figure supplement 3.**
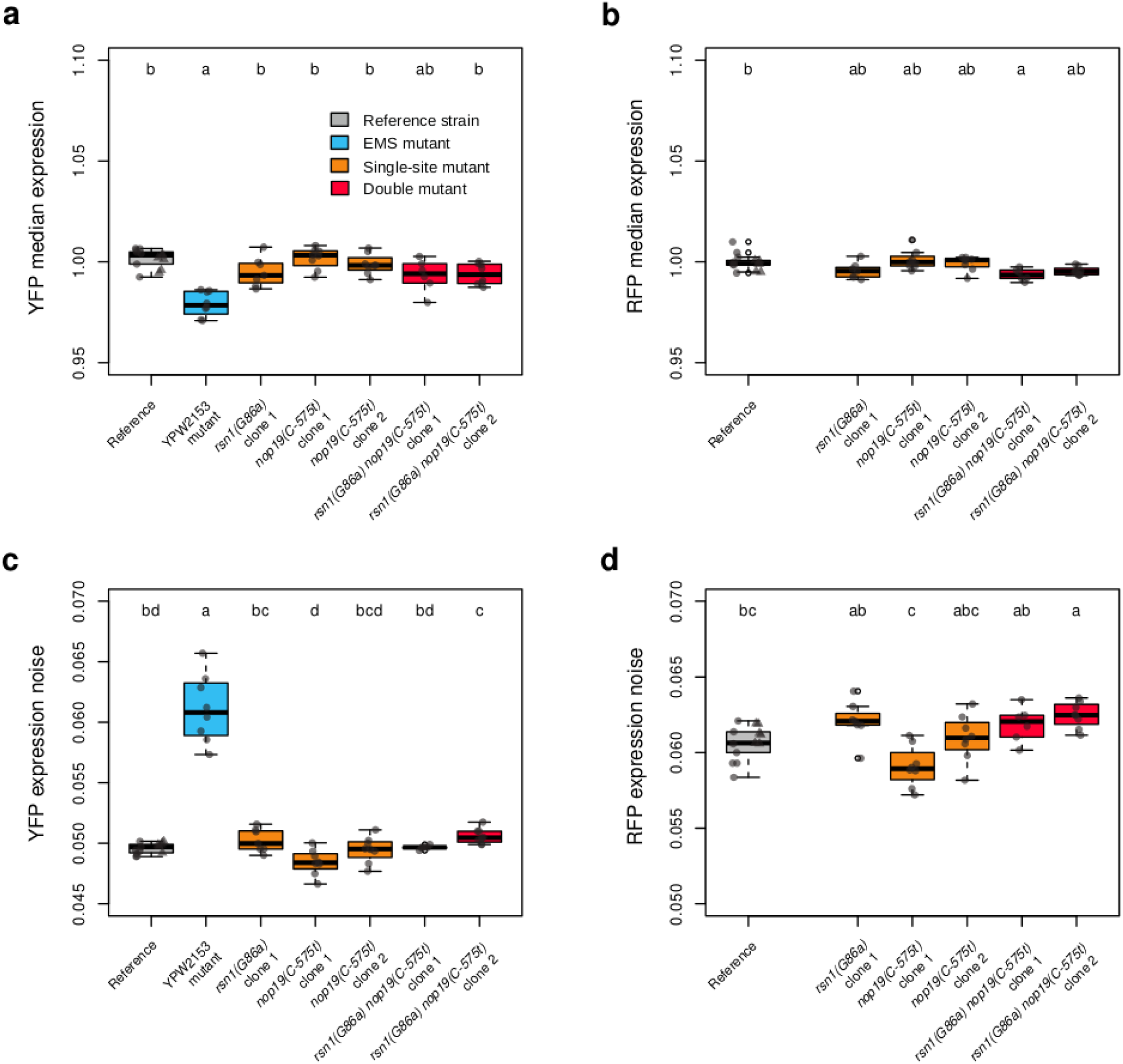
Combined effects of *rsn1(G86a)* and *nop19(C-575t)* mutations on YFP and RFP median expression and expression noise. Thick lines, boxes and whiskers represent the median, interquartile range (IQR) and 1.5 IQR of **(a)** YFP median expression, **(b)** RFP median expression, **(c)** YFP expression noise and **(d)** RFP expression noise measured among 6 to 14 samples. Colors corresponding to different genotypes as indicated in the legend of panel **(a)**. Dots represent expression values of individual samples measured from 4226 to 4551 cells. Expression noise was computed as the standard deviation divided by the median normalized fluorescence among cells, as described in Methods. For the reference genotype, circles and triangles represent data collected for two independent clones obtained from the same transformation experiment. For genotypes carrying single mutations, data collected for independent clones from the same transformation are shown in two orange boxes. Pairwise *t*-tests were performed to compare the mean of median expression levels among replicates between pairs of genotypes, with *P*-values adjusted using Benjamini-Hochberg correction. For each panel, the median expression is statistically different between two genotypes (adjusted *P*-value < 0.05) if no letter is shared above the boxes of the two genotypes. Above each plot, positions of mutations are indicated relative to the start codon of the gene with positive and negative numbers if mutations are respectively downstream (*i.e.* in coding sequence) and upstream of the start codon. Capital and lowercase letters indicate respectively reference and alternate nucleotides at each mutation site.

**Figure 2 – figure supplement 4.**
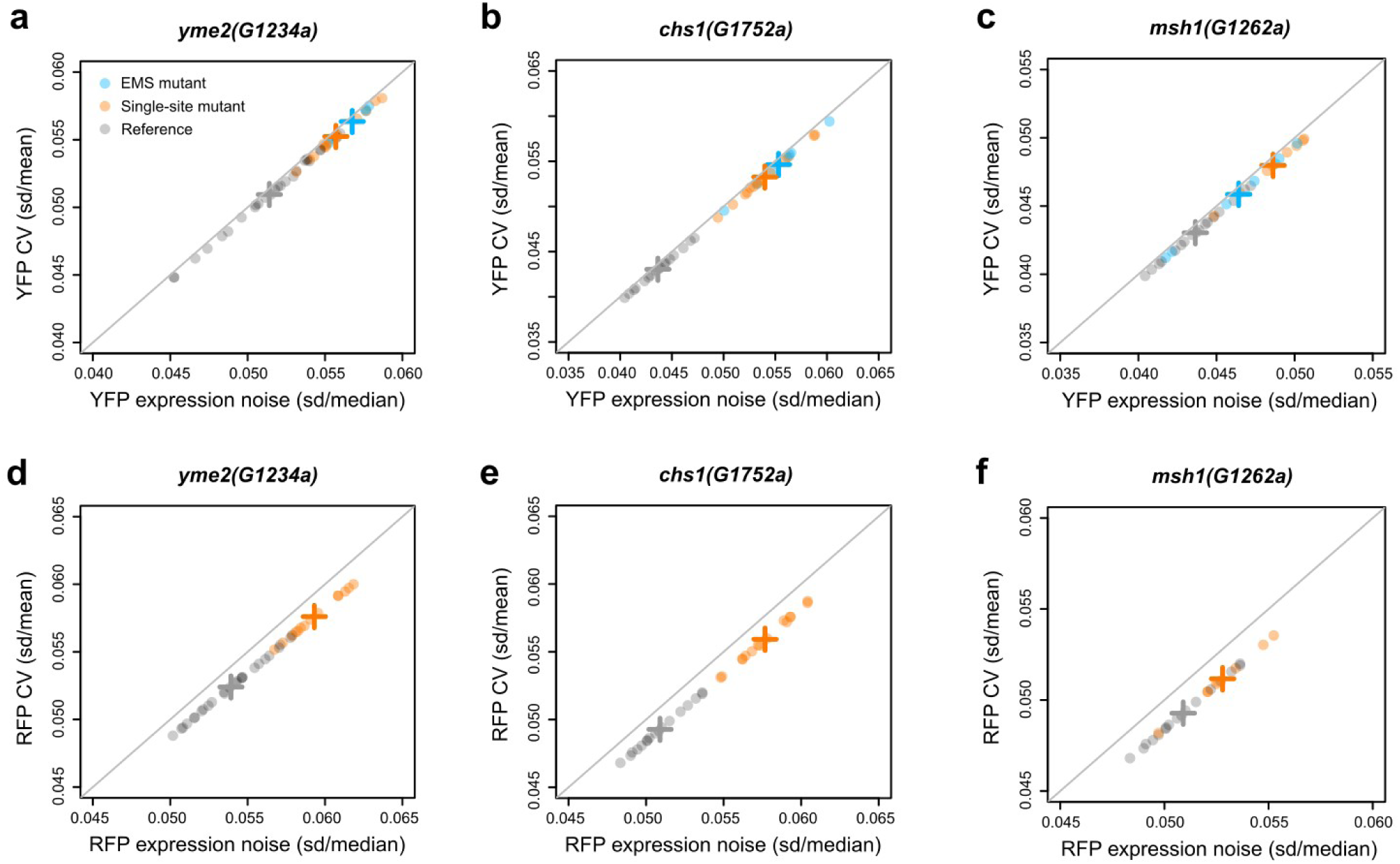
Comparison of two metrics of expression noise. The noise metric used in this study corresponds to the standard deviation divided by the median of normalized fluorescence among cells of each sample (x-axis). We compared this metric to the coefficient of variation corresponding to the standard deviation divided by the mean of normalized fluorescence among cells (y-axis). Data represented on this figure are the same as on Figure 2. Expression noise was measured for **(a-c)** YFP and for **(d-f)** RFP reporter genes in strains with wild-type or mutant alleles of **(a,d)** *yme2*, **(b,e)** *chs1* and **(c,f)** *msh1* genes. Dots represent noise values from each sample and thick crosses represent mean expression noise among all replicate samples for each genotype. Gray lines correspond to y = x functions. For each plot, the coefficient of determination (*R^2^*) was above 0.995.

For the other three candidate mutations tested (*m4*, *m5* and *m6*), no significant change in expression noise was observed between site-directed mutants and the reference strain for YFP (**Figure 2d-f**) and RFP (**Figure 2 – figure supplement 2j-l**); except for one of the two tested clones carrying mutation *m5* and only for RFP, which we considered to be a false positive. These mutations also did not significantly impact the median expression level of YFP (**Figure 2 – figure supplement 1d-f**) and RFP (**Figure 2 – figure supplement 2d-f**). Because mutations *m5* and *m6* were identified from the same EMS mutant, we hypothesized that these mutations may be required together to significantly impact expression noise. We therefore constructed a double mutant carrying both *m5* and *m6*, but this mutant did not show any significant change of expression noise relative to the reference strain ( **Figure 2 – figure supplement 3**). Several hypotheses may explain why we did not find a causative mutation for the two EMS mutants from which *m4*, *m5* and *m6* mutations were identified (see Discussion).

### A missense mutation in *YME2* gene increases intrinsic noise of *P_TDH3_-YFP* expression

To investigate potential mechanisms of action for each causative mutation, we next measured how these mutations affected two distinct sources of noise: intrinsic and extrinsic noise. Even though intrinsic and extrinsic noise both contribute to the total expression noise of a gene, they are confounded when expression is measured from a single allele. However, these sources of noise can be distinguished when measuring expression from two reporter genes (in our case *P_TDH3_-YFP* and *P_TDH3_-RFP*) in individual cells (**Figure 3a**), as previously demonstrated (Elowitz et al., 2002). Indeed, while intrinsic noise creates differences of expression levels between the two reporter genes within a cell, extrinsic noise creates expression variation among cells that is well correlated between the two reporters (**Figure 1c**). Intrinsic noise is expected to result from stochastic processes involved in the independent transcription, translation and post-translational regulation of each reporter gene, while extrinsic noise results from more global differences among cells such as differences in physiological states or in the amount of transcription factors regulating both reporters (Raser & O’Shea, 2005). We calculated intrinsic and extrinsic noise of mutant and reference strains from YFP and RFP fluorescence levels of individual cells as described in Fu & Pachter (2016) (see Methods).

**Figure 3.**
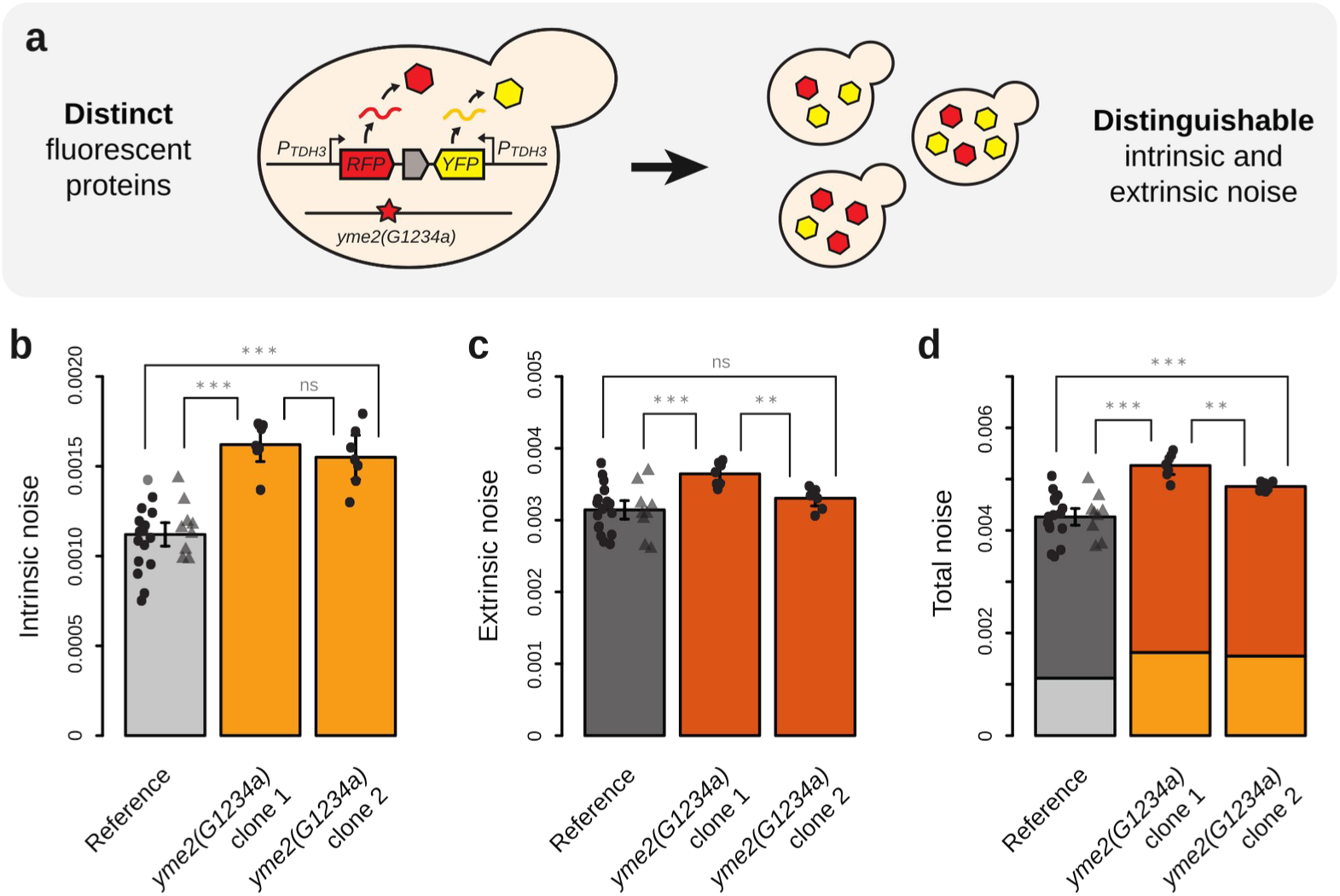
Quantifying the effects of a non-synonymous mutation in *yme2* on intrinsic and extrinsic noise using a dual reporter system. (a) A construct consisting of *P_TDH3_-RFP* and *P_TDH3_-YFP* reporter genes transcribed from opposite strands and separated by a *KanMX4* drug resistance cassette was inserted at the *ho* locus in genetic backgrounds with wild-type or mutant (*G1234a)* alleles of *yme2*. By quantifying both YFP and RFP fluorescence of individual cells using flow cytometry, it is possible to distinguish the effects of *yme2* mutation on intrinsic and extrinsic noise of *P_TDH3_* activity. **(b)** Intrinsic, **(c)** extrinsic and **(d)** total noise of *P_TDH3_* activity in strains carrying the dual reporter system with wild-type (light and dark gray bars) or mutant (light and dark orange bars) alleles of *yme2*. Bars show mean values among 7 to 26 replicate populations. Error bars are 95% confidence intervals of means. Dots show values of individual samples obtained from 4312 to 4689 cells. Light and dark bars indicate respectively the contributions of intrinsic and extrinsic noise to total noise. For the reference genotype, circles and triangles represent data collected for two independent clones obtained from the same transformation experiment. For *yme2* mutants, data collected for independent clones from the same transformation are shown in two bars with the same color. Pairwise *t*-tests were performed to compare mean values between bars: ns *P* ≥ 0.05, * 0.05 > *P* ≥ 0.01, ** 0.01 > *P* ≥ 0.001, *** *P* < 0.001.

**Figure 3 – figure supplement 1.**
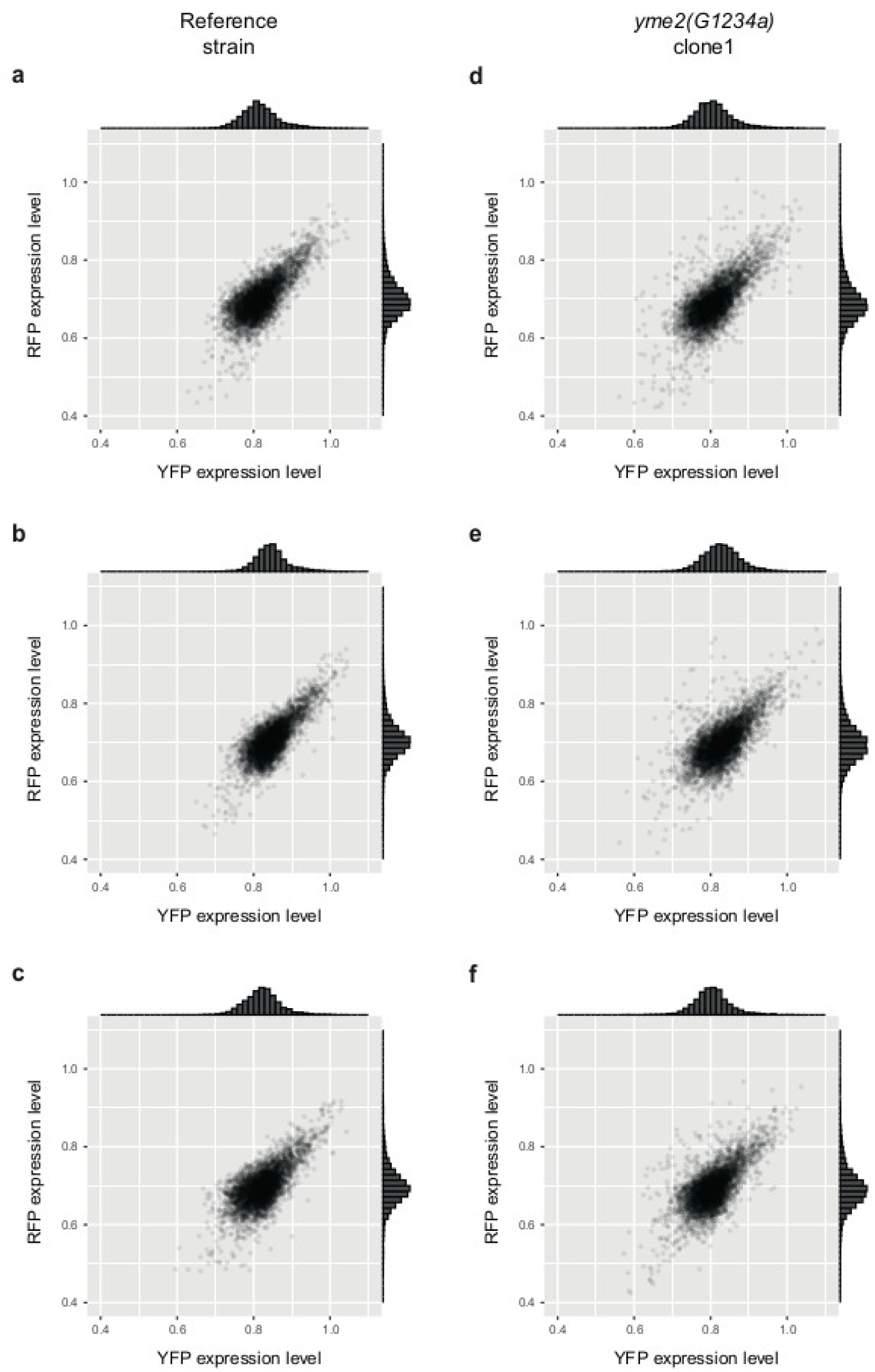
YFP and RFP expression levels of individual cells from wild-type and *yme2(G1234a)* strains expressing *P_TDH3_-YFP* and *P_TDH3_-RFP* reporter genes. Expression of the two fluorescent reporters in **(a-c)** three replicate populations of strain GY2458 carrying a wild-type allele of *YME2* and **(d-f)** three replicate populations of strain GY2634 carrying mutation *G1234a* in *YME2* coding sequence (4312 to 4689 cells per sample). Histograms show the distribution of data on each axis. Correlations between YFP and RFP expression levels are visually lower for *yme2(G1234a)* cells than wild-type cells, consistent with the increased intrinsic noise quantified from mutant cells.

**Figure 3 – figure supplement 2.**
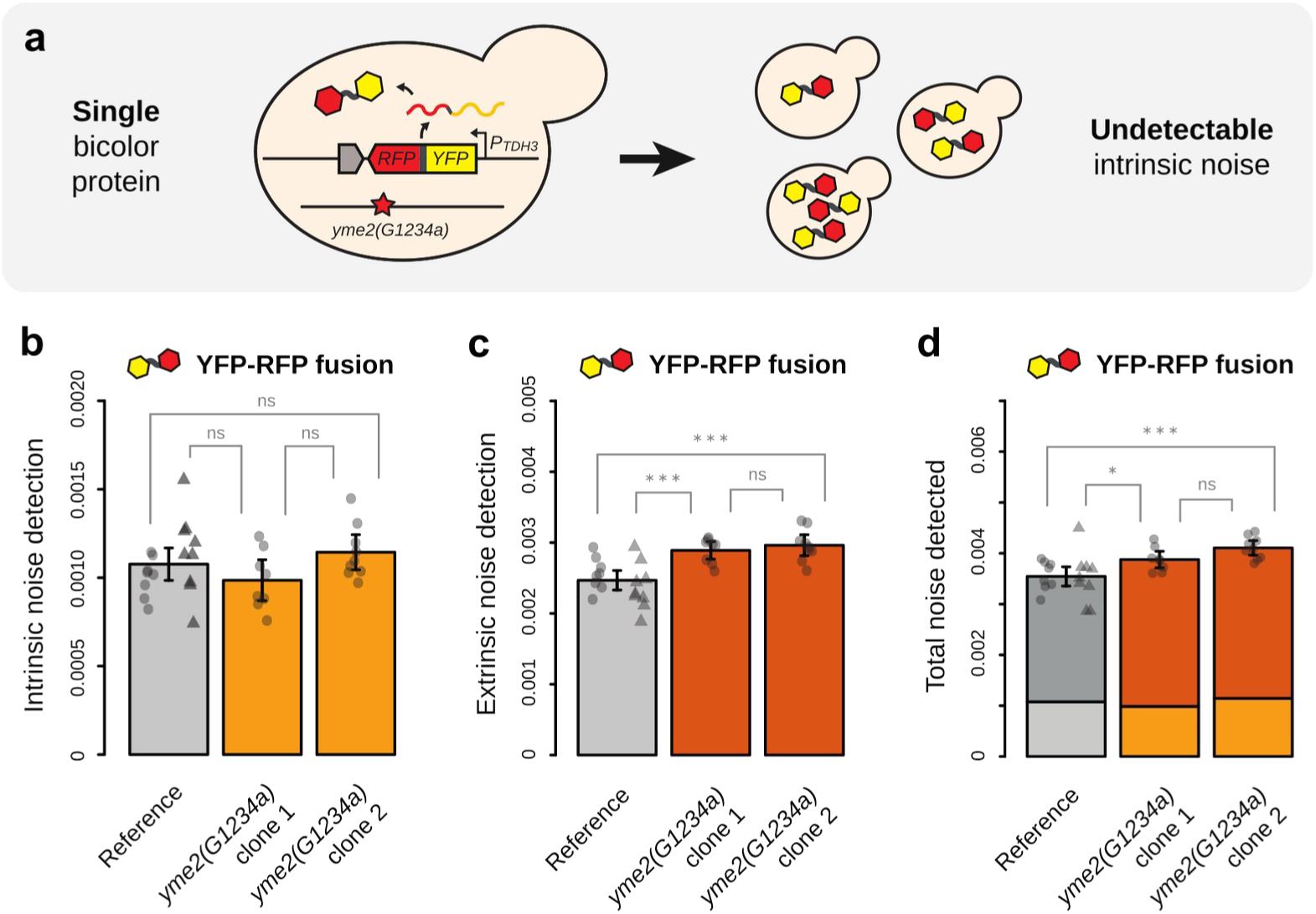
Quantifying the effects of a non-synonymous mutation in *yme2* on intrinsic and extrinsic noise using a dual reporter system. (a) A construct consisting of fused *YFP* and *RFP* coding sequences expressed under control of a single *P_TDH3_* copy was inserted at *ho* locus in genetic backgrounds with wild-type or mutant (*G1234a)* alleles of *yme2*. This system was used to test whether the change in intrinsic noise detected in Figure 3b using the dual reporter system was a technical artifact or not. Indeed, the mutation should not have detectable impact on intrinsic noise when the two fluorescent proteins are fused, because their expression is forced to covary. **(b)** Detected intrinsic noise, **(c)** detected extrinsic noise and **(d)** detected total noise in strains expressing YFP-RFP fusion proteins and with wild-type (light and dark gray bars) or mutant (light and dark orange bars) alleles of *yme2*. Bars show mean values among 8 to 17 replicate populations. Error bars are 95% confidence intervals of means. Dots show values of individual samples obtained from 4424 to 4714 cells. Light and dark bars indicate respectively the contributions of intrinsic and extrinsic noise to total noise. **(b-d)** For the reference genotype, circles and triangles represent data collected for two independent clones obtained from the same transformation experiment. For *yme2* mutants, data collected for independent clones from the same transformation are shown in two bars with the same color. Pairwise *t*-tests were performed to compare mean values between bars: ns *P* ≥ 0.05, * 0.05 > *P* ≥ 0.01, ** 0.01 > *P* ≥ 0.001, *** *P* < 0.001.

We first focused on mutation *m1* corresponding to a missense mutation located in the coding sequence of the *YME2* gene (*G1234a* substitution; **Table 3**). Site-directed mutants carrying the mutant allele *yme2(G1234a)* showed an increased intrinsic noise relative to the reference strain (**Figure 3b**; *t*-test, adjusted *P* < 0.001 for both clones). This effect is visible through the lower correlation observed between YFP and RFP expression levels of mutant cells than wild-type cells (**Figure 3 – figure supplement 1**). The impact of *yme2(G1234a)* mutation on extrinsic noise was less clear as only one of the two tested clones showed a significant increase in extrinsic noise (**Figure 3c**; *t*-test, adjusted *P* < 0.001 for clone 1 and adjusted *P* ⩾ 0.05 for clone 2). Overall, the impact of *yme2(G1234a)* mutation on intrinsic noise explained most of the variation in total expression noise of the two reporter genes (**Figure 3d**). This is particularly surprising since YME2p is an integral protein of the mitochondria inner membrane not known to regulate the expression of nuclear genes. YME2p is known to be involved in the fusion and fission of mitochondrial nucleoids and in the integrity of the mitochondrial genome (Hanekamp & Thorsness, 1996; Park et al., 2006). An indirect effect of mitochondrial activity on transcriptional or post-transcriptional regulation of nuclear genes would be expected to affect the expression of both reporter genes in a coordinated fashion, resulting in variation of extrinsic noise, which was not what we observed. Therefore, we tested whether experimental artifacts may explain the elevated intrinsic noise of *yme2(G1234a)* mutant using a YFP-RFP protein fusion (**Figure 3 – figure supplement 2**), but found no evidence it was the case (**Supplementary File 5**).

**Table 3.**
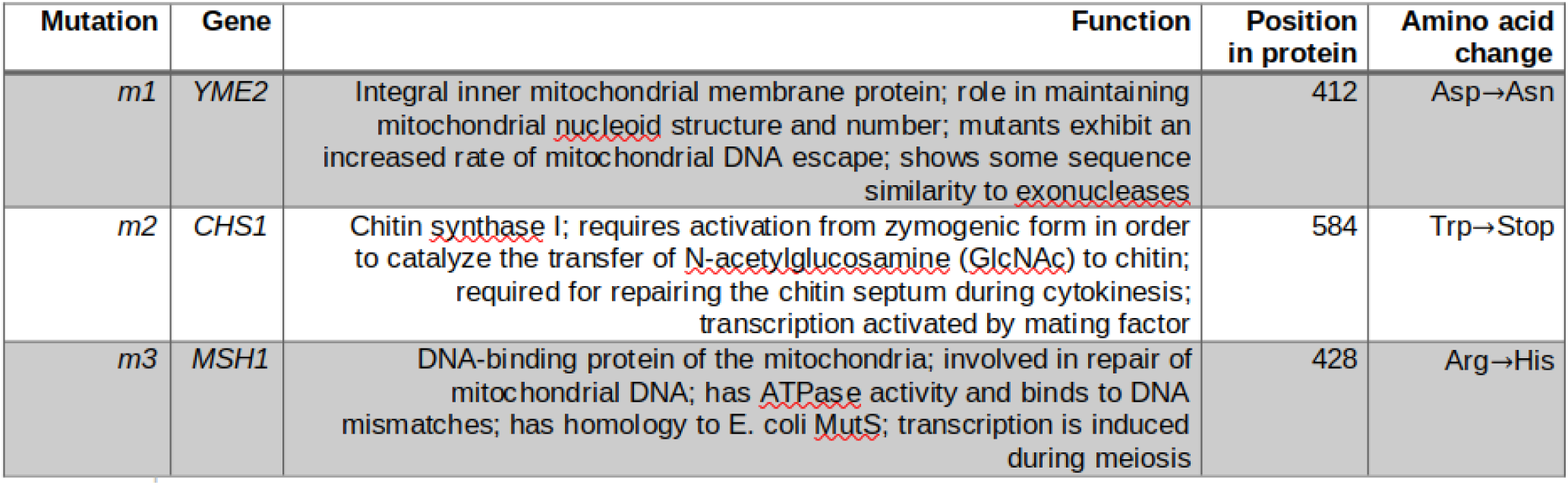
Mutations with confirmed effect on *P_TDH3_-YFP* expression noise.

| Mutation | Gene | Function | Position in protein | Amino acid change |
| --- | --- | --- | --- | --- |
| *m1* | *YME2* | Integral inner mitochondrial membrane protein; role in maintaining mitochondrial nucleoid structure and number; mutants exhibit an increased rate of mitochondrial DNA escape; shows some sequence similarity to exonucleases | 412 | Asp→Asn |
| *m2* | *CHS1* | Chitin synthase I; requires activation from zymogenic form in order to catalyze the transfer of N-acetylglucosamine (GlcNAc) to chitin; required for repairing the chitin septum during cytokinesis; transcription activated by mating factor | 584 | Trp→Stop |
| *m3* | *MSH1* | DNA-binding protein of the mitochondria; involved in repair of mitochondrial DNA; has ATPase activity and binds to DNA mismatches; has homology to E. coli MutS; transcription is induced during meiosis | 428 | Arg→His |

We noticed in flow cytometry data a small but statistically significant decrease of median cell size (estimated from forward scatter signal) in *yme2(G1234a)* mutants relative to the reference strain (**Figure 4a**). When clustering cells in five quantiles based on cell size for each genotype, we observed that the increase of intrinsic noise associated with *yme2(G1234a)* mutation was more pronounced in smaller cells (**Figure 4b**). The extrinsic noise of wild-type and mutant cells was indistinguishable regardless of their size, even though a negative relationship was globally observed between extrinsic noise and the size of genetically identical cells (**Figure 4c**). In budding yeast with asymmetric cell division, smallest cells in non-synchronized populations often correspond to daughter cells just after division. Although variation of intrinsic noise is typically thought to result from alteration in processes regulating gene expression (Hypothesis 1 in **Figure 4d**), an alternative hypothesis would be that the effect of *yme2(G1234a)* mutation on intrinsic noise was caused by altered partitioning of YFP and RFP molecules in daughter cells after division, potentially because of their reduced size compared to wild-type daughter cells (Hypothesis 2 in **Figure 4d**). To test these hypotheses, we constructed a *P_TDH3_-YFP-RFP* gene fusion where coding sequences of *YFP* and *RFP* were separated by a short sequence encoding the cleavage site for the tobacco etch virus (TEV) protease (ENLYFQS peptide sequence). This construct was inserted at *ho* locus in strains that constitutively express the wild-type TEV or the improved uTEV3 protease from Sanchez and Ting (2020), with either the wild-type or *G1234a* mutant allele of *YME2*. We reasoned that sources of noise acting on expression at any step from transcription to translation would be undetectable in these strains because YFP and RFP are expressed from the same coding sequence. However, if the increased intrinsic noise was caused by altered partitioning of YFP and RFP molecules, we should still be able to detect it in these strains because YFP-RFP proteins are cleaved into YFP and RFP peptides that are free to segregate independently in mother and daughter cells after division (**Figure 4d**, **Supplementary File 5**). Western blotting with an antibody against RFP showed that YFP-RFP was fully cleaved in strains expressing the improved uTEV3 protease (undetectable level of YFP-RFP fusion protein), but only partially cleaved in strains expressing the wild-type TEV (**Figure 4e**). Therefore, we measured expression noise in strains expressing uTEV3. We observed no detectable effect of mutation *yme2(G1234a)* on intrinsic noise in strains producing YFP and RFP peptides from cleavage of YFP-RFP fusion proteins (**Figure 4f**; *t*-test, adjusted *P* > 0.05), regardless of cell size (**Figure 4h**). However, a significant increase of extrinsic noise was observed for *yme2(G1234a)* strains relative to wild-type strains (**Figure 4g**; *t*-test, adjusted *P* < 0.001 for clone 1, adjusted *P* < 0.01 for clone 2). These results are not consistent with the hypothesis that *yme2(G1234a)* mutation affects expression noise via altered partitioning of proteins during cell division. Instead, the mutation must cause an increased intrinsic noise by acting on a process regulating gene expression at some point from transcription to protein degradation, via an unknown mechanism.

**Figure 4.**
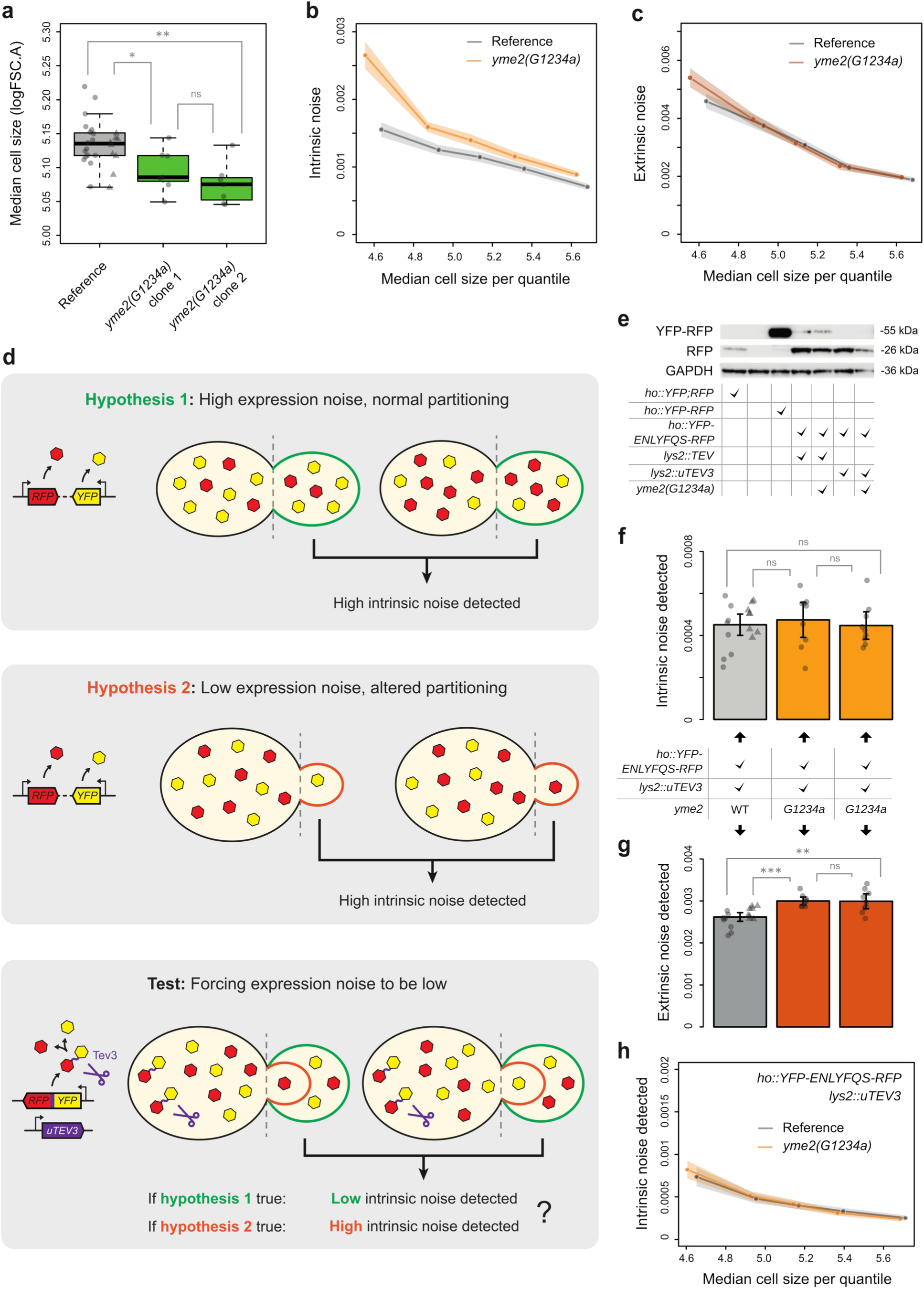
***yme2(G1234a)* mutation increases intrinsic noise via altered gene expression and not via altered protein partitioning at cell division.** (**a**) Median size of wild-type and *yme2(G1234a)* mutant cells quantified from forward scatter area (FSC.A) in flow cytometry data. Thick lines, boxes and whiskers represent the median, interquartile range (IQR) and 1.5 IQR of median cell size measured among 7 to 26 samples. Dots represent the median cell size of individual samples measured from 4312 to 4689 cells. For the reference genotype, circles and triangles represent data collected for two independent clones obtained from the same transformation experiment. For genotypes carrying single mutations, data collected for independent clones from the same transformation are shown in two green boxes. Pairwise *t*-tests were performed to compare median cell sizes between boxes: ns *P* ≥ 0.05, * 0.05 > *P* ≥ 0.01, ** 0.001 > *P* ≥ 0.001, *** *P* < 0.001. **(b)** Intrinsic noise and **(c)** extrinsic noise of five groups of cells with different cell sizes for strains with wild-type (gray) and *G1234a* (orange) alleles of *yme2*. Cell groups correspond to five fractions of the population of cells delimited by 5-quantiles of cell sizes (same number of cells in each group but different cell sizes). Dots indicate the median cell size (x-axis) and intrinsic **(b)** or extrinsic **(c)** noise (y-axis) of each cell group (906 cells per group in 24 replicate samples for wild-type and 909 cells per group in 18 replicate samples for *yme2* mutant). Shaded areas represent 95% confidence intervals. **(d)** Schemes representing two alternative hypotheses to explain how *yme2(G1234a)* mutation can lead to higher intrinsic noise detection and a strategy to test these hypotheses (see **Supplementary File 5** for a detailed rationale). **(e)** Western blot used to detect the presence of YFP-RFP fusion protein, RFP single protein and GAPDH constitutive control in seven genotypes (as indicated below the plot). YFP-RFP and RFP are detected using an antibody against mCherry and distinguished based on the size of bands after electrophoresis. GAPDH is detected with anti-GPADH antibody. The last two lanes on the right show complete cleavage of YFP-ENLYFQS-RFP fusion protein by uTEV3 protease both in wild-type and *yme2(G1234a)* cells. ENLYFQS is the peptide sequence targeted by the protease. **(f)** Detected intrinsic noise and **(g)** extrinsic noise in strains expressing YFP-ENLYFQS-RFP fusion protein cleaved by uTEV3 and with wild-type (light and dark gray bars) or mutant (light and dark orange bars) alleles of *yme2*. Bars show mean values among 9 to 16 replicate populations. Error bars are 95% confidence intervals of means. Dots show values of individual samples obtained from 4061 to 4673 cells. For the reference genotype, circles and triangles represent data collected for two independent clones obtained from the same transformation experiment. For *yme2(G1234a)* strains, data collected for independent clones from the same transformation are shown in two bars with the same color. Pairwise *t*-tests were performed to compare mean values between bars: ns *P* ≥ 0.05, * 0.05 > *P* ≥ 0.01, ** 0.01 > *P* ≥ 0.001, *** *P* < 0.001. **(h)** Intrinsic noise of five groups with different cell sizes for strains expressing YFP-ENLYFQS-RFP fusion protein cleaved by uTEV3 with wild-type (gray) and mutant *G1234a* (orange) alleles of *yme2*. Cell groups correspond to five fractions of the population of cells delimited by 5-quantiles of cell sizes. Dots indicate the median cell size (x-axis) and intrinsic noise (y-axis) of each cell group (902 cells per group in 17 replicate samples per group for wild-type and 906 cells per group in 18 replicate samples per group for *yme2(G1234a)*). Shaded areas represent 95% confidence intervals of intrinsic noise.

### A nonsense mutation in *CHS1* increases extrinsic noise via impaired chitin septum reparation in daughter cells

We next quantified the impact of mutation *m2* on extrinsic and intrinsic noise using site-directed mutants that expressed both *P_TDH3_-YFP* and *P_TDH3_-RFP*. *m2* is a nonsense mutation in the *CHS1* gene (*G1752a* substitution) that introduces an early stop codon reducing the open reading frame by 48.4% (**Table 3**). We found that *chs1(G1752a)* allele had no clear impact on intrinsic noise (**Figure 5a**; *t*-test, adjusted *P* ⩾ 0.05 for clone 1 and *P* < 0.01 for clone 2 with decreased intrinsic noise). However, it significantly increased extrinsic noise (**Figure 5b**; *t*-test, adjusted *P* < 0.01 for clone 1 and *P* < 0.001 for clone 2) and total noise (**Figure 5c**; *t*-test, adjusted *P* < 0.01 for clone 1 and *P* < 0.001 for clone 2) relative to the reference allele. Consistent with this increase in extrinsic noise, *chs1(G1752a)* cells showed stronger correlation of YFP and RFP expression levels than wild-type cells, with a higher proportion of mutant cells showing either low or high expression of both reporters (**Figure 5 – figure supplement 1**). As expected, we also detected an increased extrinsic noise in *chs1(G1752a)* mutant cells that expressed a YFP-RFP fusion protein (**Figure 5 – figure supplement 2**).

**Figure 5.**
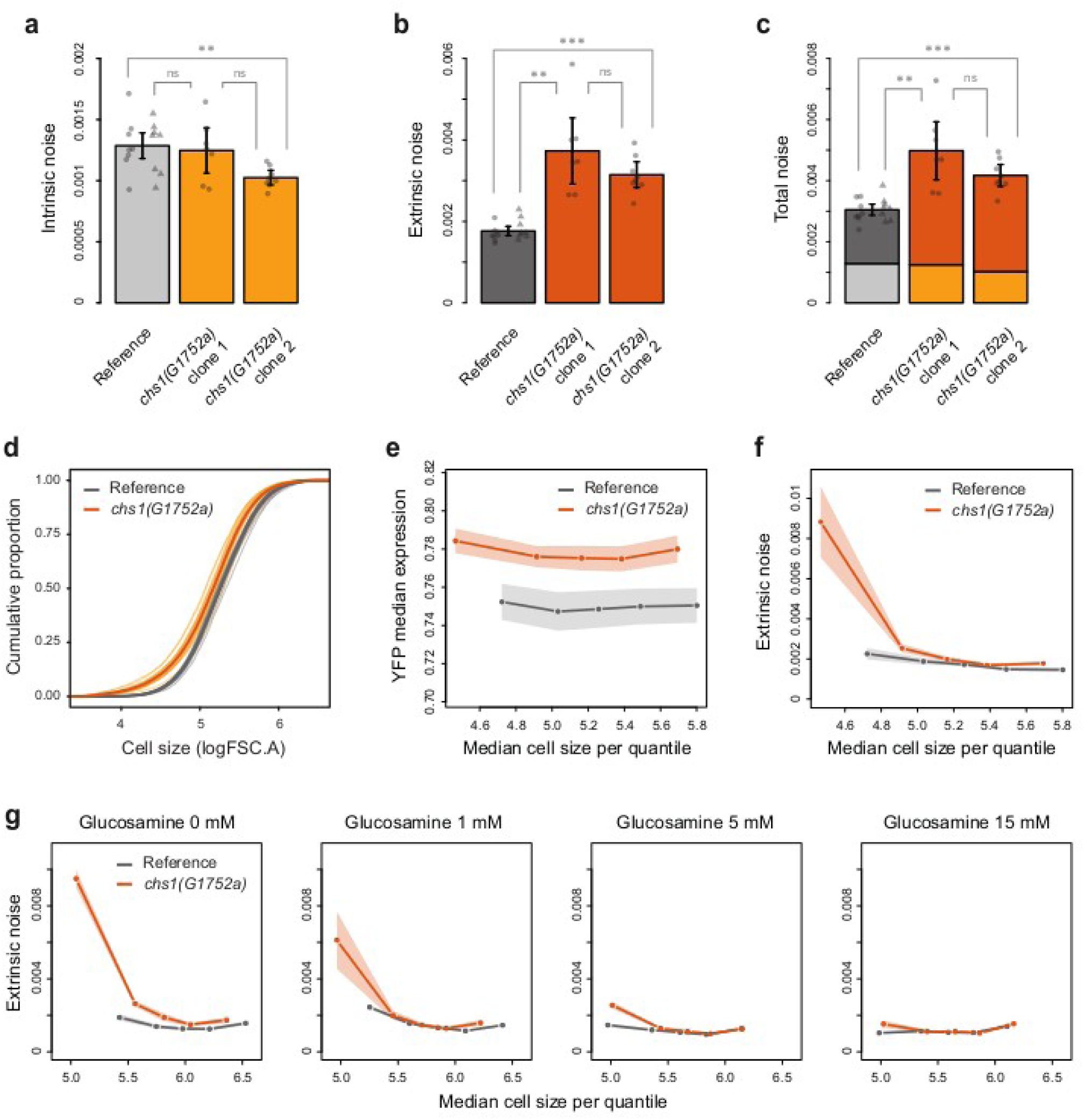
***chs1(G1752a)* mutation increases extrinsic noise exclusively in small cells in absence of glucosamine precursor for chitin synthesis.** (**a**) Intrinsic, **(b)** extrinsic and **(c)** total noise of *P_TDH3_* activity in strains carrying the dual reporter system (*P_TDH3_-YFP* and *P_TDH3_-RFP*) with wild-type (light and dark gray bars) or mutant (light and dark orange bars) alleles of *chs1*. Bars show mean values among 7 to 16 replicate populations. Error bars are 95% confidence intervals of means. Dots show values of individual samples obtained from 3684 to 4657 cells. **(d)** Cumulative distribution of cell sizes for wild-type (gray lines) and *chs1(G1752a)* cells (orange lines) expressing the dual reporter system, shown for individual samples (thin lines) or for all samples sharing the same genotype (thick lines). Cell sizes were quantified by flow cytometry (FSC.A: area of forward scatter). The number of samples and cells analyzed is the same as in **(a-c)** as data are from the same experiment. **(e)** YFP median expression and **(f)** extrinsic noise of five groups of cells with different cells sizes for strains with wild-type (gray) and *G1752a* (orange) alleles of *chs1*. Cell groups correspond to five fractions of the population of cells delimited by 5-quantiles of cell sizes (same number of cells in each group but different cell sizes). Dots indicate the median cell size (x-axis) and YFP median expression **(e)** or extrinsic **(c)** noise (y-axis) of each cell group (1790 cells per group in 16 replicate samples for wild-type and 1603 cells per group in 15 replicate samples for *chs1* mutant). **(g)** Extrinsic noise of cells of different sizes measured after growth in media with different concentrations of glucosamine (0, 1, 5 or 15 mM) for strains with wild-type (gray) and *G1752a* (orange) alleles of *chs1*. Cells were grouped based on median cell size as explained for **(e-f)**. Dots indicate the median cell size (x-axis) and extrinsic noise (y-axis) for 1660 to 1878 cells per group measured in 5 to 16 replicates. **(e-g)** Shaded areas represent 95% confidence intervals.

**Figure 5 – figure supplement 1.**
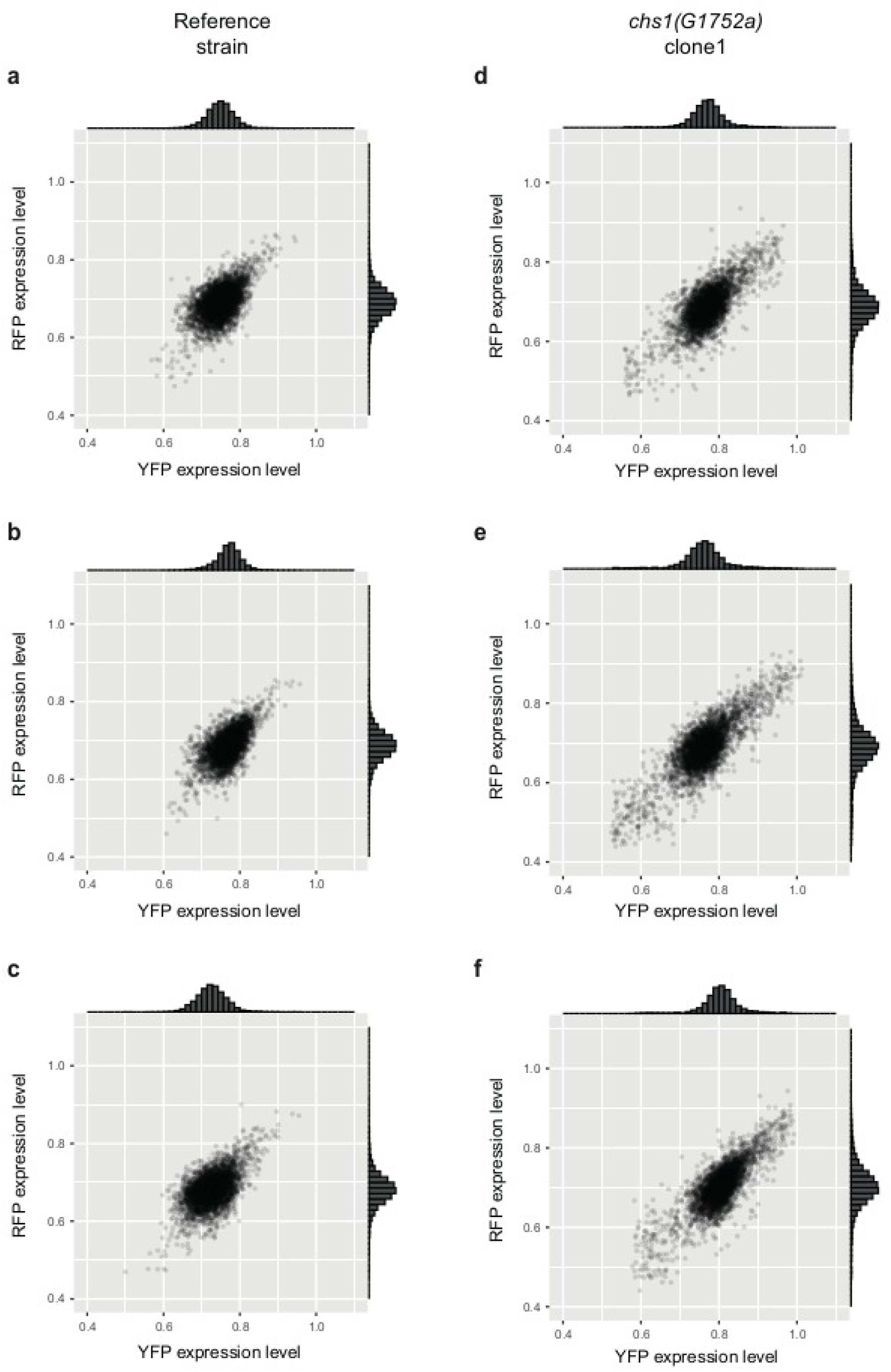
YFP and RFP expression levels of individual cells from wild-type and *chs1(G1752a)* strains expressing *P_TDH3_-YFP* and *P_TDH3_-RFP* reporter genes. Expression of the two fluorescent reporters in **(a-c)** three replicate populations of strain GY2458 carrying a wild-type allele of *CHS1* and **(d-f)** three replicate populations of strain GY2534 carrying mutation *G1752a* in *CHS1* coding sequence (3917 to 4627 cells per sample). Histograms show the distribution of data on each axis. The variability and covariation of YFP and RFP expression levels are both visually higher for *chs1(G1752a)* cells than for wild-type cells, consistent with the increased extrinsic noise quantified from mutant cells.

**Figure 5 – figure supplement 2.**
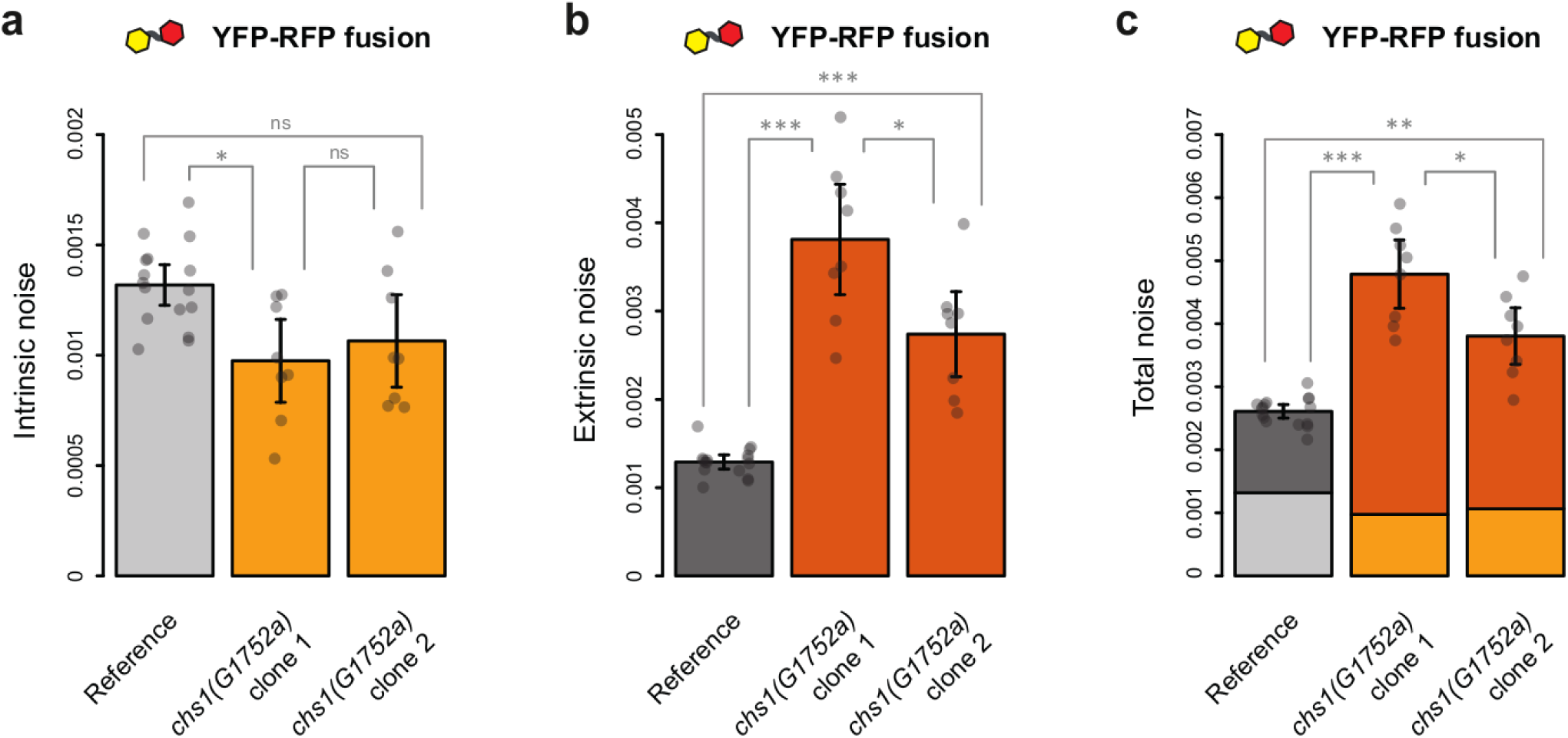
Fusion of YFP and RFP fluorescent proteins does not change the detected effect of *chs1(G1752a)* mutation on extrinsic noise. (a) Detected intrinsic noise, **(b)** detected extrinsic noise and **(c)** detected total noise in strains expressing YFP-RFP fusion proteins and with wild-type (light and dark gray bars) or mutant (light and dark orange bars) alleles of *chs1*. Bars show mean values among 8 to 16 replicate populations. Error bars are 95% confidence intervals of means. Dots show values of individual samples obtained from 4196 to 4899 cells. Pairwise *t*-tests were performed to compare mean values between bars: ns *P* ≥ 0.05, * 0.05 > *P* ≥ 0.01, ** 0.001 > *P* ≥ 0.001, *** *P* < 0.001.

**Figure 5 – figure supplement 3.**
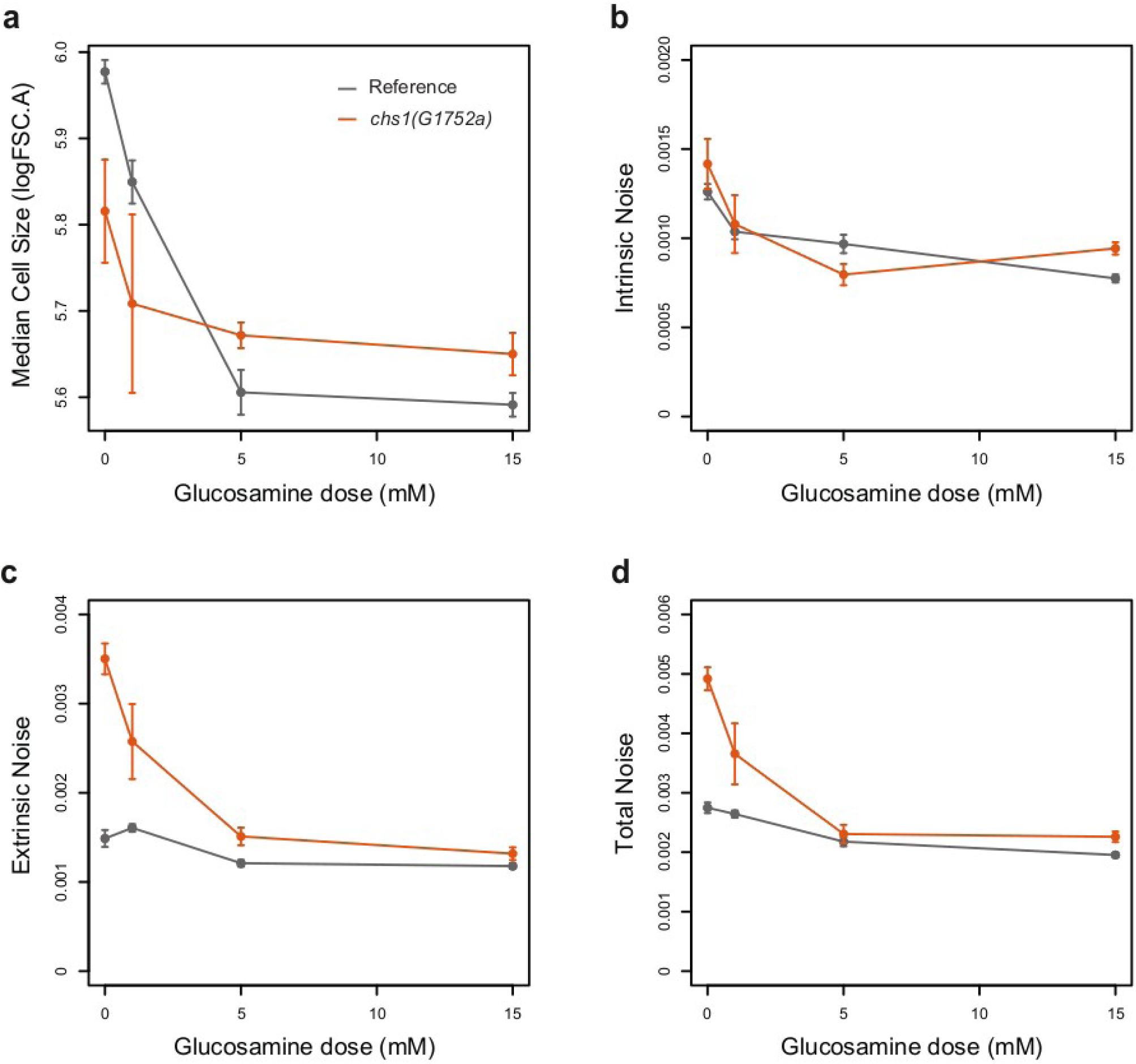
Impact of glucosamine concentration on (a) median cell size, (b) intrinsic noise, (c) extrinsic noise and (d) total noise. These traits were quantified in wild-type (gray dots) and *chs1(G1752a)* (orange dots) strains expressing *P_TDH3_-YFP* and *P_TDH3_-RFP* reporter genes. Error bars show 95% confidence intervals of means measured among 5 to 16 replicate samples.

**Figure 5 – figure supplement 4.**
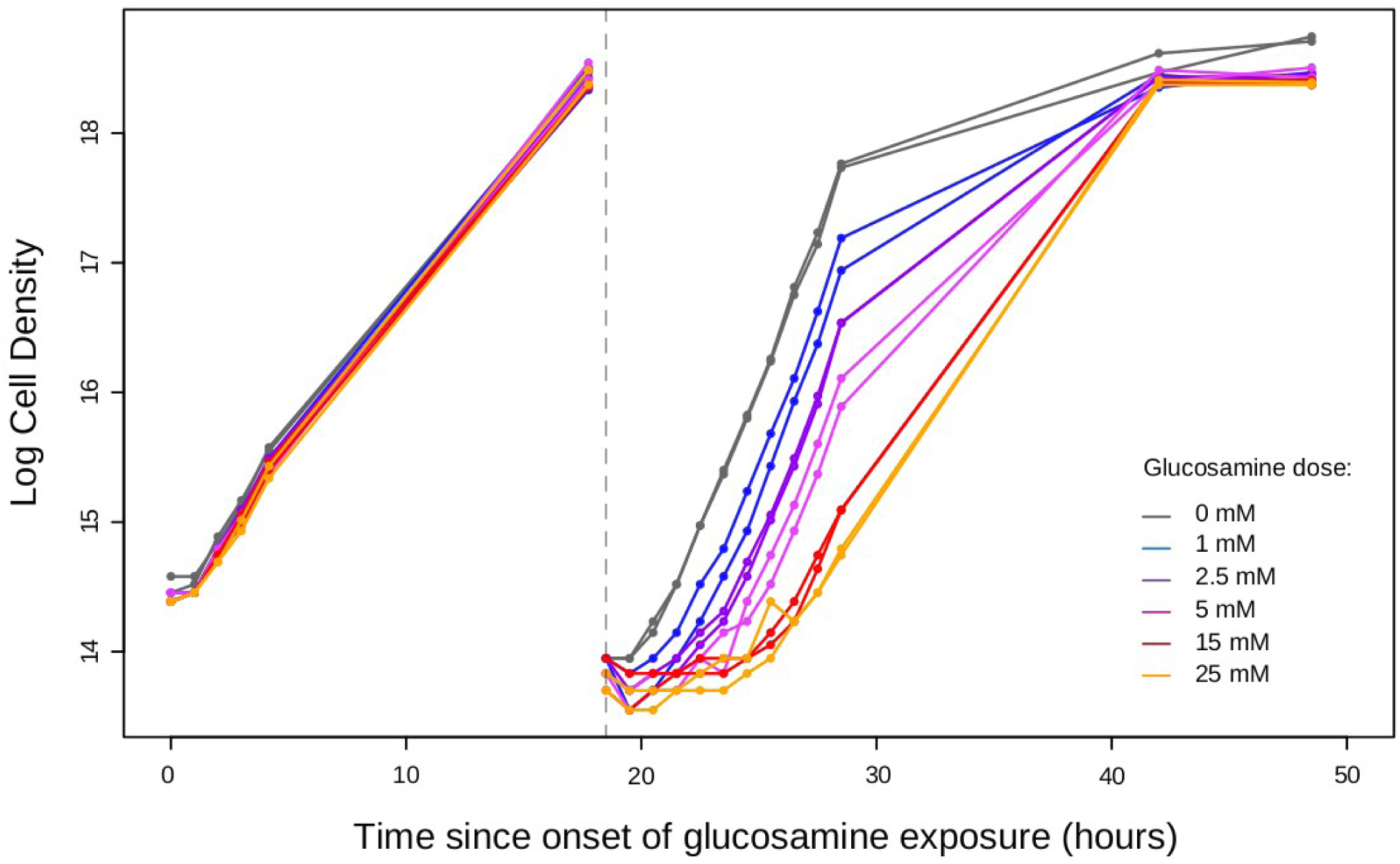
Growth of wild-type cell populations exposed to different concentrations of glucosamine. Densities of cell cultures were quantified at different time points (dots) over two cycles of growth using a spectrophotometer. At *t* = 0, cells were transferred to different media containing the indicated concentrations of glucosamine (colors), then grown for 18.5 hours, diluted to fresh media containing the same concentrations of glucosamine and grown again for 30 hours.

**Figure 6.**
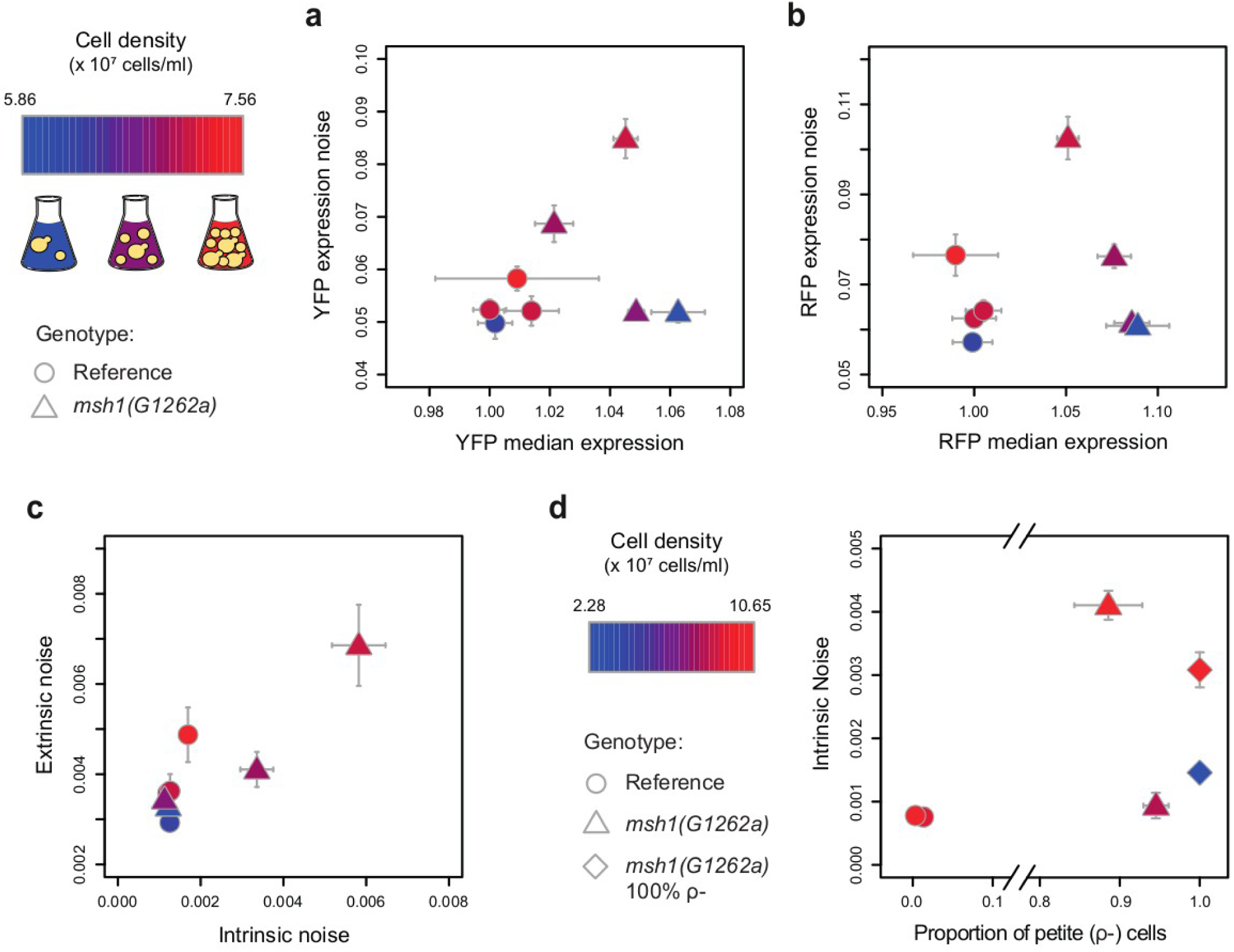
The effect of *msh1(G1262a)* mutation on expression noise depends on cell density. Expression noise was quantified either from YFP fluorescence **(a)**, RFP fluorescence **(b)** or both **(c)** in strains carrying the dual reporter system (*P_TDH3_-YFP* and *P_TDH3_-RFP*) with wild-type (circles) or mutant (triangles) alleles of *msh1*. **(d)** Intrinsic noise plotted against the proportion of respiratory deficient cells (petites ρ-) for a strain with wild-type allele of *msh1* (circle), a strain derived from a ρ+ cell with *msh1(G1262a)* mutation (triangle) and a strain derived from a ρ-cell with *msh1(G1262a)* mutation (diamond). The proportion of ρ-cell was quantified for each strain as described in Materials and methods. **(a-d)** Dots indicate mean values among 4 replicates and error bars are 95% confidence intervals of the mean. Colors indicate the average density of cell cultures measured after growth and just before fixing cells to quantify their fluorescence levels, with color legend shown on top left for **(a-c)** and at the bottom for **(d)**.

**Figure 6 – figure supplement 1.**
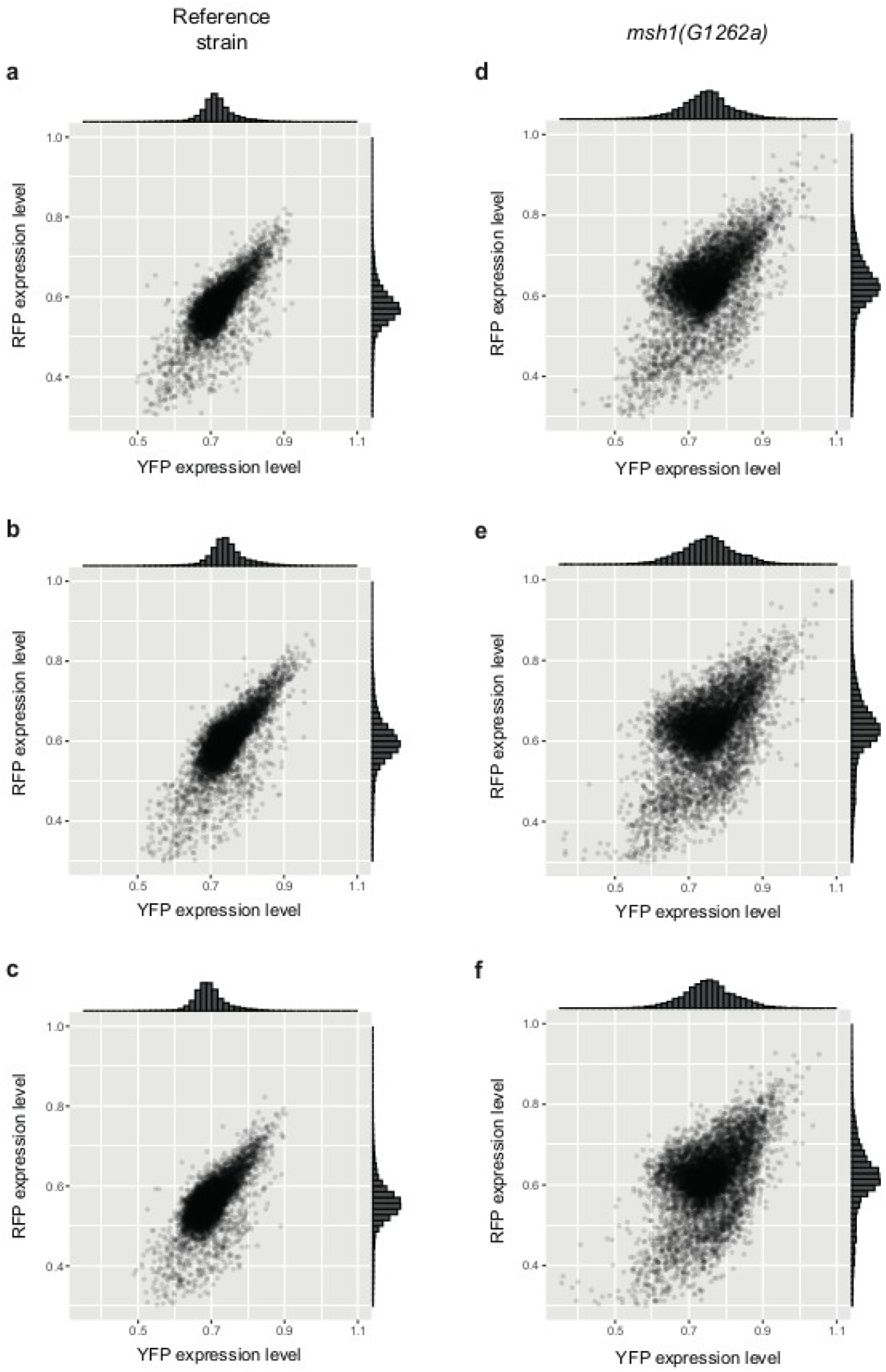
YFP and RFP expression levels of individual cells from wild-type and *msh1(G1262a)* strains expressing *P_TDH3_-YFP* and *P_TDH3_-RFP* reporter genes. Expression of the two fluorescent reporters in **(a-c)** three replicate populations of strain GY2458 carrying a wild-type allele of *MSH1* and **(d-f)** three replicate populations of strain GY2534 carrying mutation *G1262a* in *MSH1* coding sequence (7885 to 8394 cells per sample). Histograms show the distribution of data on each axis.

**Figure 6 – figure supplement 2.**
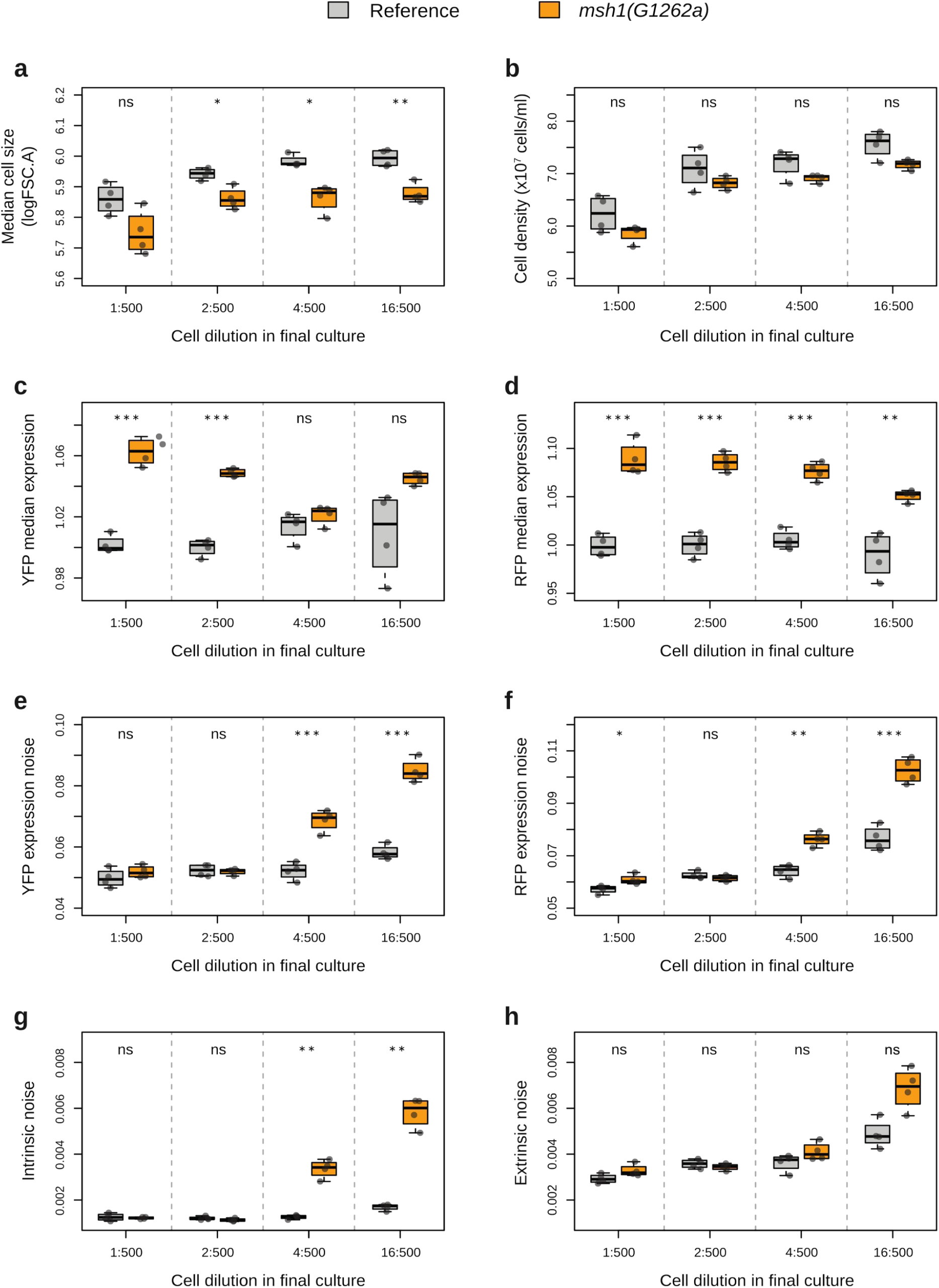
Effect of *msh1(G1262a)* mutation on 8 quantitative traits depending on cell culture dilutions. (a) Median cell size, **(b)** cell density, **(c)** YFP median expression, **(d)** RFP median expression, **(e)** YFP expression noise, **(f)** RFP expression noise, **(g)** intrinsic noise and **(h)** extrinsic noise were quantified in strains with the dual reporter system carrying wild-type (gray) or *G1262a* (orange) alleles of *msh1*. **(a-h)** Thick lines, boxes and whiskers represent the median, interquartile range (IQR) and 1.5 IQR of traits indicated on y-axis measured among 4 replicate samples. Dots represent mean values for individual samples measured from 7203 to 9218 cells. Starting from a culture at saturation (∼10^8^ cells/ml), cells were diluted to 500 µl of fresh SD medium at 4 different dilution ratios as indicated on x-axis and incubated for another 18 hours at 30°C before fixation and phenotyping by flow cytometry. As intended, the different dilution ratios resulted in different cell densities after incubation as shown in **(b)**. Pairwise *t*-tests were performed to compare mean phenotypic values among replicates between wild-type and *msh1(G1262a)* strains for each dilution ratio, with p-values adjusted using Benjamini-Hochberg correction: ns *P* ≥ 0.05, * 0.05 > *P* ≥ 0.01, ** 0.001 > *P* ≥ 0.001, *** *P* < 0.001.

**Figure 6 – figure supplement 3.**
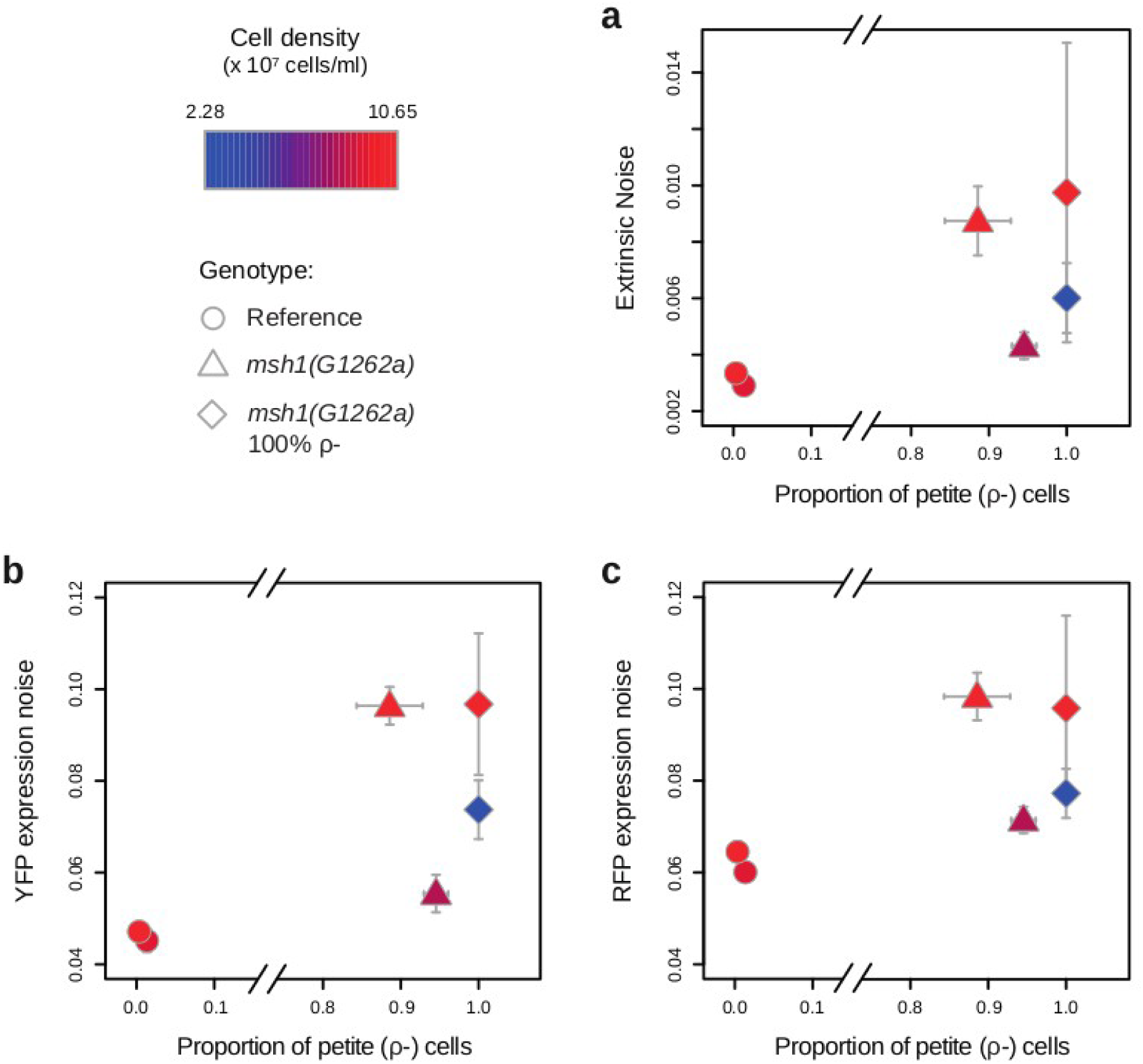
Cell density changes the effect of *msh1(G1262a)* on expression noise independently from an increased proportion of respiratory deficient cells. (a) Extrinsic noise, **(b)** YFP expression noise and **(c)** RFP expression noise plotted against the proportion of respiratory deficient cells (petites ρ-) for a strain with wild-type allele of *msh1* (circle), a strain derived from a ρ+ cell with *msh1(G1262a)* mutation (triangle) and a strain derived from a ρ-cell with *msh1(G1262a)* mutation (diamond). The proportion of ρ-cell was quantified for each strain as described in Materials and methods. Dots indicate mean values among 4 replicates and error bars are 95% confidence intervals of the mean. Colors indicate the average density of cell cultures measured after growth and just before fixing cells to quantify their fluorescence levels, with color legend shown on top left.

We next investigated what mechanisms could explain the effect of *chs1(G1752a)* mutation on expression noise based on the known function of *CHS1*. *CHS1* encodes a chitin synthase specifically involved in the repair of the chitin septum after cell division in daughter cells (Cabib et al., 1989, 1992), which are the smaller cells resulting from asymmetric division in budding yeast. Detachment between mother and daughter cells involves partial degradation of the septum on the daughter side, which can sometimes damage the cell wall of daughter cells and lead to their lysis in the absence of repair. *chs1(G1752a)* allele generates an early stop codon that leads to a truncated protein lacking several residues and domains that are essential for *chs1* function (Chen et al., 2023); therefore, it is likely to be a null allele of *chs1*. Flow cytometry data showed an enrichment of cells with a smaller size for the *chs1(G1752a)* site-directed mutant relative to the reference strain (**Figure 5d**), consistent with an accumulation of daughter cells with damaged or lysed cell wall in the mutant. When splitting wild-type and *chs1(G1752a)* samples in different intervals of cell size, we observed that the median expression level of *P_TDH3_-YFP* did not depend on cell size (**Figure 5e**). Therefore, the increased expression noise of *chs1(G1752a)* strain is not explained by a difference of median expression level between damaged daughter cells and larger cells in this mutant. Instead, we observed a large increase of extrinsic noise mostly in the smallest cells of the *chs1(G1752a)* mutant: larger *chs1(G1752a)* cells and wild-type cells of all sizes showed much lower extrinsic noise (**Figure 5f**). This observation is consistent with an increased extrinsic noise in damaged daughter cells that accumulate in the *chs1(G1752a)* mutant.

To further show that the effect of *chs1(G1752a)* mutation on extrinsic noise was due to impaired chitin septum repair, we measured expression noise of wild-type and mutant cells grown in presence of glucosamine in the culture medium. Supplementing the culture medium with glucosamine is known to promote chitin synthesis via the activity of CHS3p (Bulik *et al*., 2003), preventing damage to the chitin septum in daughter cells. Therefore, any consequence of *chs1(G1752a)* mutation caused by impaired chitin septum repair is expected to be suppressed in presence of glucosamine. We indeed observed that the extrinsic noise of the smallest *chs1(G1752a)* mutant cells decreased with increasing doses of glucosamine in the culture medium (**Figure 5g**). At 15 mM glucosamine, there was almost no difference of extrinsic noise and of cell size between wild-type and *chs1(G1752a)* cells. When considering all cell sizes (**Figure 5 – figure supplement 3c**), the increase of extrinsic noise in *chs1(G1752a)* cells relative to wild-type cells varied from 136% in absence of glucosamine (*t*-test, *P* = 3.35 x 10^-7^) to only 12% in presence of 15 mM glucosamine in the culture medium (*t*-test, *P* = 5.8 x 10^-3^). We also observed a gradual decrease of median cell size with increasing dose of glucosamine, a pattern that was more pronounced for wild-type cells than for *chs1(G1752a)* cells (**Figure 5 – figure supplement 3a**). This is consistent with the previous finding that yeast cells grown in presence of glucosamine retain less water, for unknown reasons (Bulik *et al*., 2003). Glucosamine-induced reduction in cell size is not sufficient to explain the effect of glucosamine on extrinsic noise, because glucosamine treatment reduced cell size mostly for wild-type cells and it reduced extrinsic noise mostly for *chs1(G1752a)* cells (**Figure 5 – figure supplement 3**). Importantly, the doses of glucosamine used did not significantly impact cell growth rates during the first phase of growth after glucosamine was added to the medium (**Figure 5 – figure supplement 4**). Altogether, these observations strongly suggest that *chs1(G1752a)* mutation increases extrinsic noise of *P_TDH3_* activity exclusively in daughter cells with damaged chitin septum, a phenomenon that is suppressed in presence of glucosamine. The mechanism linking chitin septum repair to extrinsic noise of gene expression remains to be characterized.

### Density-dependent effect of a missense mutation in *MSH1* gene on intrinsic and extrinsic noise

Mutation *m3* is a missense mutation located in the coding sequence of the *MSH1* gene (*G1262a* substitution; **Table 3**). As mentioned above, we observed considerable variation of *P_TDH3_-YFP* median expression and expression noise in the EMS mutant carrying mutation *m3* (YPW2143) between independent experiments. In the initial mutagenesis screen (**Figure 1a**), mutant YPW2143 showed a modest decrease of median expression (−0.9%) and a large increase of expression noise (+72.8%) relative to the reference strain. However, in a subsequent experiment (**Figure 2d**, **Figure 2 – figure supplement 1c**), the same mutant showed a substantial increase of median expression (+6.5%) and a much lower increase of expression noise (+6.3%). Since these two experiments were performed in different laboratories, the observed phenotypic variation could be caused by uncontrolled variation in experimental conditions. We hypothesized that the initial density of cell cultures – a parameter difficult to control exactly when handling a large number of samples in parallel – might influence the effect of *msh1(G1262a)* mutation on *P_TDH3_-YFP* expression. To test this hypothesis, we initiated cell cultures from four different dilutions of a preculture and incubated these cultures for the same duration (20 hours) at 30°C before quantifying absorbance as a proxy for final cell density and expression using flow cytometry. This experimental procedure was applied to haploid strains expressing both *P_TDH3_-YFP* and *P_TDH3_-RFP* reporters with either the wild type allele or the *G1234a* allele of *MSH1*. We found a strong association between the initial dilution of cell cultures and the effects of *msh1(G1234a)* mutation on expression noise of either *P_TDH3_-YFP* (**Figure 6a**) or *P_TDH3_-RFP* (**Figure 6b**). Expression noise increased with initial cell density both for the reference strain and for the *msh1(G1262a)* mutant, but this trend was stronger for the *msh1(G1262a)* mutant, resulting in higher expression noise for the mutant relative to the reference strain at the highest initial density tested (*i.e.* 16:500 dilution). At the lowest initial cell density (*i.e.* 1:500 dilution), we observed a negligible increase of expression noise for *P_TDH3_-YFP* (+4.2%; *t*-test, adjusted *P* = 0.41) and *P_TDH3_-RFP* (+6.4%; *t*-test, adjusted *P* = 0.035) in the *msh1(G1262a)* mutant relative to the reference strain (**Figure 6 – figure supplement 2**). In contrast, at the highest initial cell density (*i.e.* 16:500 dilution), expression noise was increased by 45.7% for *P_TDH3_-YFP* (*t*-test, adjusted *P* = 3.0 x 10^-4^) and by 34% for *P_TDH3_-RFP* (*t*-test, adjusted *P* = 9.9 x 10^-4^) in *msh1(G1262a)* mutant cells (**Figure 6 – figure supplement 2**). We also observed a complex relationship between initial cell density and median expression of *P_TDH3_-YFP* and *P_TDH3_-RFP* (**figure 6 – figure supplement 2**): median expression remained constant among the different initial densities tested for the reference strain; however, for *msh1(G1262a)* mutant cells, median expression changed significantly depending on initial cell density and this variation was not the same for *P_TDH3_-YFP* and *P_TDH3_-RFP* median expression. The fact that initial cell density altered the effect of *msh1* mutation on expression noise and median expression may explain the variability we previously observed between independent experiments. However, variation of initial cell density is not necessarily the direct cause of expression variation. A more direct cause could be a variation of final cell densities after growth of cultures initiated at different cell densities. This seems unlikely, though, as we observed significant differences of intrinsic (*t*-test, adjusted *P* = 1.98 x 10^-3^) and extrinsic (*t*-test, adjusted *P* = 4.42 x 10^-2^) noise for *msh1* mutant cells diluted at 2:500 and at 4:500 (**Figure 6 – figure supplement 2**) despite no significant difference of final cell densities (**Figure 6 – figure supplement 2b**; *t*-test, adjusted *P* = 0.24). Cultures initiated at different cell densities are expected to reach different phases of growth after 20 h of incubation at 30°C, which could explain why we observed density-dependent effects of *msh1(G1262a)* mutation on median expression and expression noise.

We found that *msh1(G1262a)* mutation impacted both extrinsic and intrinsic noise (**Figure 6c**), but only at the highest initial cell density tested (*i.e.* 16:500 pre-culture dilution) and the effect on intrinsic noise was greater that the effect on extrinsic noise (**Figure 6 – figure supplement 2**). No significant difference of intrinsic noise (*t*-test, adjusted *P* = 0.69) and extrinsic noise (*t*-test, adjusted *P* = 0.16) was observed between *msh1(G1262a)* mutant cells and wild-type cells for a 1:500 pre-culture dilution, while a 244% increased intrinsic noise (*t*-test, adjusted *P* = 2.8 x 10^-3^) and a 41% increased extrinsic noise (*t*-test, adjusted *P* = 0.058) was observed in mutant relative to wild-type cells for a 16:500 pre-culture dilution. The effect of *msh1* mutation on intrinsic noise is visible at the single cell level via the lower correlation observed between YFP and RFP expression levels of mutant cells relative to wild-type cells (**Figure 6 – figure supplement 1**).

*MSH1* encodes a DNA-binding protein of the mitochondria involved in the repair of mitochondrial DNA. Disruption of this gene is known to cause a high frequency of respiratory-deficient cells (*ρ-* petite cells), which may generate cell-to-cell heterogeneity and could provide a simple explanation for the increased expression noise observed in the *msh1(G1262a)* mutant. We compared the frequencies of *ρ-* cells in wild-type and *msh1(G1262a)* mutant strains after growth in a rich medium (YPD) containing a fermentable carbon source (see Methods) and using two different pre-culture dilutions (1:500 and 8:500). Both *ρ+* and *ρ-* cells can grow in this medium. As expected, we observed a much lower frequency of *ρ-* cells in the wild-type strain (1.3% and 0.3% *ρ-* cells for pre-culture dilutions of 1:500 and 8:500; **Figure 6d**) than in the *msh1(G1262a)* mutant strain (94.5% and 88.6% *ρ-* cells for pre-culture dilutions of 1:500 and 8:500; **Figure 6d**). We isolated a *ρ-* colony from the *msh1(G1262a)* mutant strain and noticed, as expected, that it produced 100% of *ρ-* cells after growth in YPD medium. We reasoned that if the impact of *msh1(G1262a)* mutation on expression noise was explained by the co-existence of *ρ+* and *ρ-* cells in the mutant strain, then expression noise should be reduced in the *msh1(G1262a)* strain with 100% of *ρ-* cells, and its noise should no longer depend on initial cell density. However, fixing the frequency of *ρ-* cells to 100% did not reduce the effect of *msh1(G1262a)* mutation on intrinsic and extrinsic noise (**Figure 6d**; **Figure 6 - figure supplement 3**). In addition, the expression noise of *msh1(G1262a)* strain was higher in cultures initiated at high cell densities than in cultures initiated at low cell density regardless of the frequency of *ρ-* cells in the population (**Figure 6d**; **Figure 6 - figure supplement 3**). These observations indicate that the effect of *msh1(G1262a)* mutation on expression noise driven by the *TDH3* promoter is not directly explained by cell-to-cell heterogeneity in respiratory capacity caused by the mutation.

### Effects of *yme2*, *chs1* and *msh1* mutations on expression noise are not specific to the *TDH3* promoter

So far, we showed that *yme2(G1234a)*, *chs1(G1752a)* and *msh1(G1262a)* mutations altered in *trans* the expression noise of two reporter genes regulated by the *TDH3* promoter. However, *trans*-acting mutations may potentially impact the transcriptional activity of other promoters that share common regulatory mechanisms with the *TDH3* promoter. To determine whether the three mutations specifically altered transcriptional noise of the *TDH3* promoter or not, we quantified their effects on expression noise of a reporter gene placed under transcriptional control of each of three other promoters (**Figure 7a**). We chose the promoters of three genes (*FBA1*, *TEF1* and *ACT1*) that were found to be expressed at high levels in rich medium in previous studies (Newman et al., 2006; Vande Zande et al., 2022), so that we could accurately quantify expression noise using a fluorescent reporter. In addition, these promoters were reported to be regulated by diverse mechanisms that were more or less similar to regulatory mechanisms of the *TDH3* promoter: *P_TDH3_*, *P_FBA1_* and *P_ACT1_* display a canonical TATA box, which is not the case for *P_TEF1_* (Rhee & Pugh, 2012); and the SAGA coactivator complex plays a dominant role in *P_TDH3_* and *P_FBA1_* activity, while transcription from *P_TEF1_* and *P_ACT1_* is dominated by activity of TFIID (Huisinga & Pugh, 2004).

**Figure 7.**
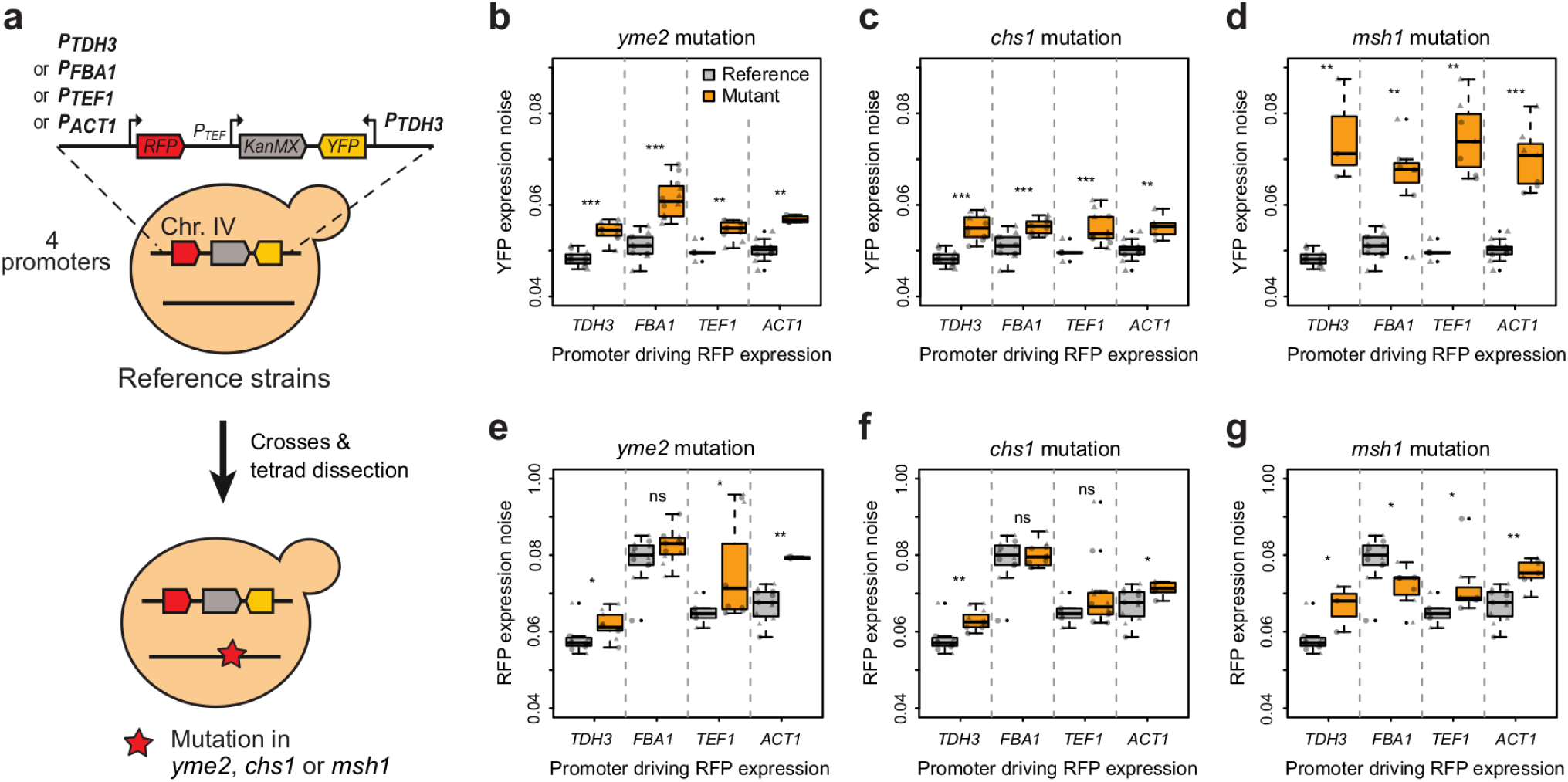
Pleiotropic effects of the three mapped mutations on expression noise measured from different gene promoters. **(a)** Fluorescence levels of single cells were measured in strains expressing *RFP* under control of either *TDH3*, *FBA1*, *TEF1* or *ACT1* yeast promoters. *YFP* was expressed under control of the *TDH3* promoter in all strains. Each strain contained either the wild-type or mutant allele of *yme2*, *chs1* and *msh1* genes. **(b-c)** YFP expression noise and **(e-g)** RFP expression noise of genotypes with different promoters driving RFP expression and reference (gray) or mutant (orange) alleles of **(b,e)** *yme2*, **(c,f)** *chs1* and **(d,g)** *msh1*. Thick lines, boxes and whiskers represent the median, interquartile range (IQR) and 1.5 IQR of expression noise measured among 4 to 12 samples. Dots represent the expression noise of individual samples measured from 2686 to 4606 cells. Data collected for independent clones with the same genotype are represented on the left and on the right of each box. Circles and triangles correspond to samples assayed from two different 96-well plates (no significant plate effect was detected). Results of Mann-Whitney-Wilcoxon tests performed to compare expression noise between wild-type and mutant alleles are represented above each pair of genotypes, with *p*-values adjusted using Benjamini-Hochberg correction (ns: 0.05 ≤ *p*; *: 0.01 ≤ *p* < 0.05; **: 0.001 ≤ *p* < 0.01; ***: *p* < 0.001).

**Figure 7 – figure supplement 1.**
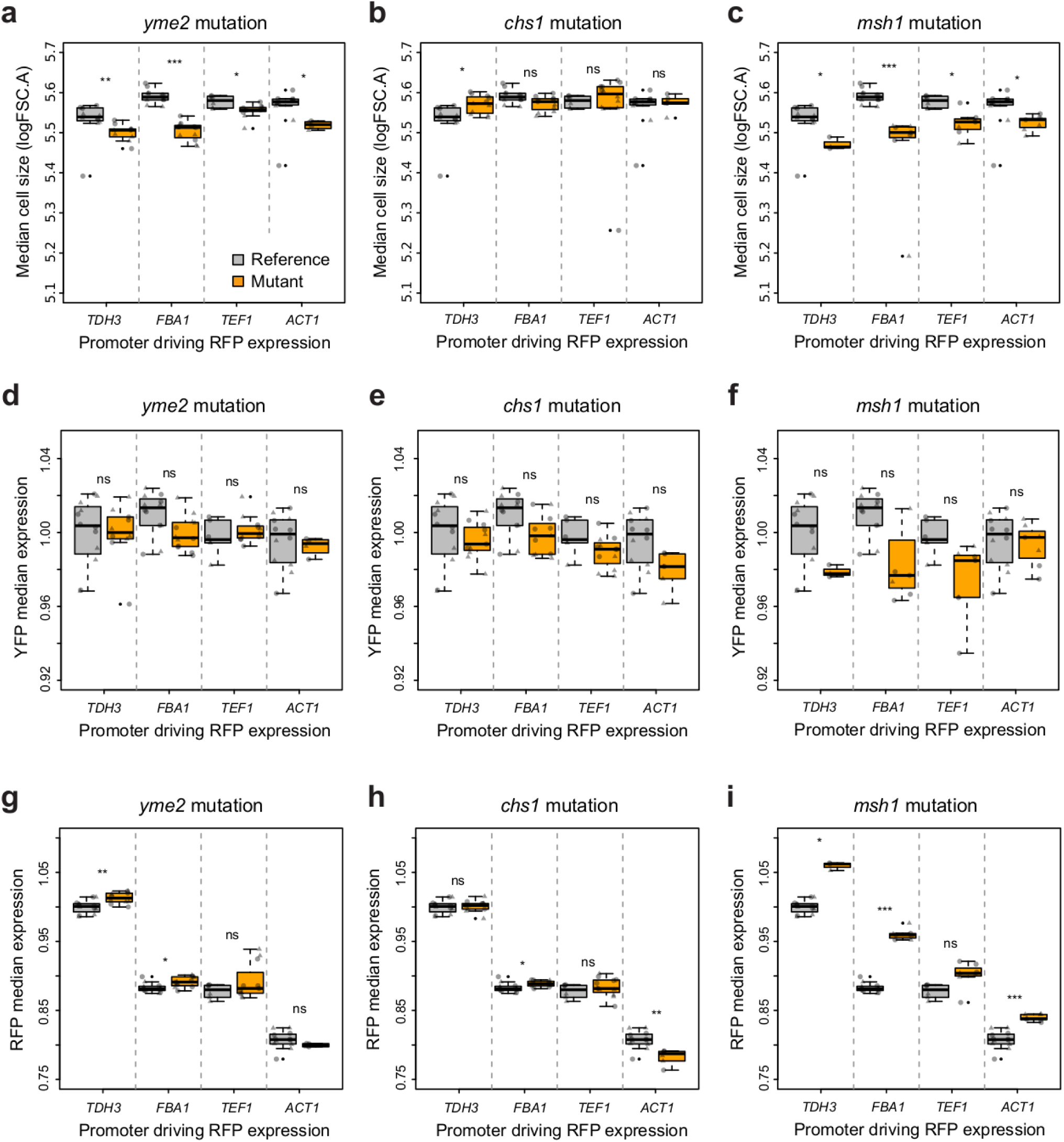
Median cell size and fluorescence of strains expressing RFP under control of four different promoters. (a-c) Median cell size (logarithm of forward scatter), **(d-f)** median YFP expression and **(g-i)** median RFP expression noise of genotypes with different promoters driving RFP expression and reference (gray) or mutant (orange) alleles of (a,d,g) *yme2*, (b,e,h) *chs1* and (c,f,i) *msh1*. Thick lines, boxes and whiskers represent the median, interquartile range (IQR) and 1.5 IQR of expression noise measured among 4 to 12 samples. Dots represent phenotypes of individual samples measured from 2686 to 4606 cells. Data collected for independent clones with the same genotype are represented on the left and on the right of each box. Circles and triangles correspond to samples assayed from two different 96-well plates (no significant plate effect was detected). Results of Mann-Whitney-Wilcoxon tests performed to compare phenotypes between wild-type and mutant alleles are represented above each pair of genotypes, with *p*-values adjusted using Benjamini-Hochberg correction (ns: 0.05 ≤ *p*; *: 0.01 ≤ *p* < 0.05; **: 0.001 ≤ *p* < 0.01; ***: *p* < 0.001).

**Figure 7 – figure supplement 1.**
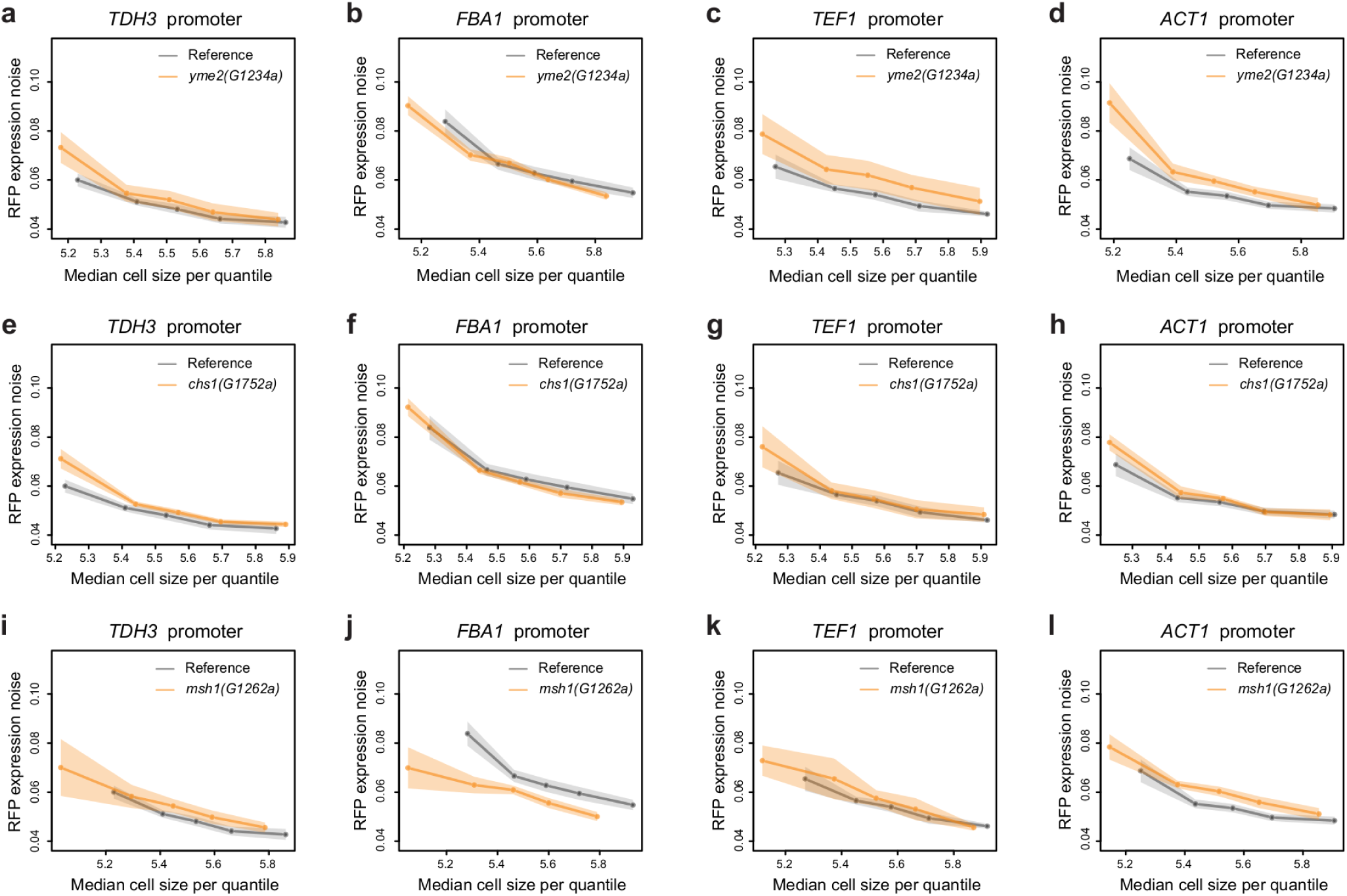
Impact of mutations on the relationship between cell size and RFP expression noise from four different promoters. RFP expression noise is represented for five groups of cells with different cell sizes for strains with wild-type (gray) and mutant (orange) alleles of (a-d) *yme2*, (e-h) *chs1* and (i-l) *msh1*. Cell groups correspond to five fractions of the population of cells delimited by 5-quantiles of cell sizes (same number of cells in each group but different cell sizes). Dots indicate the median cell size (x-axis) and RFP expression noise (y-axis) of each cell group (722 to 870 cells per group in 4 to 12 replicate samples). Shaded areas represent 95% confidence intervals.

We inserted either *P_FBA1_-RFP*, *P_TEF1_-RFP* or *P_ACT1_-RFP* reporter genes in the genome of a strain that contained the *P_TDH3_-YFP* reporter at the *ho* locus, following the same approach as previously used to insert *P_TDH3_-RFP*. We introduced each mutation in genomes harboring these different reporter genes using crosses and tetrad dissections (see Methods). We then quantified cell size, YFP fluorescence and RFP fluorescence for thousand of cells of each strain using flow cytometry. In these strains, median and noise of YFP expression (**Figure 7b-d**, **Figure 7 – figure supplement 1**) were consistent with previous measures of *P_TDH3_-YFP* expression reported above. However, different promoters led to different levels of RFP expression, with 11.7%, 12.2% and 19.3% reductions of median expression for *P_FBA1_*, *P_TEF1_*and *P_ACT1_* relative to *P_TDH3_* (**Figure 7 – figure supplement 1**). *yme2(G1234a)* and *chs1(G1752a)* mutations had very weak effects on median expression for all promoters, while *msh1(G1262a)* mutation increased RFP median expression to a similar extent for the four promoters (**Figure 7 – figure supplement 1**). In addition, different promoters led to different levels of RFP expression noise, with a higher noise driven by the *FBA1* promoter than by *TDH3*, *TEF1* and *ACT1* promoters (**Figure 3e-g**). Most importantly, *P_TDH3_* was not the only promoter for which expression noise was affected by *yme2*, *chs1* and *msh1* mutations. *yme2(G1234a)* mutation significantly increased expression noise relative to the reference allele for *P_TDH3_*, *P_TEF1_* and *P_ACT1_*, but not for *P_FBA1_* (**Figure 7e**). *chs1(G1752a)* mutation significantly increased expression noise for *P_TDH3_* and *P_ACT1_*, but not for *P_FBA1_* and *P_TEF1_* (**Figure 7f**). Finally, while *msh1(G1262a)* mutation increased expression noise for *P_TDH3_*, *P_TEF1_* and *P_ACT1_*, it was found to decrease expression noise when RFP was expressed under control of *P_FBA1_* (**Figure 7g**). Overall, these results show that *trans-*acting mutations can affect expression noise of genes regulated by different promoters, even though the magnitude and direction of effect may not be the same for all promoters.

## DISCUSSION

In this study, we used random mutagenesis followed by genetic mapping to identify three *trans-*acting mutations that increased cell-to-cell variability of expression of a yeast reporter gene. This unbiased approach made it possible to discover any mutation that could possibly alter the expression noise of the focal gene. Indeed, all three mutations identified here were located in the coding sequence of genes that were not known regulators of *TDH3* promoter activity and were therefore not expected to regulate *P_TDH3_-YFP* expression noise. Importantly, these three mutations had very little impact on the median expression of *P_TDH3_-YFP*. Their action on expression noise can therefore not be explained by the well-known scaling of expression noise with expression mean (Newman et al., 2006). The three mutations targeted proteins with different biochemical functions involved in different – but not necessarily unrelated – cellular processes. *chs1(G1752a)* non-sense mutation increased extrinsic noise exclusively in the smallest cells of yeast populations, which was likely caused by damaged cell walls of daughter cells improperly repaired in the absence of chitin synthase I activity. The second mutation targeted the inner mitochondrial membrane protein Yme2 and increased intrinsic noise of *P_TDH3_*expression. The third mutation, targeting the Msh1 ATPase involved in mitochondrial DNA repair, also increased *P_TDH3_* intrinsic noise and this effect varied with the growth phase of cells. Intriguingly, although *yme2(G1234a)* and *msh1(G1262a)* mutations impacted genes with distinct functions, both genes are important for the maintenance of mitochondrial genome integrity, leaving open the possibility that these mutations modulate intrinsic expression noise via a common mechanism. After describing the potential limits of our screen-based approach, we discuss this possibility, as well as the main implications of these results, below.

We decided to identify genetic modulators of expression noise by performing a random mutagenesis screen rather than testing candidate genes because it allowed the discovery of unknown mechanisms without ascertainment biases. Although EMS mutagenesis mostly induces G:C to A:T substitutions, which could slightly bias the screen towards GC-rich genes and coding sequences (relative to regulatory sequences), we do not expect this potential bias to significantly impact our conclusions. Indeed, *YME2*, *CHS1* and *MSH1* coding sequences are not particularly enriched in G or C nucleotides (GC content is 39.7%, 36.8% and 37.0% in these three genes relative to 39.6% among all ORFs) and, in the yeast genome, GC content is not much lower in intergenic sequences (34.4%) than in ORFs (39.6%). We note that bulk-segregant analysis (BSA-seq), which is generally very powerful to map causal mutations in yeast (Duveau et al., 2014; Ehrenreich et al., 2010; Magwene et al., 2011), was only successful here for two of the five mutants that we analyzed (we identified the third causal mutation by another approach based on tetrads). Several possibilities may explain the limited power of BSA-seq in the present study. First, the statistical power to map a variant impacting expression noise is expected to be lower than to the power to map a variant impacting mean expression (Sarkar et al., 2019). Second, based on previous simulations (Duveau et al., 2014), our procedure included three cycles of growth and FACS selection. This is predicted to increase detection power, but only for mutations that do not impair fitness during the growth phase of each cycle. It is therefore possible that some mutations causing elevated expression noise escaped BSA-seq detection because they also impacted growth. Another possibility is that the effect of these mutations may be particularly sensitive to microenvironmental conditions, which could have differed between the characterization of the mutants and the conditions of BSA-seq selection. In the present mapping study where the genetic diversity between the two parental strains was low, tetrad analysis proved to be more powerful than BSA-seq to map mutations altering gene expression noise.

Few studies previously characterized *trans-*acting genetic variants impacting expression noise. Besides pioneering work showing the effect of chromatin remodelers on noise using gene deletions (Raser & O’Shea, 2004), expression-noise modulators were also identified as natural polymorphisms in genes involved in cell-sensing and responses to environmental variation. The first examples discovered were the *MUP1* and *ERC1* genes, which regulate the import of extracellular methionine in yeast cells, and where natural polymorphisms affected noise in expression of the methionine-sensitive reporter *P_MET17_-GFP* (Fehrmann et al., 2013). Another example was later identified in humans, where a disease-associated variant in the *PDE4D* phosphodiesterase - controlling cAMP signalling - was associated with noise in expression of the HLA-DR receptor at the surface of CD8+ T cells and of the CD38 receptor at the surface of CD4+CD45RA+ naive T cells (Lu et al., 2016). Therefore, we could have expected our screen to reveal mutations perturbing cell sensing, but this was not the case. Our screen also did not reveal mutations in transcription factors known to regulate *TDH3* promoter activity, or in the TATA box binding protein (TBP), which were obvious candidates because mutations in transcription factor binding sites and in the TATA box of the *TDH3* promoter are known to sometimes alter expression noise (Duveau et al., 2018; Hornung et al., 2012; Loell et al., 2022; Metzger et al., 2015). However, even though such mutations shift expression noise outside of the global mean-noise relationship observed among genes (Newman et al., 2006), they always impact both mean expression and noise. Here, we focused on mutations that only changed expression noise, which may explain why we did not find mutations in known transcriptional regulators of *TDH3*. In addition, the number of mutations we mapped (3) is small in comparison to the size of the mutational target represented by these genes in the genome (∼0.05% of the genome) and it is possible that a larger screen could have revealed such genes. It is also possible that most mutations altering these candidate genes are lethal – TBP and many transcription factors are essential proteins – and thus could not be recovered from our mutagenesis screen. Conversely, it is not necessarily surprising that the classes of genes we identified here have not been previously associated with *trans*-modulation of gene expression noise. Indeed, previous reports focused on natural variants which, unlike the artificial random mutations studied here, had been exposed to the filter of natural selection. The mutations we identified here would probably be counter-selected in nature since they cause dysfunctions in cell wall repair and mitochondrial genome integrity. The classes of genes we identified as *trans*-acting noise modulators therefore illustrate the complementarity between the analysis of natural variants and the analysis of laboratory mutations.

In a previous study, we screened the same collection of EMS mutants to identify 69 *trans*-acting mutations that changed the median expression of *P_TDH3_-YFP* (Duveau et al., 2021). These mutations were located in genes with different functions (iron metabolism, purine synthesis, transcription) from the three genes we identified here, indicating that different cellular processes are involved in the regulation of median expression level and expression noise. However, all of these mutations (those impacting noise and those impacting median expression) were non-synonymous coding mutations outside annotated regulatory sequences. Therefore, when not exposed to natural selection, random mutations in coding sequences are more likely to alter gene expression (either median level or noise) of another gene in *trans* than mutations in regulatory sequences. To draw more general conclusions about the genetic modulation of expression noise variation, more mutations will need to be identified, in different systems, cell types and organisms, and for the regulation of multiple genes.

The identity of two of the three mutations we identified (*yme2(G1234a)* and *msh1(G1262a)*) suggests that the integrity of the mitochondrial genome is needed to limit expression noise of the *P_TDH3_* nuclear gene promoter. Previous studies linked mitochondrial variability among clonal yeast or human cells to the variability of various cellular traits, including proliferation and transcription rates (Dhar et al., 2019; Guantes et al., 2015; Neves et al., 2010). In these past studies, cell-to-cell variability in mitochondrial content or activity acted as an extrinsic source of expression noise: cells inherited different numbers of mitochondria after division, causing correlated variation in the expression of many genes among cells (Johnston et al., 2012). The situation we observe here is different: the *yme2* and *msh1* mutations increased the variability of expression of two reporter genes within the same cell (intrinsic noise), but not their co-variation among cells (extrinsic noise). This is a very surprising observation because it cannot be explained by the classic extrinsic model, where the mutation would generate variable mitochondrial defects among cells that would in turn cause differences of gene expression among these cells. Instead, our results suggest a more complex model: the mutation amplifies stochastic variability within cells without necessarily generating diverse mitochondrial states among cells. In this model of increased intrinsic noise, the same mitochondrial defect in different cells can lead to different expression levels. How may this happen? It is known that intrinsic noise can be altered by changing the frequency and magnitude of transcriptional or translational bursts. A factor decreasing burst frequency while increasing burst size could increase intrinsic noise without altering mean expression (Pedraza & Paulsson, 2008; Tan & van Oudenaarden, 2010; Zenklusen et al., 2008) , as we observed in *yme2(G1234a)* and *msh1(G1262a)* mutants. Thus, we can hypothesize that a signal is activated in response to mitochondrial defects and transmitted to the molecular complexes involved in transcription or translation. A promising candidate for such a signal would be the retrograde response. This pathway is activated in response to mitochondrial dysfunction and enables cells to compensate for the lack of mitochondrial activity via the regulation of nuclear genes (Butow & Avadhani, 2004; Z. Liu & Butow, 2006). The retrograde response was best characterized in yeast and was also described in other eukaryotes, including human cells, which revealed that its mechanisms and functions are partially conserved across species (Jazwinski & Kriete, 2012). We hypothesize that *yme2(G1234a)* and *msh1(G1262a)* mutations may both activate the retrograde response by causing mitochondrial defects, which could then lead to alteration of transcriptional bursting at the *TDH3* promoter, generating intrinsic noise. The known binding site (GGTCAC) of the retrograde transcription factor (Rtg1-Rtg3 heterodimer; Jia et al., 1997; X. Liao & Butow, 1993) is absent in the *TDH3* promoter, suggesting it is not a direct target of the pathway. However, previous findings suggest that the expression level of *CIT2 –* the best-characterized target gene of the retrograde pathway (X. S. Liao et al., 1991) – may impact *TDH3* promoter activity, thereby an indirect effect of the retrograde response is possible. Indeed, we previously showed that mutations in the Cyc8-Tup1 complex, a direct regulator of *CIT2* transcription (Conlan et al., 1999), altered *TDH3* promoter activity (Duveau et al., 2021). Further work is necessary to determine whether activation of the retrograde response can indeed increase intrinsic noise of the *TDH3* promoter, and whether *yme2(G1234a)* and *msh1(G1262a)* mutations act via this mechanism. For instance, one could quantify intrinsic noise of wild-type and mutant cells when negative or positive regulators of the retrograde signaling such as rtg1, rtg2, rtg3 or mks1 are mutated (Garrigós et al., 2024; Jazwinski & Kriete, 2012). Depending on the result, it will then become interesting to investigate whether mitochondrial retrograde signaling has a similar effect on expression noise of other genes and in other species.

One limitation of our fluorescent screening approach is that it could only identify *trans*-acting mutations if they altered expression noise driven by the *TDH3* promoter. However, we showed that *yme2(G1234a)*, *chs1(G1752a)* and *msh1(G1262a)* mutations also altered expression noise driven by other promoters. Therefore, our conclusions regarding the properties of genetic modulators of expression noise may not necessarily be specific to the *TDH3* promoter. In particular, *yme2* and *msh1* mutations had similar impact on expression noise driven by *TDH3*, *TEF1* and *ACT1* promoters, suggesting mitochondrial genome integrity may be a general mechanism controlling expression noise of several genes. We were surprised to observe that *yme2*, *chs1* and *msh1* mutations had different effects on expression noise driven by *TDH3* and *FBA1* promoters. Indeed, these promoters share several properties and both genes are involved in glycolysis, a pathway for which several genes are known to be co-regulated. However, regulation of *FBA1* transcription was shown to involve binding of a specific transcription factor to a specific promoter motif different from other glycolytic genes (Compagno et al., 1991), which may explain why we observed a much higher expression noise in absence of mutation for *P_FBA1_* relative to *P_TDH3_* and why we observed different effects of *yme2*, *chs1* and *msh1* mutations on expression noise between the two promoters.

In conclusion, the present study provides examples of *trans*-acting mutations affecting gene expression noise, revealing an unexpected link between mitochondrial defects and intrinsic variability in expression of a nuclear gene. In the future, identifying more *trans*-acting mutations, and quantifying their pleiotropic effects on expression noise at the transcriptomic scale and on fitness (as done in Vande Zande et al., (2022) for mutations impacting median expression levels) will be essential to understand how random mutations contribute to natural variation of expression noise.

## MATERIALS AND METHODS

### Biological resources

All yeast strains, oligonucleotides and plasmids used in this study are listed in Supplementary File 1. A genealogy of yeast strains with a brief summary of their genotypes can be found in Supplementary File 2. Laboratory stocks were traced using MyLabStocks web application (Chuffart & Yvert, (2014); https://gitbio.ens-lyon.fr/LBMC/yvertlab/mylabstocks).

### Reference strain & EMS mutants

The collection of 1241 mutant strains obtained by ethyl methanesulfonate (EMS) mutagenesis of the reference strain YPW1139 was described in Metzger et al. (2016) and Duveau et al. (2021). YPW1139 was itself obtained from YPW978, a *MATα* haploid strain with S288c genetic background carrying alleles of *RME1* and *TAO3* from SK1 strain that conferred increased sporulation efficiency (Deutschbauer & Davis, 2005) and alleles of *SAL1*, *CAT5* and *MIP1* from RM11 strain that conferred decreased frequency of respiratory deficient (petites) cells (Dimitrov et al., 2009). YPW1139 was obtained by introducing at the *ho* locus a reporter gene that contained the *TDH3* promoter upstream of the *Venus YFP* coding sequence and the *CYC1* terminator along with a *KanMX4* drug resistance gene. As shown in Supplementary Files 1 & 2, all strains constructed in this study were derived from YPW1139.

### Strain constructions

We describe below different procedures that were applied to generate yeast strains used in this study. For each genotype (with few exceptions), we constructed and used in experiments two strains that originated from two independent colonies obtained after the same transformation. Comparing phenotypes of these independent transformants allowed us to distinguish true effects of a targeted genetic change from undesired effects of potential spontaneous secondary mutations.

#### Reference strains with two fluorescent reporters

To be able to distinguish intrinsic and extrinsic sources of expression noise, we constructed strains GY2458 and GY2459 by inserting a *P_TDH3_-RFP*-*T_CYC1_*construct at *ho* locus in the genome of strain YPW1139. The resulting genotype is *ho::P_TDH3_-YFP-T_CYC1_-KanMX4-T_CYC1_-RFP-P_TDH3_*with the two reporter genes in opposite orientation. We used the *mCherry* variant of *RFP*. The genomic insertion was done using a CRISPR/Cas9 approach adapted to yeast (Laughery et al., 2015). First, we used overlap extension PCR to generate a *P_TDH3_-RFP* fragment with homologies to the insertion site at 5’ and 3’ ends. *P_TDH3_* was amplified by PCR from genomic DNA of strain YPW1139 using oligonucleotides 1Q65 and 1S07. *RFP-T_CYC1_* was amplified by PCR from genomic DNA of strain GY2433 using oligonucleotides 1S04 and 1S05. Equimolar amounts of both amplicons were fused by overlap extension PCR, and 70 bp of homology to the insertion site was added by a final PCR with oligonucleotides 1S06 and 1S08. In parallel, we generated plasmid pGY639 by cloning hybridized oligonucleotides 1R98 and 1R99 between BclI and SwaI restriction sites of plasmid pML104 (encoding guide RNA and Cas9). The repair fragment and pGY639 were transformed in YPW1139 using a classic LiAc/polyethylene glycol and heatshock protocol (Gietz & Schiestl, 2007). Transformants were selected on SD -Ura agar plate (synthetic defined medium lacking uracil). Successful transformation was verified by PCR and sequencing of the insert. Two positive transformants were transferred to a YPG plate to counter select petite cells and then transferred to a SD + 5-FOA plate (0.9 g/L 5-Fluoroorotic acid) to counter select cells carrying the pGY639 plasmid. The two strains were stored at - 70°C in 15% glycerol.

The same approach was used to generate strains with different promoters regulating RFP expression (genotypes *ho::P_TDH3_-YFP-T_CYC1_-KanMX4-T_CYC1_-RFP-P_FBA1_, ho::P_TDH3_-YFP-T_CYC1_-KanMX4-T_CYC1_-RFP-P_TEF1_* and *ho::P_TDH3_-YFP-T_CYC1_-KanMX4-T_CYC1_-RFP-P_ACT1_*). We used the same parental strain and CRISPR/Cas9 plasmid as mentioned above, but different oligonucleotides to generate repair fragments for each promoter. Sequences of these oligonucleotides are listed in Supplementary File 1.

#### Single-site mutants

Point mutations identified via genetic mapping were inserted individually in the genome of strain GY2458. *yme2(G1234a)*, *chs1(G1752a)*, *ald4(C-344t)*, *nop19(C-575t)* and *rsn1(G86a)* mutations were inserted using a similar CRISPR/Cas9 approach as described above and in Duveau *et al*. (2021). For each mutation, a specific recognition site of the guide RNA was cloned in plasmid pML104 corresponding to a sequence adjacent to a PAM motif and near the mutation site in the genome. The repair fragment was obtained by hybridization of two complementary 90-bases oligonucleotides containing the target mutation and flanking genomic sequences (Supplementary File 1). *msh1(G1262a)* mutation was obtained using the *delitto perfetto* approach (Stuckey et al., 2011), because a suitable recognition site for the guide RNA could not be found near the mutation site. First, we inserted a cassette containing *Ura3* and *hphMX4* selection markers (amplified by PCR from plasmid pCORE-UH) at the mutation site in *msh1*. Transformants were selected consecutively on SD medium lacking uracil and on YPD medium complemented with 300 mg/L of hygromycin B (Sigma-Aldrich H3274-1G). Correct insertion of the *Ura3-hphMX4* cassette without mutation was verified by PCR and Sanger sequencing. Second, the *Ura3-hphMX4* cassette inserted at *msh1* locus was replaced by a repair fragment that contained *msh1(G1262a)* mutation. This time transformants were selected on SD medium complemented with 0.9 mg/L of 5-FOA. Correct insertion of the mutation was validated by Sanger sequencing. However, resulting strains were unable to grow on a non-fermentable carbone source (petites), a phenotype frequently observed in *msh1* mutant. We restored respiratory efficiency by crossing these strains with YPW1240 (*MAT***a** strain with same genetic background as YPW1139) and by selecting *MATα*; *msh1(G1262a)*; *ho::P_TDH3_-YFP-T_CYC1_-KanMX4-T_CYC1_-RFP-P_TDH3_*segregating cells after meiosis of the diploid. For each single-site mutant, two independent clones were stored at −80C in 15% glycerol.

To insert each mutation in the genomes of strains with different promoters regulating RFP expression, we used an approach based on tetrad dissections because the fraction of positive clones was low when we used the CRISPR/Cas9 approach. First, we crossed strain GY2903 ( *MATα Msh1 ho::P_TDH3_-YFP-T_CYC1_-KanMX4-T_CYC1_-RFP-P_FBA1_*), strain GY2905 (*MATα Msh1 ho::P_TDH3_-YFP-T_CYC1_-KanMX4-T_CYC1_-RFP-P_TEF1_*) and strain GY2909 (*MATα Msh1 ho::P_TDH3_-YFP-T_CYC1_-KanMX4-T_CYC1_-RFP-P_ACT1_*) each with strain GY2941 (*MAT***a** *msh1(G1262a) ho::P_TDH3_-YFP-T_CYC1_-NatMX4*) that was obtained from another cross described below between YPW2143 and YPW1240. Diploids were selected on YPD agar plates containing Nourseothricin (60 mg/L) and G418 (200 mg/L). Then, meiosis was induced by incubating patches of cells for three days in potassium acetate solution (10 g/L KAc). After sporulation, tetrads were digested in 0.5 mg/mL zymolyase 20T (Seikagaku) and 1M sorbitol for 10 minutes at 30°C. 16 tetrad were dissected on YPD agar plates complemented with 1M Sorbitol. After growth, colonies were screened to identify two segregants with the desired genotype from each cross (genotype *MATα msh1(G1262a) ho::P_TDH3_-YFP-T_CYC1_-KanMX4-T_CYC1_-RFP-P_FBA1_*in strains GY2946 and GY2947, genotype *MATα msh1(G1262a) ho::P_TDH3_-YFP-T_CYC1_-KanMX4-T_CYC1_-RFP-P_TEF1_*in strains GY2952 and GY2953, genotype *MATα msh1(G1262a) ho::P_TDH3_-YFP-T_CYC1_-KanMX4-T_CYC1_-RFP-P_ACT1_* in strains GY2955 and GY2956). Mating type was determined by PCR amplification with oligonucleotides 1E95 and 1E96 followed by gel electrophoresis, because amplicon size depended on the mating type. *Msh1* allele was genotyped by Sanger sequencing of a PCR amplicon obtained with oligonucleotides 1T04 and 1T06. Drug resistance markers (genetically linked to reporter genes) were determined based on growth profiles on YPD agar plates containing either Nourseothricin (60 mg/L) or G418 (200 mg/L). We also selected three other segregants (GY2949 with genotype *MAT**a** ho::P_TDH3_-YFP-T_CYC1_-KanMX4-T_CYC1_-RFP-P_FBA1_*, GY2950 with genotype *MAT**a** ho::P_TDH3_-YFP-T_CYC1_-KanMX4-T_CYC1_-RFP-P_TEF1_* and GY2954 with genotype *MAT**a** ho::P_TDH3_-YFP-T_CYC1_-KanMX4-T_CYC1_-RFP-P_ACT1_*) from this cross to be used in the next crosses. Second, we crossed strains GY2534 ( *MATα chs1(G1752a) ho::P_TDH3_-YFP-T_CYC1_-KanMX4-T_CYC1_-RFP-P_TDH3_*) and GY2634 (*MATα yme2(G1234a) ho::P_TDH3_-YFP-T_CYC1_-KanMX4-T_CYC1_-RFP-P_TDH3_*) each with strains GY2949, GY2950 and GY2954obtained from the previous cross. We followed the same tetrad dissection procedure as described above to obtain segregants with desired genotypes (strains GY2958 and GY2959 with genotype *MATα chs1(G1752a) ho::P_TDH3_-YFP-T_CYC1_-KanMX4-T_CYC1_-RFP-P_FBA1_*, strains GY2962 and GY2963 with genotype *MATα chs1(G1752a) ho::P_TDH3_-YFP-T_CYC1_-KanMX4-T_CYC1_-RFP-P_TEF1_*, strains GY2966 and GY2967 with genotype *MATα chs1(G1752a) ho::P_TDH3_-YFP-T_CYC1_-KanMX4-T_CYC1_-RFP-P_ACT1_,* strains GY2960 and GY2961 with genotype *MATα yme2(G1234a) ho::P_TDH3_-YFP-T_CYC1_-KanMX4-T_CYC1_-RFP-P_FBA1_*, strains GY2964 and GY2965 with genotype *MATα yme2(G1234a) ho::P_TDH3_-YFP-T_CYC1_-KanMX4-T_CYC1_-RFP-P_TEF1_*, strains GY2968 and GY2969 with genotype *MATα yme2(G1234a) ho::P_TDH3_-YFP-T_CYC1_-KanMX4-T_CYC1_-RFP-P_ACT1_*). In this cross, drug resistance was not segregating with *RFP* reporter gene, so we used a PCR screening approach to determine which promoter was upstream of *RFP* coding sequence. *Yme2* allele was genotyped by Sanger sequencing of a PCR amplicon obtained with oligonucleotides 1S82 and 1S83. *Chs1* allele was genotyped by Sanger sequencing of a PCR amplicon obtained with oligonucleotides 1T91 and 1T92.

#### YFP-RFP gene fusion

We edited the genome of strains YPW1139 (*MATα*) and YPW1240 (*MAT***a**) using CRISPR/Cas9 to fuse the *RFP* coding sequence at the 3’ end of the *YFP* coding sequence with addition of a 30-bp linker sequence between the two reporters. The repair fragment was obtained by PCR amplification of *RFP* from GY2433 genomic DNA using oligonucleotides 1U75 and 1U76. CRISPR plasmid pGY666 was obtained by cloning hybridized oligonucleotides 1U79 and 1U80 in pML104. Correct insertion was verified by PCR and Sanger sequencing. We then inserted separately mutations *yme2(G1234a)* and *chs1(G1752a)* in the genome of the *MATα* strain that expressed the YFP-RFP fusion protein. The *yme2(G1234a)* single-site mutant was crossed to the *MAT***a** strain with *YFP-RFP* gene fusion. We then performed tetrad dissection after meiosis of the resulting diploid, and selected segregant clones with *MATα; yme2(G1234a)*; *ho::P_TDH3_-YFP-RFP* genotype. *chs1(G1752a)* mutation was introduced using the CRISPR/Cas9 approach described above.

#### YFP-RFP gene fusion with uTEV cleaving site

We also constructed strains with a *P_TDH3_-YFP-RFP* gene fusion at *ho* where coding sequences of YFP and RFP were separated by a short sequence encoding a cleavage site (ENLYFQS polypeptide) for the tobacco etch virus protease uTEV (Kapust et al., 2001). Using CRISPR/Cas9, we first inserted the uTEV cleavage site between YFP and RFP coding sequences in the genome of strains described above carrying a *YFP-RFP* gene fusion and with or without *yme2(G1234a)* mutation. CRISPR plasmid pGY682 was obtained by cloning hybridized oligonucleotides 1W21 and 1W22 in pML104. The repair fragment containing uTEV cleavage site and homology to 3’ end of *YFP* and 5’ end of *RFP* coding sequences was obtained by PCR amplification of oligonucleotides 1W23 and 1W24. Correct insertion was verified by PCR and Sanger sequencing. Second, we inserted a *P_TDH3_-uTEV* construct at *LYS2* locus in these strains using CRISPR/Cas9 gene editing. The CRISPR plasmid targeting a guide RNA at *LYS2* was obtained by cloning hybridized oligonucleotides and in pML104. The repair fragment was obtained by overlap extension PCR. *P_TDH3_*was amplified by PCR from genomic DNA of strain YPW1139 using oligonucleotides 1R33 and 1W26. *uTEV* coding sequence was amplified from plasmid pRK792 (obtained from Addgene) using oligonucleotides 1W27 and 1W28. Equimolar amounts of both amplicons were fused by overlap extension PCR, and 70 bp of homology to the insertion site was added by a final PCR with oligonucleotides 1W29 and 1W30. Transformants were selected as described above for reference strains. Correct insertion was verified by PCR and Sanger sequencing using oligonucleotides 1W31, 1W32 and 1W33.

### Sequencing genomes of EMS mutants

The genome of the parental strain YPW1139 was sequenced in a previous study (Duveau et al., 2021). Here, we sequenced the genomes of EMS mutant strains YPW2088, YPW2162, YPW2143, YPW2210 and YPW2153 to identify all point mutations that were present in each strain. After growth of each strain in 5 mL of YPD medium for 16 hours at 30°C, genomic DNA was extracted from ∼2 x 10^8^ cells per strain using a Wizard genomic DNA purification kit (Promega). Illumina sequencing libraries were then prepared using a DNA Library Prep kit from ActiveMotif. Samples were sequenced on a MiSeq instrument by ProfileXpert, using a MiSeq standard V2 2 x 250 bp sequencing kit. The sequencing depth was ∼40x per genome (15 M paired-end reads). Sequencing data were analyzed with a Nextflow pipeline (Di Tommaso et al., 2017). Sequences corresponding to P5 and P7 Illumina adapters were removed using *cutadapt* (Martin, 2011), then trimmed reads were aligned to S288c reference genome using *bowtie2* (Langmead & Salzberg, 2012). Variant calling was performed using *freebayes*, and false positive calls were filtered out based on mapping quality MQ. We also removed variants shared by strain YPW1139 and mutant strains, as well as mutations found in at least four mutant strains. Structural variant calling was performed by aligning trimmed reads using *bowtie2* with local alignments to S288c reference genome, and using *GRIDSS* (Cameron et al., 2021) with a joint calling. Results were then filtered using an in-house python script, which removed calls that were not passing *GRIDSS* internal filters, that were found on non-chromosomal segments of S288c, or that showed Variant Allele Fraction below 0.1 as defined in *GRIDSS* documentation. The final list of structural variants was manually reviewed and annotated using IGV (Robinson et al., 2011) and SGD (https://www.yeastgenome.org/). The list is available in **Supplementary File 4**. All scripts used for analyzing whole-genome sequencing data can be found at: https://gitbio.ens-lyon.fr/LBMC/yvertlab/mapping-noise.

### Mapping mutations altering expression noise

#### BSA-seq approach

We previously used a BSA-seq approach to identify mutations associated with variation of *P_TDH3_-YFP* median expression (Duveau et al., 2021). Here, we adapted this approach for mapping mutations associated with variation of *P_TDH3_-YFP* expression noise. First, each haploid mutant strain (resistant to G418) was crossed to the mapping strain YPW1240 (resistant to Nourseothricin) on YPG agar plates. Hybrids were selected after replication on YPD agar complemented with 60 mg/L of Nourseothricin and 200 mg/L of G418. Then,meiosis was induced by incubation for 5 days on YPA agar plates (10 g/L Bactopeptone, 10 g/L Bacto Yeast extract, 2% Kac). Random spores were isolated after incubation in 0.3 µg/L zymolyase 100T for 1 hour at 30°C as previously described (Duveau et al., 2021). Briefly, spores and vegetative cells were pelleted by centrifugation in a microcentrifuge tube, resuspend in 100 µL H_2_O and then vigorously vortexed for 2 minutes. At this stage, spores adhered to tube walls, allowing elimination of vegetative cells and undigested tetrad via several washes with 1 mL of sterile water. Spores were then resuspended in 0.02% Triton-X using brief sonication. Next, 10^6^ spores that did not express RFP (*MATα*) were sorted for each sample by FACS (BD FACS Aria II) at the University of Michigan Flow Cytometry Core. RFP fluorescence was quantified using a 561 nm laser for excitation and a 582/15 nm optical filter for emission. Sorted cells were cultivated in 1 mL of YPD for 24 hours at 30°C. We next sorted each sample into three bulks of 10^5^ cells based on cell fluorescence levels using FACS. YFP fluorescence was quantified using a 488 nm laser for excitation and a 530/30 nm optical filter for emission. The low fluorescence bulk corresponded to cells sorted among the 2% cells in the population with the lowest YFP fluorescence. The high fluorescence bulk corresponded to cells sorted among the 2% cells in the population with the highest YFP fluorescence. The mid fluorescence bulk corresponded to cells sorted among the 10% cells in the population with a fluorescence closest to the median YFP fluorescence. When establishing gates for cell sorting, we were particularly careful to keep the median cell size (area of forward scatter) as similar as possible among the three bulks. Sorted cells were then cultivated for 24 hours in 1.5 ml of YPD. Next, we proceeded with a second round of cell sorting based on YFP fluorescence. 10^5^ cells were sorted by FACS for each fluorescence bulk. For each fluorescent bulk, we only sorted cells with YFP levels from the same portion of the population as in the previous round of FACS. After growth for 24 hours in 1.5 ml of YPD, a third round of cell sorting and growth was performed using the same procedure. Next, genomic DNA was extracted from each sample using a Gentra Puregene yeast/Bact kit (Qiagen). DNA libraries were prepared from 1 ng of genomic DNA using a Nextera XT DNA Library Prep kit (Illumina). Deep sequencing of DNA libraries was performed on a HiSeq 4000 instrument (Illumina) at the University of Michigan Sequencing Core Facility, using a 2 x 150 bp sequencing kit. Demultiplexing of sequencing reads were performed using *bcl2fastq2* v2.17. Sequencing data were next analyzed using a Nextflow pipeline on clusters of the PSMN at ENS de Lyon. The first steps of the pipeline were the same as described above for the sequencing of EMS mutants. Variant calling was performed using *freebayes*, which provided the number of reads supporting each allele at each variant site. We then used custom *R* scripts to only retain variants that were identified in the genome of the corresponding EMS mutant for each sample. Finally, likelihood ratio tests (*G*-tests) were used to determine for each variant site whether the proportion of wild-type and mutant alleles was statistically different in the mid fluorescence bulk and in the combined low and high fluorescence bulks. We reasoned that a mutation associated with an increased expression noise should be found at a lower frequency in the mid bulk than in combined data from low and high bulks. We expected to observe the opposite pattern for a mutation associated with a decreased expression noise, and no significant difference among bulks for a mutation without effect on expression noise and not linked to a causative mutation. All scripts use for analyzing BSA-seq data can be found at: https://gitbio.ens-lyon.fr/LBMC/yvertlab/mapping-noise.

#### Tetrad analysis approach

Mutation *msh1(G1262a)* was the best candidate to explain the increased expression noise of mutant YPW2143 after using BSA-seq, but no significant association could be established due to unexpected allele frequencies in one of the three segregant bulks. To statistically test the association between expression noise and *msh1* genotype, we used linkage analysis based on tetrad dissection. First, we induced meiosis in the diploid strain YPW2318 (obtained by crossing YPW2143 mutant with YPW1240 mapping strain) by incubation for 5 days in YPA medium (10 g/L Bactopeptone, 10 g/L Bacto Yeast extract, 2% KAc). After sporulation, tetrads were digested in 0.5 mg/mL zymolyase 20T (Seikagaku) and 1M sorbitol for 10 minutes at 30°C. 32 tetrads were dissected with a platinum needle mounted on a inverted microscope, resulting in the isolation of the 4 spores from each tetrad on YPD agar plates with 1M sorbitol. After 48 h of incubation at 30°C, colonies were replicated with sterile velvets on selective media to determine how three markers segregated among the 4 spores in each tetrad: *NatMX4* was selected on Nourseothricin (60 mg/L), *KanMX4* was selected on G418 (200 mg/L) and *HphMX* was selected on Hygromycin B (200 mg/L). In addition, the segregation of a red fluorescent reporter inserted at the *MAT* locus (*mata2::yEmRFP-HphMX)* was characterized using a fluorescence stereomicroscope (Nikon SMZ800N). 21 tetrads showed a 2:2 segregation of all markers with a co-segregation of red fluorescence and Hygromycin B resistance, as expected. We then quantified *P_TDH3_-YFP* median expression and expression noise for the 80 segregants from 20 of these tetrads using flow cytometry as described below (data were discarded for the last tetrad due to a possible contamination). In parallel, we used Sanger sequencing of PCR amplicons obtain with oligonucleotides 1T05 and 1T06 to determine the genotype at *MSH1* locus in the 80 segregants. Statistical association between *MSH1*genotype and expression noise was determined using a linear model with expression noise treated as a continuous response variable and genotypes at *MSH1*, *MAT* and *NatMX4* treated as categorical explanatory variables.

### Quantification of median expression and expression noise using flow cytometry

Measurements of median expression and expression noise of the *P_TDH3_-YFP* reporter gene in the 1241 EMS mutants and control strains YPW1139 and YPW978 were performed at the University of Michigan as previously described (Duveau et al., 2021; Metzger et al., 2016), using a BD Accuri C6 flow cytometer (BD Biosciences). Subsequent experiments to measure *P_TDH3_-YFP* expression in single-site mutants were performed at ENS Lyon following a similar procedure to prepare samples, but using a MACS Quant VYB flow cytometer (Miltenyi Biotec). For each experiment, yeast strains were revived from −80°C glycerol stocks on YPG plates (20 g/L yeast extract, 10 g/L peptone, 5% glycerol v/v and 20 g/L agar) at least 3 days and no more than 2 weeks before starting the experiment. After 2 days of incubation at 30°C, YPG plates were kept at room temperature. Then, a small amount of cells (∼10^5^ cells) was inoculated in each well of a 96-deep well plate containing 0.5 mL of filter-sterilized YPD medium (20 g/L yeast extract, 10 g/L peptone, 20 g/L dextrose from Fisher bioreagents) and one 3-mm glass bead per well according to a randomized plate design. We observed that changing the brand of reagents used to make YPD medium could lead to inconsistent measures of expression noise. Culture plates were incubated for 24 h at 30°C with 250 rpm orbital shaking. Cells were maintained in suspension both because of the shaking and the presence of a single glass bead in each well. After this first phase of growth, 2 µL of culture from each well (∼ 2 x 10^5^ cells) were transferred to another plate containing 0.5 mL of fresh YPD per well and a second growth phase was carried on by incubating new plates at 30°C for 18 h with 250 rpm orbital shaking. For the experiment where we exposed cells to glucosamine, we added either 0 mM, 1 mM, 5 mM or 15 mM of D-Glucosamine hydrochloride (ThermoFisher) to the YPD medium for the two phases of growth in 96-well plates. For the experiment where the second culture was initiated at different cell densities (Figure 6), we transferred 1, 2, 4 or 16 µL of culture in different wells and we estimated cell density after growth by measuring the optical density at 660 nm of 0.2 mL of each culture using a Sunrise (Tecan) plate reader. Then, for all experiments, 200 µL from each sample were transferred to 96-well plates with 1.2 µm filters (Millipore MSBVS1210). Culture medium was aspirated with vacuum, then cells were washed twice in 200 µL of phosphate-buffer saline (PBS) before fixation in 200 µL of 2% paraformaldehyde (PFA) for 8 minutes. After aspiration, cells were resuspended in 200 µl of PBS containing 0.1 M Glycine for 12 minutes, and washed twice in 200 µL of PBS. A final 1/20 dilution was applied by transferring 5 µL of each sample into 195 µL of PBS in a flat bottom 96-well plate (Falcon). Plates were stored up to 24 hours at 4°C prior to flow cytometry acquisition.

Forward scatter and fluorescence were recorded for 10,000 events per sample with the MACS Quant VYB. A 488-nm laser was used for excitation and a 525/50 optical filter was used for acquisition of YFP signal, while a 561-nm laser was used for excitation and a 615/20 optical filter was used for acquisition of RFP signal. Flow cytometry data were analyzed using custom R scripts ( **Source Code 1**) with a similar approach as in Duveau et al. (2018), except for certain points mentioned below. First, we filtered out artifactual events and doublets (events corresponding to multiple cells) based on values of the area and height of forward scatter (FSC.A and FSC.H) as previously described. Then, we excluded cells that were clear outliers in each fluorescence channel using flowClust package. Raw fluorescence values of cells are strongly correlated with forward scatter (a proxy for cell size). We therefore computed normalized fluorescence values that were independent from cell size, as previously described (Duveau et al., 2018). For each sample, we then computed the median cell size among cells, as well as the median fluorescence and the coefficient of variation (CV) of fluorescence for each fluorescence channel (YFP and RFP) as measures of expression noise. Extrinsic and intrinsic noise were calculated using formula from Fu and Pachter (2016). First, we applied quantile normalization to YFP and RFP fluorescence values of cells (after cell size normalization) using *normalize.quantiles* function from R package *preprocessCore*. Then, intrinsic noise was calculated as:

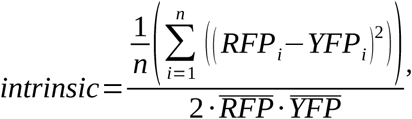

and extrinsic noise was calculated as:

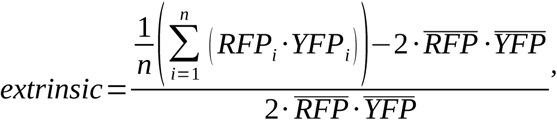

where *n* is the number of cells in a sample, *RFP_i_* and *YFP_i_* are quantile-normalized fluorescence levels of each cell and *RFP* and *YFP* are means of quantile-normalized fluorescence levels among cells. For each sample, median cell size, median fluorescence, CV of fluorescence, intrinsic noise and extrinsic noise were also calculated for five subsets of cells defined based on five quantiles (quintiles) of FSC.A values (area of forward scatter). Statistical comparisons between genotypes or conditions were performed via pairwise *t*-tests using data from replicate samples (samples sharing the same genotype and condition). Benjamini-Hochberg procedure (FDR) was used to adjust *P* values for multiple testing.

### Western blot

The protease from tobacco etch virus (TEV) cleaves the specific site ENLYFQS in proteins (Kapust et al., 2001; Parks et al., 1994), resulting into two residues when the target sequence is inserted in a protein sequence. A western blot on whole-cell protein extract was performed to determine how complete was the cleavage of YFP-ENLYFQS-RFP fusion protein by the classic TEV and improved uTEV protease both in wild-type and *yme2(G1234a)* cells. Control genotypes that expressed either no RFP, RFP alone or YFP-RFP fusion protein were also included (seven strains in total). YFP-RFP and RFP are detected using a primary antibody targeted against mCherry and distinguished based on band sizes after electrophoresis. GAPDH constitutive control were detected with anti-GAPDH antibody for all proteins extract.

Yeast strains were thawed from glycerol stocks kept at –80°C onto YPG agar plates 3 days before experiment. After 2 days of incubation at 30°C, YPG plates were placed at room temperature. The day before the experiment, a small amount of cells (∼10^5^ cells) was inoculated in 4 mL of YPD medium and incubated overnight at 30°C with 250 rpm orbital shaking. 1 mL of cell culture was then transferred to 50 mL of YPD medium and incubated for another 4 hr at 30°C to reach the exponential phase of growth. All cells were pelleted by centrifugation and resuspended in 1 mL of sterile water.

After another centrifugation at 10,000 g for 1 minute, cell pellets were resuspended on ice in 300 µL of fresh lysis buffer (0,1 % w/v sodium deoxycholate, 1 mM EDTA pH8, 50 mM HEPES-KOH pH 7.5, 140 mM NaCl, 1 % w/v Triton W-100 and proteinase inhibitors (Sigma P8215). Reagents were mixed by pipetting up and down. To disrupt cell walls and membranes, we added 500 μL of cold acid-washed glass beads (Sigma G8772) and the tubes were placed on a PreCellys apparatus. Tubes were shaken at 6,800 rpm through 2 cycles, each cycle consisting of 10 s ON, 10 s OFF, 10 s ON. Tubes were placed for 5 min on ice between the two cycles. Lysates were collected in a microtube and stored at −80°C. Protein concentration was measured using the Bio-Rad Protein Assay (#500-0006) from optical density at 595 nm (Bradford) based on linear regression curve made with standards. The samples were diluted in lysis buffer to load 50 µg of proteins and denatured 5 min at 95°C.

Proteins were separated on a 12 % SDS-PAGE gel electrophoresis at 100 V for 2 hours and then transferred to nitrocellulose membrane at 0,2 µM by semi-dry transfer. The membrane was blocked 1 hour at room temperature with 5 % non-fat dry milk in Tris-buffered saline (TBS) containing 0,02 % Tween. The membrane was incubated with primary antibodies against mCherry in TBS with 0,02% Tween and 0,3% milk (1:1000; Invitrogen, PA5-34974) overnight at 4°C. The membrane was washed three times during 10 minutes in TBS with 0,02% Tween and incubated with the secondary antibodies against rabbit in TBS with 0,02% Tween and 0,3% milk (1:10000; Odyssey; 926-32210) for 1 hour at room temperature. The membrane was washed as before and the protein band was visualized by a ChemiDoc MP imaging system (Biorad) with ECL kit (Thermo Scientific) and analyzed using ImageJ. YFP-RFP fusion protein had an expected size of 55 kDa and the RFP residue alone had an expected size of 26 kDa. The membrane was washed one last time and incubated with GAPDH-HRP conjugate antibody in TBS with 0,02% Tween and 0,3% milk [1:2000; AbCam; ab85760] for 1 hour at room temperature. The GAPDH protein bands were revealed as before, with an expected size of 36 kDa.

## SUPPLEMENTARY FILES

**Supplementary File 1.** Yeast strains, oligonuclotides and plasmids used in this study.

**Supplementary File 2.** Genealogy of yeast strains constructed in the study. **Supplementary File 3.** Properties of all point mutations identified in five mutants. **Supplementary File 4.** Structural variants detected with *GRIDSS* analysis.

**Supplementary File 5.** Additional text describing the rationale behind three experimental choices.

## SOURCE CODE

**Source Code 1.** R scripts used to analyze flow cytometry data.

## AUTHORS CONTRIBUTION

FD and PJW conceived the study; FD and LM supervised the work; NM, FD, EP, QB, AD, MP and HDB performed experiments; FD, JKS, AB and QB analyzed data; FD, GY and PJW interpreted results; FD wrote the manuscript; GY reviewed and edited the manuscript.

## ACKNOWLEDGEMENTS

We thank Gérard Triqueneaux for sharing protocols, Taslima Haque for comments on the manuscript, the SFR Biosciences Gerland-Lyon Sud (UAR3444/US8) and University of Michigan Flow Cytometry Core for access to flow cytometers and technical assistance, the University of Michigan Sequencing Facility and ProfileXpert for Illumina sequencing, the Pôle Scientifique de Modélisation Numérique for computing resources. We also thank developers of R, Bioconductor, Ubuntu and Git.

## FUNDING

Agence Nationale de la Recherche – JCJC VORTEX (ANR-20-CE12-0022)

- Fabien Duveau

Fonds de Recherche ENS Lyon – Projet Emergent 2020-2022 (R02/B-R02-FR-S60)

- Fabien Duveau

Horizon Europe, European Research Council Consolidator eGRIDE (101126053_ERC-2023-COG)

- Fabien Duveau

National Institutes of Health (R01GM108826)

- Patricia Wittkopp

National Institutes of Health (R35GM118073)

- Patricia Wittkopp

National Science Foundation (MCB-1929737)

- Patricia Wittkopp

European Molecular Biology Organization – Long-Term Fellowship (1114-2012)

- Fabien Duveau

The funders had no role in study design, data collection and interpretation, or the decision to submit the work for publication.

## DATA AVAILABILITY

Illumina sequencing data have been deposited on the European Nucleotide Archive of the EBI under Study PRJEB104107 (https://www.ebi.ac.uk/ena/browser/home). This includes data for genome sequencing of the five EMS mutant strains described in the study, as well as data for deep genome sequencing of three bulks of segregants sorted based on fluorescence (BSA-seq data) for each of the five mutants crossed to the mapping strain YPW1240.

## Notes

https://gitbio.ens-lyon.fr/LBMC/yvertlab/mapping-noise

### Competing Interest Statement

The authors have declared no competing interest.

### Summary of Updates

The manuscript has been peer-reviewed by two reviewers through the Review Commons platform. We modified the manuscript to address several insightful comments from reviewers. Most importantly, we included results from a new experiment in Figure 7 showing that the effects of yme2, chs1 and msh1 mutations were not specific to the TDH3 promoter. We also reduced the length of the introduction and figure legends, and we moved some parts of the results to Supplementary File 5 to improve the clarity and concision of the manuscript. We included three new authors who contributed significantly to the revision.

