## Supplementary figures and images for "*Trans-*acting mutations reveal non-nuclear modulators of both intrinsic and extrinsic gene expression noise in a eukaryote"

### Supplementary File 2

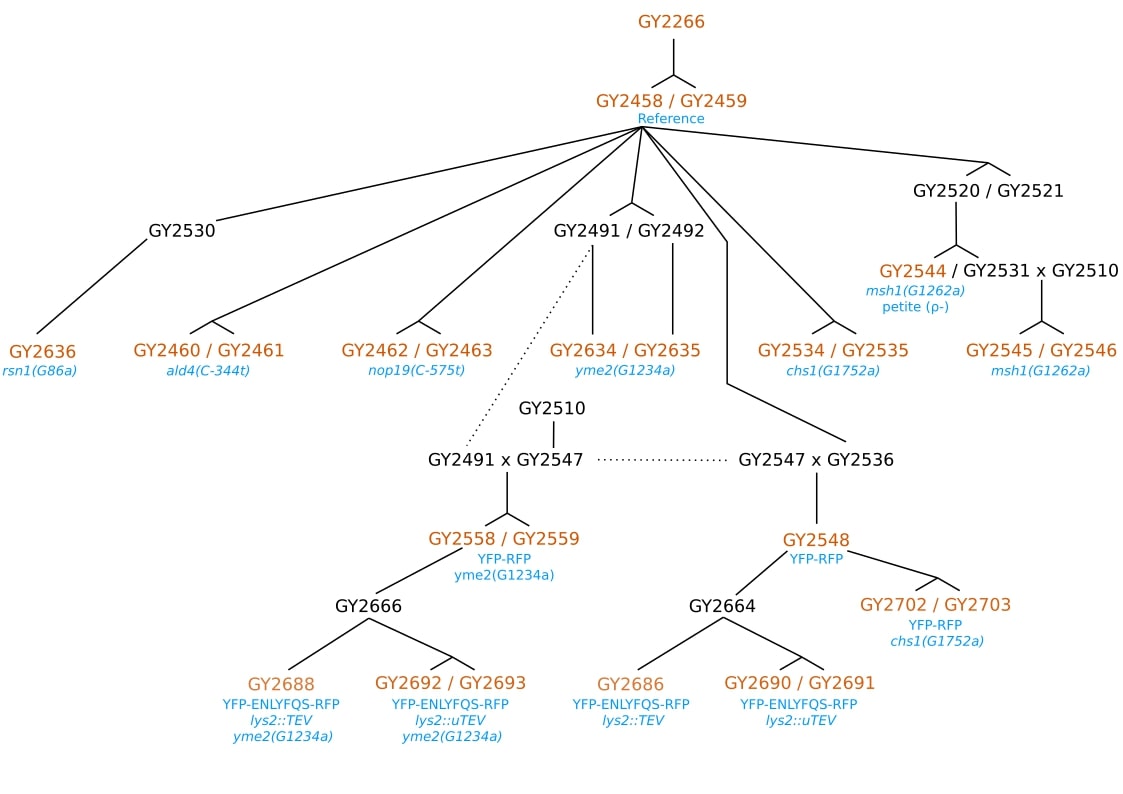
