## Supplementary File 5 for "*Trans-*acting mutations reveal non-nuclear modulators of both intrinsic and extrinsic gene expression noise in a eukaryote"

**Supplementary File 5. Additional text describing the rationale behind three experimental choices.**

### Choice of mutants with altered $P_{TDH3}$-*YFP* expression noise

To identify strains with altered expression noise without change in median expression in our collection of EMS mutants, we first quantified the average and variability of fluorescence levels (see Methods) from thousands of cells for each of the 1241 mutant strains and for the reference strain using flow cytometry (**Figure 1a**). This initial screen revealed a wide variation of expression noise among mutants that was largely – but not completely – independent from variation in median expression. Next, we selected 254 mutants that showed largest changes of expression noise relative to the reference strain and we quantified their fluorescence a second time in an independent assay (**Figure 1 – figure supplement 1**). Among these 254 mutants, we chose five mutants for further analyses that showed significant changes of expression noise relative to the reference strain in both assays (*t*-test, adjusted $P < 0.01$), with no significant change in median expression in the first assay (*t*-test, adjusted $P \geq 0.05$). These five mutants were picked arbitrarily among nine mutants that fulfilled our selection criteria. Three of the selected mutants had an increased level of $P_{TDH3}$-*YFP* expression noise and two mutants had a decreased level of noise (**Figure 1a**, **Table 1**). This procedure ensured that the five selected mutants showed reproducible changes in expression noise that were not explained by variation of median expression levels.

### No effect of *yme2* mutation on intrinsic noise detected from $P_{TDH3}$-*YFP-RFP* fusion

Because the elevated intrinsic noise observed in *yme2(G1234a)* mutant was unexpected, we tested whether it could be explained by an experimental artifact. In particular, the detection of intrinsic noise may not only be caused by biological noise, but also by technical noise in the measurements of YFP and RFP fluorescence using flow cytometry. To exclude this possibility, we fused the coding sequences of *YFP* and *RFP* downstream of the *TDH3* promoter sequence ($P_{TDH3}$-*YFP-RFP*) and inserted the resulting transgene at the *ho* locus in genetic backgrounds carrying either the wild-type *YME2* allele or the *yme2(G1234a)* mutation. We reasoned that expressing a YFP-RFP protein fusion would lead to an equal amount of yellow and red chromophores within a cell (variation can still exist among cells), which would completely prevent the detection of biological intrinsic noise (**Figure 3 – figure supplement 2a**). Therefore, any difference of intrinsic noise detected in this context could be attributed to technical noise affecting the detection of YFP and RFP signals. We observed no significant effect of *yme2(G1234a)* mutation on intrinsic noise detected in strains expressing $P_{TDH3}$*-YFP-RFP* (**Figure 3 – figure supplement 2b**), while the mutation significantly increased levels of extrinsic noise (**Figure 3 – figure supplement 2c**) and total noise (**Figure 3 – figure supplement 2d**). This result confirmed that the impact of *yme2(G1234a)* mutation on intrinsic noise detected with two reporter genes was caused by actual differences in the amount of YFP and RFP molecules within each cell. When YFP and RFP signals were forced to co-vary using a YFP-RFP protein fusion, then *yme2(G1234a)* mutation still increased expression noise but could only be detected as an extrinsic source of noise.

**Experimental strategy to disentangle the contributions of protein partitioning and expression regulation on intrinsic noise detection**

Genetic variation of intrinsic noise detected using a dual reporter system may by explained by two alternative hypotheses (**Figure 4d**). In hypothesis 1, higher intrinsic noise is explained by higher variability of expression of the two fluorescent reporters (yellow and red hexagons in **Figure 4d**), either via altered regulation of transcription or translation. In hypothesis 2, higher detection of intrinsic noise is not caused by altered regulation of expression, but instead by altered partitioning of fluorescent proteins at cell division. If fewer proteins are inherited by daughter cells (because of smaller bud size or other mechanisms), this could lead to higher variability of colors detected by fluorescence in daughter cells and therefore higher intrinsic noise detected. A strategy to test these hypotheses is to express a YFP-RFP fusion protein that can be cleaved by a protease in two separate YFP and RFP fluorescent proteins after translation. With this system, we expect *yme2* mutation to have no detectable impact on intrinsic noise if it acts at the expression level (hypothesis 1) because the expression of the two reporters is forced to covary. However, we expect to detect increased intrinsic noise if *yme2* mutation acts on protein partitioning (hypothesis 2) because cleaved proteins can segregate independently at cell division.
